# Evolutionary tuning of rhodopsin-mediated light protection

**DOI:** 10.64898/2026.05.05.722510

**Authors:** Nadir H. Dbouk, Madeleine Bagshaw, Anamika A. Bose, Audrey M. Arner, Kun Zhao, Neena Praveen, Isabella Bautista, Narmin Musayeva, Liying Xue, Marin McElhinney, Rohan Kasthuri, Alexander G. Bick, Amanda J. Lea, Gianni M. Castiglione

**Author notes:** **Major Classification:** Biological Sciences. **Minor Classification:** Evolution (Primary); Biophysics and Computational Biology (Secondary).

## Abstract

Visual sensitivity introduces the inherent risk of blinding photodamage. Rhodopsin directly mediates this vulnerability, as activation drives excessive signaling via the Meta-II (MII) state, which subsequently releases the toxic, desensitizing agonist all-*trans* retinal (atRAL). In dim-light specialists, slow atRAL release is assumed to reflect extended MII signaling; however, this model ignores that arrestin quenches signaling in milliseconds and that apparent release reflects reversible atRAL rebinding. Here we reveal that extended post-bleach lifetimes— including an atRAL-driven shift toward non-signaling Meta-III (MIII)—evolved independently of intrinsic G-protein signaling to dually protect retinal cells against photochemical toxicity and shield neighboring receptors from agonist noise. Dim-light specialists whose rod-dense retinas experience daytime solar flux, such as megabats, possess an extremely cytoprotective rhodopsin that retains nearly 60% of atRAL at equilibrium by partitioning into MIII—validating long-standing predictions that MIII buffers retinoid toxicity while explaining why bat rhodopsin signals more weakly than faster-releasing human or bovine orthologs. To definitively uncouple sequestration from intrinsic signaling, we engineered an active-state-locked, signaling-null rhodopsin that achieves 100% atRAL sequestration and confers extreme cytoprotection in retinal cells. Mimicking partial bleaches with heterogeneous active and inactive receptor populations, co-incubation reveals that this signaling-null sink acts *in trans* to boost wild-type rhodopsin regeneration and signaling by shielding neighboring dark-state receptors from agonist interference. Lastly, while consensus human rhodopsin is non-protective, alleles restoring cytoprotective sequestration emerged in high-irradiance geographies, conferring a 38% reduced risk of blindness in modern populations. Our findings redefine rhodopsin as a tunable light buffer shaping adaptation and human disease.

**Significance Statement:** Vision requires light-activated rhodopsin to release the reactive chromophore all-trans retinal (atRAL), which can accumulate to toxic levels and drive blinding macular degenerations. Across vertebrate evolution, slow atRAL release in dim-light animals has long been assumed to reflect a prolonged G-protein signaling state. Here we overturn this paradigm, revealing that slow retinal release instead reflects reversible atRAL sequestration into a non-signaling Meta-III state that acts as a “light buffer”. This sequestration dually protects retinal cells against acute atRAL toxicity and enhances visual sensitivity by shielding neighboring receptors from agonist noise. While consensus human rhodopsin lacks this protection, high-capacitance alleles emerged in high-irradiance geographies and reduce blindness risk by 38% in modern populations.

## INTRODUCTION

Vision is predicated on a volatile chemical mixture of oxygen, light, and reactive vitamin A-derived chromophores^1–3^. While this vulnerability is appreciated in the context of ophthalmology^4–6^, we lack a comparative understanding of how evolution may adapt these molecular systems to protect against breakdown risk in different species^3,7,8^. The vertebrate rod visual pigment, rhodopsin (RHO), is the primary source of these chromophores, and thus typically viewed as a liability to be inhibited through clinical intervention^9–11^. First characterized in Franz Boll’s seminal paper as ‘visual red’^12^, RHO has become a quintessential model in vision research, and the study of how ‘Rhodopsin-like’ (class A) G protein–coupled receptors (GPCRs) activate at atomic resolution^13–25^. Photoisomerization of the RHO inverse agonist and chromophore, 11-*cis* retinal (11CR), to the agonist all-*trans* retinal (atRAL), causes RHO activation through several photo-intermediates, culminating in the Meta-II (MII; R*) conformation, the most signaling-active form capable of engaging the G protein transducin (G_t_) and ultimately relaying a neural signal^20,21,26–31^. Mutations in RHO that impair folding, chromophore binding, trafficking, or signaling disrupt this cascade and are well-established causes of retinal diseases such as autosomal dominant retinitis pigmentosa (adRP) and congenital stationary night blindness, among others^32–37^. Defects in the rate of atRAL release by RHO—a process that allows regeneration of dark-state RHO sensitivity^23^— are less well studied.

Beyond its critical regulatory, structural, and dim-light signaling functions, RHO is a central mediator of bright-light damage^38–40^. Photochemical damage is traditionally divided into Classes I and II depending on wavelength, exposure time, and cell type affected^41–44^. While the effects of blue light (Class II) are complex^43,45,46^, Class I photochemical damage induced by white light is mediated by the excessive activation of RHO^41,47^, leading to aberrant phototransduction and retinoid accumulation and subsequent retinal degeneration^41,47,48^. Crucially, in the absence of RHO, extreme photic stress fails to induce photoreceptor apoptosis, demonstrating that the receptor is strictly required for this degenerative process ^38^. Because R* is essential for activating the phototransduction cascade, degeneration was initially attributed to excessive signaling and the resulting metabolic stress^41,47,49^. Yet, photoreceptors still undergo light-induced damage when RHO is present but downstream effectors such as G_t_ are absent, demonstrating that degeneration can proceed independently of canonical cascades^39^. This implicates retinoids (e.g., atRAL) as key mediators of light-induced toxicity^44,48,50–56^. Following RHO activation, released atRAL is partitioned between two fates: enzymatic reduction and clearance, or reversible condensation with phosphatidylethanolamine (PE) to form N-retinyldene-PE (N-ret-PE) ^57–67^. N-ret-PE lipid adducts are flipped into the outer segment by ABCA4, allowing the atRAL to eventually be reduced by retinol dehydrogenases (RDH8/12) and trafficked to the pigment epithelium (RPE)^33,68–70^. However, N-ret-PE adducts can undergo secondary reactions to form irreversible bisretinoids such as A2E and RALdi—the major fluorophores of lipofuscin, a major clinical hallmark of age-related and juvenile macular degenerations^50,59,61,64–68,71–83^.

Aside from these products, free atRAL itself is intrinsically cytotoxic due to its highly reactive aldehyde and conjugated polyene structure^50,61,64,66,84,85^, inducing cellular toxicity via oxidative stress, ferroptosis, and mitochondria-associated cell death^48,52–55,84^, and triggering inflammation, complement activation, and microglia and macrophage infiltration into the subretinal space ^77,86,87^. Critically, ∼60% of atRAL remains free at steady state (i.e. not in lipid-adducts)^88^, and can trigger photoreceptor degeneration when atRAL production^44,53,56^ outpaces clearance by ABCA4 and RDH8/12 ^51,52,56,57,61,64,89,90^. Despite redundant clearance mechanisms, the mass action kinetics of RDHs ensure that free atRAL persists in the photoreceptor at concentrations that scale to ambient light intensity^51,89^. Because N-ret-PE formation outpaces enzymatic reduction, atRAL release from RHO is rate-limiting for visual cycle and dark-state RHO regeneration^23,57,91,92^, making control of initial atRAL bursts critical for photoreceptor survival and performance ^50,51,93^.

Rods and scotopic vision are particularly vulnerable during age-related macular degeneration (AMD), with delayed dark-adaptation serving as a biomarker for early AMD^94,95^. Free atRAL may partly underlie this symptom by causing a postbleaching noise^81,96,97^. Beyond its well-documented cytotoxicity, free atRAL acts as a diffusible agonist that can reassociate with an empty opsin apoprotein that spontaneously sample the active-state conformation, stimulating signaling in the dark^98–100^. As an agonist, atRAL rebinding stabilizes the R* conformation, delaying thermal decay to the dark state conformation required to bind 11-*cis* retinal, the inverse agonist that terminates signaling and regenerates sensitivity ^19,20,100,101^. Consequently, impaired atRAL clearance correlates with delayed dark adaptation and diminished quantum-catch, as seen in Stargardt disease and *ABCA4*-mediated macular degenerations^102–105^. Indeed, the loss and recovery of visual sensitivity following intense illumination is mirrored by initial rise and decline in atRAL levels^106–110^. Notably, diminished sensitivity is likely not caused by depletion of 11CR alone; it is likely also influenced by atRAL-implicated dark noise, as observed in aged mice^102,111^. Specifically, the *opsin/atRAL* complex forms a partially active signaling state that continuously stimulates transducin independent of light^81,96,100,109^. *In vivo*, this aberrant signaling generates post-bleach transduction noise—effectively creating an ‘equivalent background light’ that desensitizes the rod cell^109,111–113^. This atRAL-induced dark noise is orders of magnitude higher than that induced by either empty RHO, or all-*trans* retin**ol** bound RHO^96–99,112,114^. Although single cell recordings with exogenous excess atRAL have not consistently demonstrated desensitization^115^, these experiments do not replicate the localized production and transient accumulation of atRAL *in vivo*. Under physiological conditions, even small, spatially restricted pools of atRAL may bias opsin activity. Maintaining absolute scotopic sensitivity may therefore require suppressing local, transient atRAL accumulation.

The dual threat posed by atRAL to photoreceptors (cytotoxicity and desensitization) likely underlies the evolution of buffering mechanisms like N-ret-PE and the non-signaling photoisomerase, “retinal G protein-coupled receptor (RGR)”, which act as atRAL sinks that neutralizes the reactive aldehyde of atRAL^89,116,117^. This reflects a broader paradigm in which dedicated receptors function primarily as ligand scavengers (e.g., atypical chemokine receptors which can signal to arrestin but not G proteins^118,119^ or TNF/TRAIL decoy receptors^120^). While rhodopsin is a canonical signaling GPCR, we reasoned that it may also occupy an unprecedented and central role in modulating atRAL-induced toxicity and noise, as R* determines the production rate of free atRAL through hydrolysis and release (Fig. 1A)^88^. Such agonist buffering by a signaling-competent receptor itself would contrast to the specialized sequestration of non-signaling decoy GPCRs ^118,119^—allowing R* to function as a state-dependent capacitor that continuously shapes its own ligand microenvironment. Importantly, R* does not release atRAL irreversibly, but rather exists in a dynamic equilibrium in which atRAL can rebind in proportion to its local concentration, thereby quenching the toxic atRAL aldehyde through Schiff base formation (Fig. 1A-B) ^92,101,121^. Thus, R* has been proposed to ‘moonlight’ as a state-dependent sink for its own agonist^51,122,123^. Through reversible sequestration, R* may therefore buffer photic atRAL “loads”, smoothing the flux between production and clearance pathways, and is thus biophysically analogous to an electrical capacitor, which perform the dual function of absorbing transient surges and filtering high-frequency noise ^124–126^. We therefore propose that the atRAL-buffering capacity of R* similarly limits acute atRAL accumulation and the consequent retinal degeneration as seen in mice while also suppressing retinoid-driven delays to dark-adaptation.

**Figure 1:**
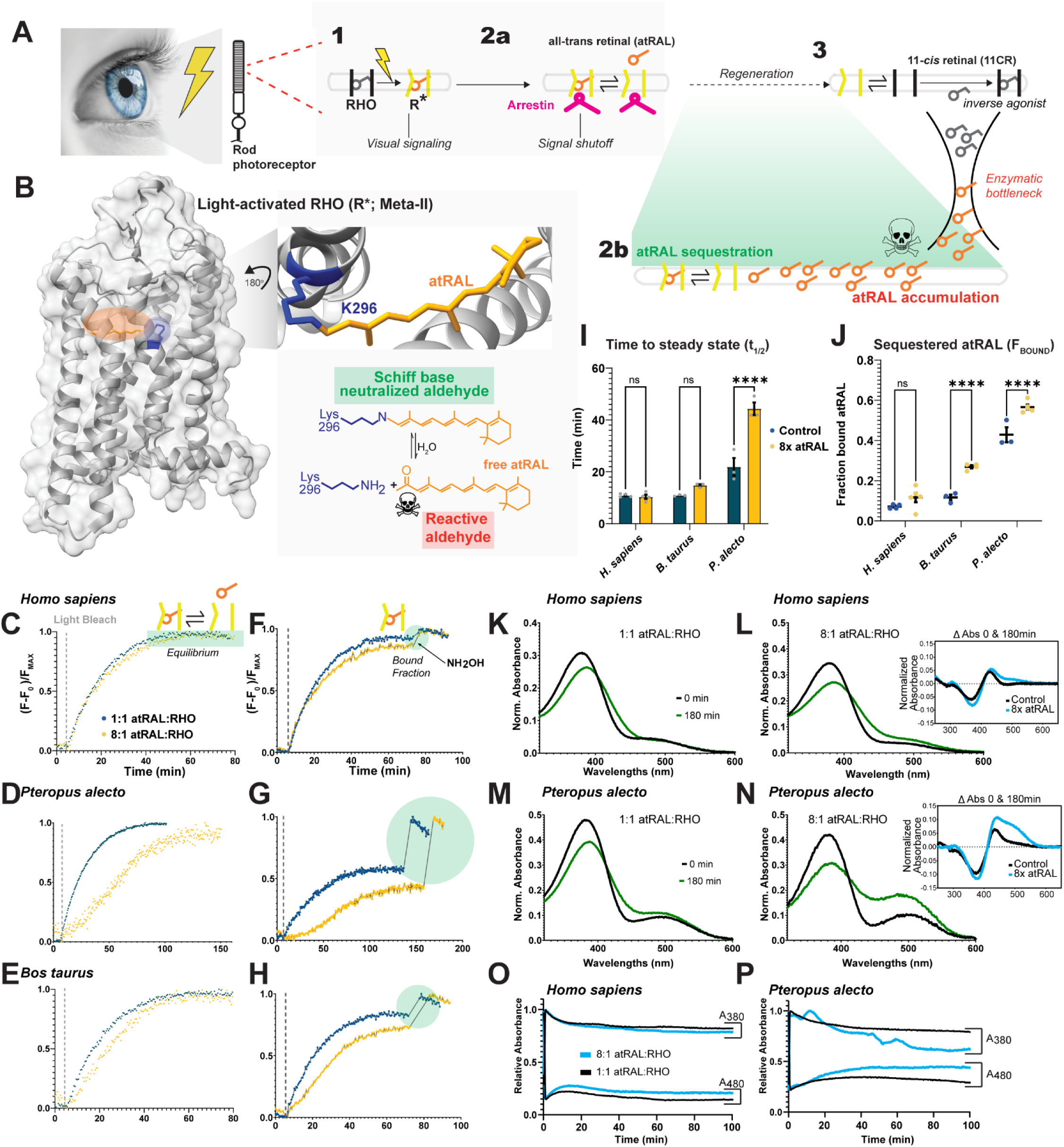
Light-activated rhodopsin acts as a tunable sink for the toxic visual byproduct all-*trans*-retinal (atRAL). **(A)** Following photoactivation, the 11-*cis*-retinal (11CR) chromophore of rhodopsin (RHO) isomerizes to all-*trans*-retinal (atRAL) triggering formation of light-activated rhodopsin (R*/Meta II). R* signaling is quenched by arrestin binding, followed by atRAL release to enable regeneration to the dark state. This process is bottlenecked by visual cycle processing of atRAL to 11CR, leading to transient toxicity due to atRAL accumulation. R* buffers free atRAL in dynamic equilibrium before conformationally decaying to the dark state and binding the inverse agonist 11CR. **(B)** Structural representation of the R* (Meta-II;MII) conformation of RHO highlighting the reversible Schiff base at Lysine 296, which neutralizes the reactive aldehyde of atRAL. **(C–E)** Retinal release kinetics after light activation monitored via intrinsic tryptophan fluorescence for recombinantly expressed *Homo sapiens* (*C*), *Pteropus alecto* (*D*), and *Bos taurus* (*E*) rhodopsin under basal stoichiometric conditions (1:1 atRAL:RHO; blue) and in the presence of 2 µM exogenous atRAL (8:1 atRAL:RHO; yellow). (**F–H**) Quantification of atRAL sequestered at equilibrium for *H. sapiens* (*F*), *P. alecto* (*G*), and *B. taurus* (*H*). Following the initial fluorescence plateau, hydroxylamine (NH_2_OH) is added to cleave the Schiff base, allowing quantification of the bound atRAL fraction (*F_bound_*). (I) Calculated time to equilibrium (*t*_1/2_) across species under control (1:1) and 8x exogenous atRAL (8:1) conditions. (J) Calculated fraction of sequestered atRAL (*F_bound_*) at steady state. (**K–N**) Post-bleach UV-Vis absorbance spectra recorded immediately after light activation (0 min; black) and at equilibrium (180 min; green) for *H. sapiens* (*K, L*) and *P. alecto* (*M, N*) RHO at 1:1 (*K, M*) and 8:1 (*L, N*) molar ratios of atRAL:RHO, showing conversion of the 380 nm MII peak into the ∼480 nm Meta-III (MIII) photointermediate in *P. alecto*. Insets in (*L*) and (*N*) display difference spectra (ΔAbs between 180 and 0 min post-bleach) under control (1:1; black) and 8x atRAL (8:1; blue) conditions. (**O–P**) Continuous time-course tracking of relative absorbance at 380 nm (*A*_380_, MII) and 480 nm (*A*_480_, MIII) over 100 minutes following photoactivation for *H. sapiens* (*O*) and *P. alecto* (*P*) RHO at 1:1 (black) and 8:1 (blue) atRAL:RHO ratios.

If R* functions as a photoprotective, noise-scrubbing capacitor, its biophysical properties should be tuned to an organism’s visual ecology if critical for visual performance. Indeed, functional variation in RHO shapes fitness-associated traits across the tree of life^127–131^. Beyond spectral tuning of absorbance^129,132–134^, a major axis of selection is the stability of R*, typically defined by Meta-II (MII) lifetime and atRAL release kinetics ^101,135^. Reduced R* stability accelerates pigment regeneration by promoting rapid return to the dark state and 11CR binding, a requirement reflected in the low stability and fast turnover of cone opsins^23,136–139^. Conversely, nocturnally and dim-light adapted species tend to have extended R* lifetimes^136,137,140–142^. Although visual arrestin binding rapidly quenches rhodopsin signaling on the millisecond timescale, the MII conformation persists substantially longer, establishing a local buffering window that limits atRAL escape before full regeneration of the dark state (Fig. 1A)^91,92,143–145^. By retaining atRAL near its site of release, we hypothesize that this reduces both aberrant signaling and cytotoxic exposure.

We previously proposed R* stability and atRAL sequestration capacities are influenced by an organism’s photodamage risk, balancing rhodopsin abundance and environmental light flux against its need for visual recovery^122,123^. Here we demonstrate that R* sequestration of atRAL protects against cellular toxicity while simultaneously promoting visual sensitivity through mechanisms not previously described. We find that ecological photic load drives the evolution of a capacitor phenotype that enhances atRAL sequestration across mammals—by favoring a non-signaling Meta-III (MIII) sequestration state—and within ancestrally high-irradiance human populations. Guided by this principle, we engineered a signaling-null synthetic RHO locked in the active conformation with atRAL capacitance exceeding natural variants. Using this construct to biophysically decouple ligand sequestration from G-protein signaling, we demonstrate that pure active-state capacitance confers extreme cytoprotection, shields neighboring dark-state receptors from agonist interference during regeneration, and amplifies the signaling of wild-type R* via mass action. Together, these results redefine the prototypical class-A GPCR, rhodopsin, as a tunable light buffer that dynamically regulates its own toxic agonist.

## RESULTS

### R* sequestration of toxic atRAL varies across visual mammals

To determine whether visual species dynamically tune the equilibrium between free atRAL and receptor-bound retinal, we monitored retinal release rates across three animal RHOs through intrinsic tryptophan fluorescence, which is quenched when atRAL occupies the binding pocket^135^. We utilized the visual nocturnal black fruit bat^146,147^, *Pteropus alecto*, which possesses a RHO with one of the highest reported half-lives for an extant species (∼40 min)^142^. As a comparison, we utilized human and bovine RHO, which are diurnal model systems that decay more rapidly (∼10 and ∼12 min, respectively)^101,148^. Through absorbance spectroscopy, we show the purified pigments displayed the expected wavelength of maximum absorbance (λ_max_) and were activated by light as previously described (Fig. S1;^123^). Using fluorescence spectroscopy, the pigments also displayed light-dependent retinal release as expected (Fig. 1C-E). Relative to previous results, our data reflects similar trends with human and bovine RHO *t*_1/2_ at 10.5 and 11.5 min respectively (Fig. 1I, Table S1), while *P. alecto* RHO decayed at a *t*_1/2_ of 22 min (Fig. 1D), notably shorter than the reported 40 min^142^. Given identical sequences and consistent λ_max_ values, the observed differences are most likely due to the absence of an additional 1D4 peptide at the C-terminus used in previous studies^149^; we omitted the additional sequence since *P. alecto* naturally has the epitope, and it still retains a >2-fold slower t_1/2_, therefore serving our purposes as a highly stable RHO comparison. We next monitored how retinal release rates were affected across species in the presence of 2 µM exogenous atRAL (8:1 molar ratio of atRAL:RHO), which has been shown to push the equilibrium away from atRAL release and towards rebinding by RHO^101^. While no change was observed in *H. sapiens* RHO retinal release rates, we observed a trend of slower decay in *B. taurus* RHO and a significant delay in *P. alecto* RHO, with the latter doubling from 22 minutes at 0μM atRAL to 44.3 minutes at 2μM atRAL (p ≤ 0.05; Fig 1C-E, I). We also observed a marked kinetic difference of *P. alecto* RHO where it follows a sigmoidal curve under 2µM atRAL rather than the expected exponential rise (Fig. 1D;^101^). Moreover, there appears to be a similar, but less pronounced effect amongst *B. taurus* retinal release kinetics (Fig. 1E) suggesting that effect is not isolated to *P. alecto* but rather this atRAL-induced kinetic property is exacerbated in chiropteran RHO. This indicates there may be more than one kinetic rate constant governing atRAL release not previously elucidated, reflecting equilibrium effects by exogenous atRAL, perhaps indicating the presence of metarhodopsin I or III^91^. Altogether, these data demonstrate that across species, increases to atRAL and R* stability push the equilibrium away from atRAL release and towards rebinding by RHO.

To gain additional insights, we next quantified the proportion of atRAL sequestered by RHO at the steady state (i.e. the plateau in fluorescence), which involves the continual release and rebinding of atRAL at equilibrium ^101,122,150^. To measure the quantity of atRAL bound at steady state, hydroxylamine (HA) was administered to cleave the covalent Schiff base linkage, displacing any re-bound atRAL from the internal binding pocket and forming a retinal oxime incapable of rebinding RHO^19,101^. The resulting increase in fluorescence enables quantification of active state (R*; presumably MII) bound atRAL using the fraction bound (*F_bound_*) ^101,122,150^. We found that in human RHO, only ∼7% of the population is bound to atRAL at steady state (Fig. 1F, J). Moreover, the addition of 2μM exogenous atRAL has no significant effect on this, consistent with the lack of effect of atRAL on human R* retinal release rate (Fig. 1I). This suggests the active conformation of human RHO with its low binding affinity for atRAL (Table S1) is not significantly stabilized by the equilibrium effects of exogenous atRAL ligand^19,101^. By contrast, Bovine RHO shows an increase from approximately 12% to 25% *F_bound_* in the presence of 2µM atRAL (Fig. 1H, J), consistent with previous results and the slowed retinal release rate under 2 µM atRAL (Fig. 1I)^101^. Critically, in the RHO of the visual chiropteran bat, *Pteropus alecto*, we observed that ∼40% of the RHO population is bound to atRAL at steady state, which increases to a massive 60% population when exogenous atRAL shifts the equilibrium towards rebinding (Fig. 1G, J; Table S1), consistent with its significantly slower retinal release (Fig. 1I). Using these *F_bound_* metrics, we calculated apparent dissociation constants (*K_d_*) across species (Table S1; methods). *Pteropus alecto* displays a >16-fold higher binding affinity for atRAL than that of human RHO. These results demonstrate that nature actively tunes atRAL binding affinity across species.

### Extended retinal release reflects an atRAL-sensitive subpopulation of Meta-III

Although slow retinal release rates are classically attributed to prolonged MII stability ^101,135,141^, our data indicate that retinal release in *Pteropus alecto* RHO under 8:1 molar excess atRAL is governed by biphasic, sigmoidal kinetics (Fig. 1D, G). Because the MII photointermediate (λ <u>≈</u> 380nm) can thermally decay via syn/anti isomerization of its protonated Schiff base into the non-signaling Meta-III (MIII) state (λ <u>≈</u> 465-480nm) prior to chromophore hydrolysis and release^145,151–154^, we hypothesized that extended retinal retention in *P. alecto* reflects a substantial subpopulation of MIII whose equilibrium with MII is shifted by local atRAL concentration. To test this directly, we used UV-Vis absorbance spectroscopy to monitor post-bleach transitions between MII (380 nm) and MIII (480 nm) over a 180-minute time course in *P. alecto* RHO compared to *H. sapiens* RHO, which did not exhibit sigmoidal release kinetics under excess atRAL (Fig. 1C, F). In *H. sapiens* RHO under basal stoichiometric conditions (1:1 atRAL:RHO), the post-bleach 380 nm peak decreased modestly over 180 minutes with no detectable increase at 480 nm (Fig. 1K), and addition of an 8:1 molar excess of atRAL:RHO induced only a minor absorbance shoulder in the 480 nm region (Fig. 1L, inset, blue curve). By contrast, *P. alecto* RHO formed a distinct ∼480 nm peak by 180 minutes post-bleach even at a basal 1:1 atRAL:RHO molar ratio, consistent with spontaneous conversion from MII to an MIII subpopulation (Fig. 1M). Challenging *P. alecto* RHO with an 8:1 molar excess of atRAL:RHO dramatically expanded this ∼480 nm MIII band over 180 minutes, establishing a prominent bimodal equilibrium between MII (380 nm) and MIII (∼480 nm) (Fig. 1N). Difference spectra (ΔAbs between 0 and 180 minutes post-bleach) confirmed a transition across an isosbestic point at ∼415 nm, where the loss of 380 nm absorbance was accompanied by a broad 465–515 nm MIII gain that was markedly amplified by excess atRAL in *P. alecto* compared to *H. sapiens* (Fig. 1L, N, insets). Tracking continuous *A*_380_ and *A*_480_ kinetics over 100 minutes corroborated these endpoint spectra and matched the timescales of our fluorescence release assays (Fig. 1O–P). Whereas *H. sapiens* RHO maintained high *A*_380_ (∼0.80) and low *A*_480_ (∼0.15–0.21) plateaus with minimal sensitivity to excess atRAL (Fig. 1O), *P. alecto* RHO displayed a ∼2-fold higher basal rise in *A*_480_ (∼0.35) that reached steady state within ∼30–40 minutes (Fig. 1P), mirroring its 22-minute basal fluorescence *t*_1/2_ (Fig. 1D, I). Under 8:1 atRAL:RHO challenge, *P. alecto* exhibited a pronounced, reciprocal divergence in which *A*_480_ climbed to a ∼0.45 steady-state plateau over the first 45 minutes while *A*_380_ declined to ∼0.62 (Fig. 1P)— matching the 44.3-minute sigmoidal lag phase observed in tryptophan fluorescence dequenching (Fig. 1D, G, I). Importantly, this post-bleach rise at 480 nm cannot represent Meta-I (which precedes MII on the millisecond timescale^28,155^) or dark-state regeneration (λ = 500.5 nm; Fig. S1, Table S1), as no 11CR is present in the purified preparation, its formation is actively promoted by exogenous atRAL (which inhibits dark-state folding), and the entire bound retinoid pool at steady state remains susceptible to hydroxylamine cleavage (Fig. 1G), unlike the hydroxylamine-resistant dark state^24,101,135^. Because MIII retains covalently bound atRAL in the binding pocket and thereby continues to quench Trp265 fluorescence, these results demonstrate that slow apparent retinal release rates reflect an atRAL-stabilized subpopulation of MIII in addition to MII, which is evolutionarily expanded in *P. alecto*.

### RHO retinal release rates do not predict signaling duration or intensity

Because MIII is a non-signaling photointermediate that does not activate transducin (G_t_)^91,145^, our finding that *P. alecto* RHO partitions a substantial fraction of bound atRAL into MIII predicts that slow retinal release rates^140,141^ do not necessarily confer enhanced G-protein signaling-consistent with *in vivo* evidence^23^. To affirm the in vivo evidence, we measured the intensity and duration of human, bovine and bat R* signaling to a G protein transducin chimera (G_t/s_) in a live cell assay^156^. This *in vitro* luminescent Glosensor assay (Promega) relies on a Gt/Gs chimeric protein, allowing RHO activation to generate a luminescent cAMP signal that reflects Gt signaling activity^156^. As a negative control, we transfected HEK293T cells with the Gt-Gs chimera, glosensor plasmids, and an empty vector with no RHO (EV), and observed no change in luminescence following light exposure (Fig. 2A). When human, bovine, or bat RHO were co-transfected with glosensor and Gt/Gs, we observed significant increases in luminescence following light exposure. Critically, the bovine and human R*-mediated Gt activation was significantly greater than that produced by *P. alecto* RHO. Since the Gt interaction domain that binds RHO is 100% conserved between these species, this provides direct evidence that *P. alecto* R* is a significantly less effective signaling receptor compared to the far less stable bovine and human R*. This aligns with previous work demonstrating that retinal release rates can be an unreliable proxy for the lifetime of the signaling-competent MII conformation ^101^. Together with our spectroscopic evidence (Fig. 1K–P), this paradox—where *P. alecto* RHO retains nearly 60% of atRAL at equilibrium yet signals more weakly than human or bovine orthologs—confirms that extended retinal release half-lives in dim-light specialists reflect a high-affinity equilibrium biased toward a non-signaling MIII ‘sequestration vault’ rather than prolonged phototransduction.

**Figure 2:**
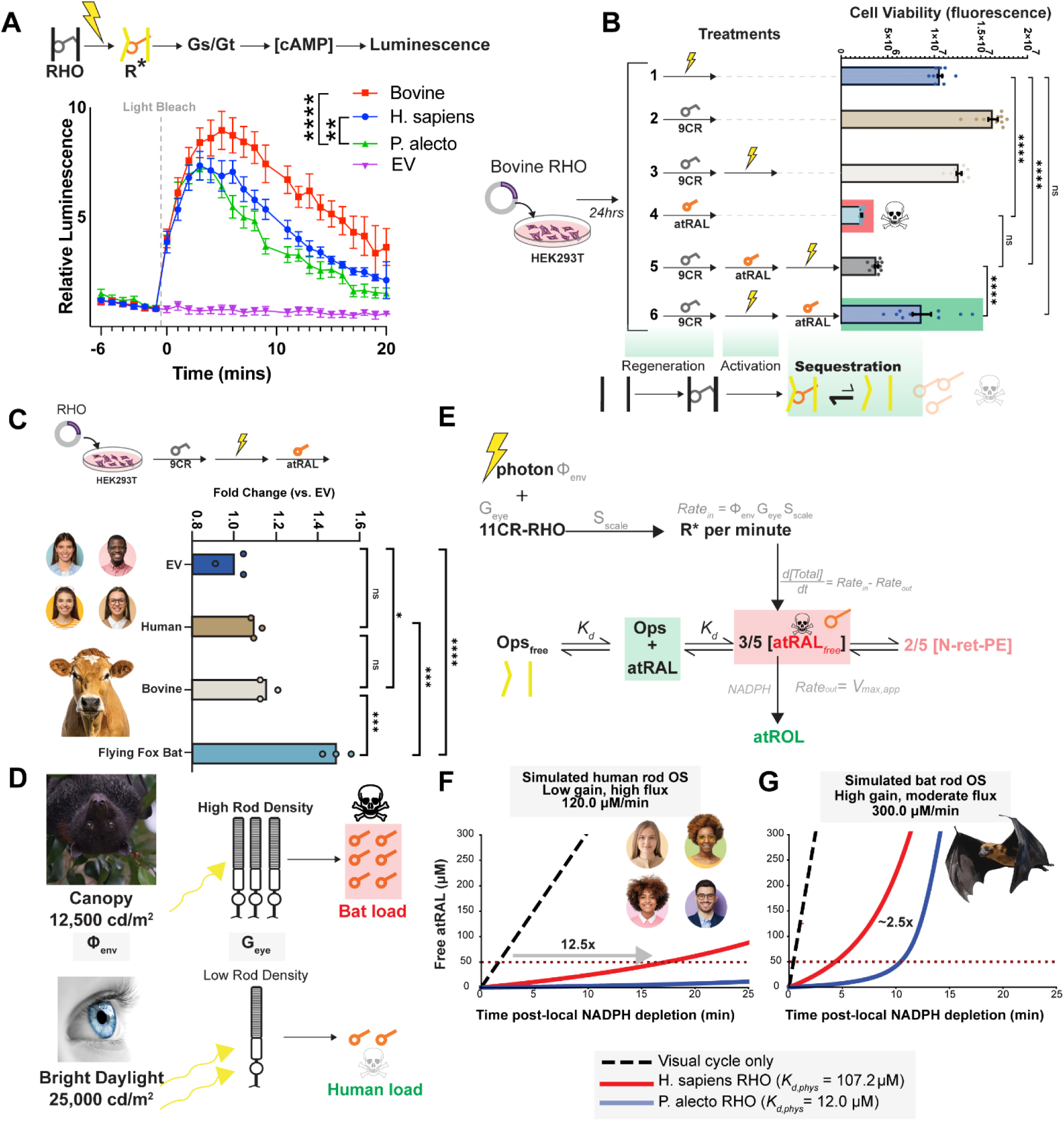
R* sequestration of atRAL is cytoprotective and calibrated to species-specific ecological photic loads. **(A)** Glosensor assay monitoring activation of G_t_/G_s_ by R* within transfected HEK293T cells after light bleach. Luminescence normalized to protein expression levels (Fig. S1). **(B)** Cell viability under various treatment paradigms modulating RHO conformation. Protection against 50 µM atRAL toxicity strictly requires prior light-activation of RHO (Treatment 6), the conformation competent to bind atRAL. **(C)** Cytoprotection against atRAL stress across heterologously expressed species variants. Cell titer blue, normalized to protein expression data (Fig. S1). **(D)** Ecological photic loads. Humans experience bright daylight (∼25,000 cd/m^2^) with a low-gain retina, while the canopy-roosting *P. alecto* bat experiences moderate flux (∼12,500 cd/m^2^) but possesses a highly amplified, high-gain retinal architecture, generating a massive predicted atRAL load (methods). **(E)** Schematic of the hybrid deterministic kinetic model simulating atRAL generation (inflow), RDH-mediated clearance, and dynamic buffering by the RHO capacitor and PE-lipid adduct condensation (methods). **(F-G)** Simulated free atRAL accumulation following local NADPH depletion under a 120 µM/min human load (F) and a 300 µM/min bat load (G). The presence of human or *P. alecto* RHO drastically extends the time to toxicity (TTT) compared to a visual-cycle-only model, with *P. alecto* RHO exhibiting specialized atRAL capacitance to manage its extreme physiological atRAL load.

### Rhodopsin-mediated sequestration of atRAL during photobleaching is cytoprotective

Why have dim-light specialists such as the flying fox bat (*P. alecto*) evolved long retinal release rates if they do not necessarily reflect long R* signaling times? Critically, delayed atRAL release from R* delays regeneration with 11-*cis* retinal and thereby the *in vivo* regeneration of visual sensitivity after light bleaches^23,91,157,158^. We hypothesized that strong atRAL binding affinity and the consequently high atRAL sequestration at steady state, may afford cytoprotection against atRAL toxicity. In vivo, this would afford a buffer, or capacitor, that sequesters high free atRAL loads, providing a temporary sink for when enzymatic clearance is overwhelmed by atRAL production rate. Such a sink already exists as N-ret-PE, although these products eventually accumulate as bisretinoids—clinical markers of macular degeneration.

To test this, we utilized HEK293T cells—which can express heterologous RHO at high levels— to determine an atRAL toxicity range. Based on previous work in other cell lines^159–164^ we measured untransfected HEK293T cell viability after a 4hr challenge with various exogenous atRAL concentrations (0–225 μM) using the CellTiter-Blue (CTB) assay. A clear dose-dependent reduction in viability was observed starting at 25–50 μM atRAL compared to the untreated control (Fig. S2). These findings are consistent with previous reports showing decreased cell viability and elevated cellular stress at these atRAL concentrations that are consistent with physiological estimates (15–50 μM^51,159–164^). To investigate the putative cytoprotective role of RHO atRAL sequestration, we utilized 50 µM atRAL as a mild challenge that stays within the estimated physiological range. To bind atRAL, RHO must be in the R* active conformation^19^. We therefore hypothesized that any putative RHO-mediated protection against atRAL toxicity would require R*. To test this, we utilized a series of treatments following transient transfection of HEK293T cells to heterologously express bovine RHO (Fig. 2B). To establish a baseline, cells were either illuminated with light for 30 seconds or regenerated with 9-*cis* retinal (9CR; a cost-effective analog of 11CR) (Fig. 2B). Under dim red light, 9CR improved cell viability relative to light exposed cells, likely due to its known inverse agonist and chaperone function promoting proper protein folding and reducing proteotoxic stress^165,166^. When 9CR addition was followed by bright light exposure (treatment 3; Fig. 2B), this beneficial function was eliminated, suggesting the light-induced release of atRAL chromophore reduced cell viability. Unlike 9CR (treatment 2), addition of equimolar concentration of atRAL (treatment 4) resulted in a severe loss of viability, consistent with dark-state RHO being able to bind 9CR but unable to bind atRAL due to conformational selectively^19^. To demonstrate this, we modulated the RHO conformation using combinations of regeneration with 9CR, followed by either atRAL challenge or light exposure. We observed no difference in toxicity when atRAL challenge was followed by light, consistent with a dark-state RHO being unable to sequester atRAL in the interim before light activation (treatment 5; Fig. 2B). Consistent with our hypothesis, atRAL toxicity was significantly rescued when prior to atRAL challenge, RHO was light activated (treatment 6; Fig. 2B). This provides evidence that RHO has a photoprotective role against atRAL toxicity that depends on photoactivation—the exact photic condition in which RHO (i.e. R*) is competent to bind atRAL^19^. This mirrors the *in vivo* sequence of events in healthy mammalian retinae where atRAL levels only increase after a light bleach ^52,69^. This indicates that the persistence of the RHO active state, R*, for timescales that exceed signal shutoff, may reflect the evolution of RHO as a cytoprotective atRAL buffer or capacitor.

If true, then RHO orthologs with higher atRAL binding affinity should confer commensurately higher cytoprotection against atRAL. Using the CTB assay and the treatment paradigm established above (Fig. 2B), we measured cell viability in HEK293T cells expressing recombinant RHO from *H. sapiens*, *B. taurus*, and *P. alecto*. We normalized these values to the expression levels of each orthologous RHO as calculated with absorbance spectroscopy of the purified pigments (Fig. S1, Table S1). Unlike bovine, human RHO conferred no significant cytoprotection against atRAL toxicity compared to the empty vector control (Fig. 2C). Consistent with our hypothesis, *P. alecto* afforded >2-fold greater cytoprotection against atRAL stress compared to empty vector and bovine RHO. This is highly consistent with the degree of differences in atRAL binding affinity between these three species (Fig. 1J; Table S1). Thus, this provides evidence that RHO-mediated atRAL cytoprotection varies between species—with likely much more variation than we captured here— indicating that natural selection tunes atRAL binding affinity as a potential photoprotective mechanism. Together with our spectroscopic and signaling data (Fig. 1K–P, Fig. 2A), the superior cytoprotection of *P. alecto* RHO indicates that shunting bound atRAL into the non-signaling MIII state provides a natural mechanism to neutralize reactive retinal aldehydes without prolonging phototransduction.

### Modelling indicates RHO capacitance of free atRAL buffers visual cycle saturation

We reasoned that this natural variation in R* cytoprotection against atRAL stress reflects evolutionary tuning in response to species-specific photodamage risk, which scales with dim-light sensitivity (e.g., rod density, RHO expression)^40,51,167^. While humans are exposed to brighter absolute solar flux irradiance (∼25,000 cd/m^2^), our retinae are of low gain compared to that of *P.alecto* (flying fox bat), which possesses extreme scotopic sensitivity (e.g., high gain) and roosts in open canopies, exposing its highly amplified retina to severe daytime solar flux^146,147,168,169^. This is predicted to result in severe acute atRAL loads (Fig. 2D). To test whether RHO-mediated atRAL binding can limit toxic accumulation during photic stress, we constructed a deterministic kinetic model of atRAL production, clearance, and buffering (see Methods). The model integrates photic inflow (RHO activation), metabolic clearance (RDH activity), and reversible sequestration of atRAL by opsin and phosphatidylethanolamine (PE).

Under human daylight conditions, atRAL generation was conservatively estimated at ∼120 µM/min, consistent with *in vivo* densitometry and biophysical constraints (see Methods). Scaling the human inflow rate by a *Pteropus*-specific morphological gain multiplier (5x) and the ratio of *P.alecto*:human environmental luminance exposures (∼12,500 cd/m²:25,000 cd/m²), we predicted a staggering 300 µM/min ecological load for the *P. alecto* megabat (Fig. 2D). To implement a maximal stress test, we assume all solar flux reaches the photoreceptor unattenuated by the cornea and vitreous humor.

Our model estimates a highly conservative atRAL clearance ceiling of 60 µM/minute (see Methods) and assumes that opsin capacitance (*K_d_* scaled to predicted *in vivo* affinity, *K_d,phys_*; see Methods) strictly dictates the steady-state availability of the free aldehyde during photic shock (Fig. 2E). At this clearance rate, unbuffered atRAL rapidly exceeds a 50 µM toxicity threshold within ∼83 s under human load and ∼21 s under megabat load (Fig. 2F, 2G). Inclusion of RHO-mediated buffering significantly delays this accumulation. Human RHO (*K_d, phys_* ≈ 107 µM) extends time-to-threshold by ∼12-fold (15.9 minutes) under human load (Fig. 2F). The higher-affinity *P. alecto* RHO (*K_d, phys_* ≈ 12 µM) prolongs the delay further, maintaining sub-threshold atRAL levels for ∼42 minutes under human load and ∼10 minutes under megabat load (Fig. 2F, 2G). These results indicate that R* binding (capacitance) substantially buffers acute atRAL accumulation and that increased capacitance provides a protective advantage under high photic load. R* affinity seems to be finely tuned to species-specific ecological demands, as both human and megabat systems push atRAL production near clearance limits for their systems. Capacitance modulation via RHO amino acid substitutions is notably a more metabolically efficient strategy to regulate atRAL burden than overexpressing clearance enzymes.

### Synthetic active-state lock uncouples capacitance from signaling to confer extreme cytoprotection

Having established the cytoprotective and predicted physiological benefits of the natural R* capacitor, we next sought to definitively isolate the biophysical function of opsin capacitance from canonical phototransduction. Because photoactivation of wild-type RHO inherently couples atRAL sequestration with G-protein signaling, we engineered bat and bovine rhodopsins (**eRHO**) locked in the active state conformation (N2C; M257Y; D282C)^170^, and fused at the C-terminus to a high affinity peptide modelled off the C-terminus of transducin (Gt_α_CT-HA) that stabilizes the active state^101^, followed by a penultimate C-terminal outer segment trafficking signal (Fig. 3A). The bovine eRHO has been previously shown to result in a 100% atRAL-bound RHO population at steady state^101^. Using fluorescence spectroscopy, we confirmed that both eRHO variants achieved a maximal 100% all-*trans*-retinal fraction bound at steady state, completely abolishing detectable ligand release (Fig. 3B-C). Consistent with our capacitor model, both eRHO constructs conferred significantly greater cellular protection in HEK293T cells against acute atRAL toxicity than their wild-type counterparts (Fig. 3D). Given the superior protection afforded by bovine eRHO, we further validated this construct in retinal cell line models confirming that it conferred robust cytoprotection against atRAL stress in both human retinal pigment epithelial cells (ARPE-19; Fig. 3E) and mouse photoreceptor-derived cells (661W; Fig. 3F). This establishes that synthetic capacitance for atRAL –achieved via a constitutively active conformation RHO—can improve cytoprotection in retinal cell types beyond that afforded by natural WT RHO orthologs.

**Figure 3:**
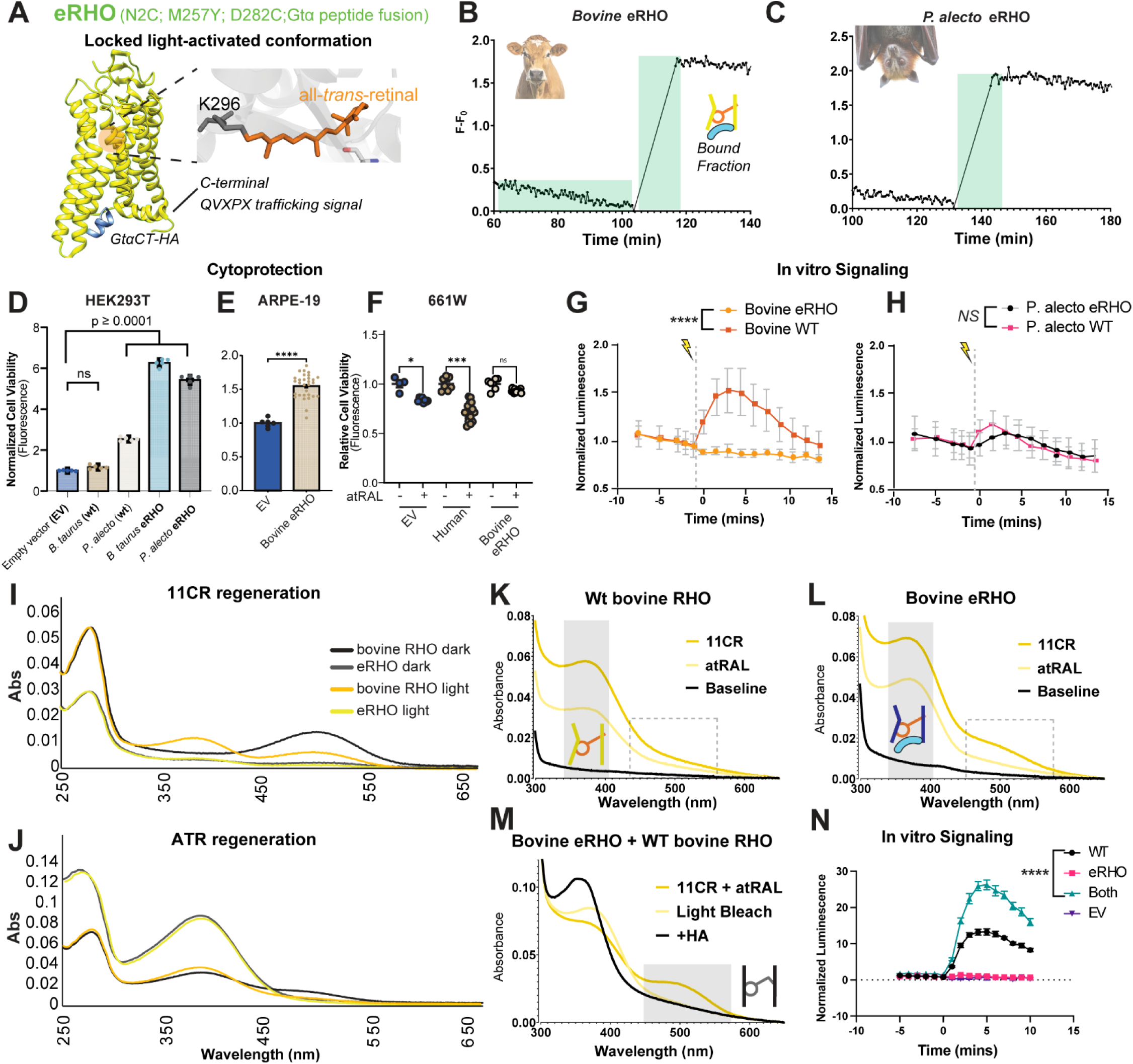
Active-state-locked synthetic RHO uncouples atRAL capacitance from signaling to confer cytoprotection and enhance wild-type RHO regeneration and signaling. **(A)** Schematic of the engineered synthetic rhodopsin (eRHO) locked in the active conformation via mutations (N2C;M257Y; D282C; native trafficking signal fused to a high affinity analogue peptide derived from the C-terminus of the α-subunit of transducin (Gt_α_CT-HA). **(B-C)** Fluorescence retinal release assays demonstrating that both Bovine and *P. alecto* eRHO constructs achieve 100% fraction bound atRAL at steady state. **(D)** Cell viability assay showing enhanced cytoprotection against atRAL for both eRHO constructs compared to their wild-type counterparts in HEK293T challenged with 50 µM atRAL. (**E-F**)eRHO also affords significant protection to ARPE-19 and 661W cells challenged with 20 µM atRAL. (**G-H**) GloSensor assay measuring Gt/Gs signaling in HEK293T cells expressing bovine eRHO vs. WT (G) or *P. alecto* eRHO vs. WT (H). Significant differences were observed for comparisons involving Bovine WT; vs. *P. alecto* eRHO (p = 0.0002), Bovine eRHO (p < 0.0001), and P. alecto WT (p = 0.0002), as well as vs. rhodopsin-absent controls (p < 0.0001). Two-way ANOVA with Tukey’s multiple comparison correction. **(I-J)** Absorbance spectra of recombinant pigments regenerated with either 11CR or atRAL **(K-M)** Absorbance spectra of recombinant apoprotein WT bovine RHO alone (*K*), bovine eRHO alone (*L*) or both apoproteins (*M*), after incubation with all-*trans* retinal, followed by 11-*cis* retinal. In (M), light bleach shows production of 380 nm peak, and subsequent treatment with hydroxylamine shifted this to ∼350nm. **(N)** Glosensor assay of HEK293T cells co-transfected with G_t_/G_s_ and either WT, eRHO, or both (1:1, holding WT RHO constant), regenerated with 9-*cis* retinal, followed by light bleach at *t=0*. Co-expression of WT and eRHO (’Both’) significantly amplified luminescence compared to WT RHO alone (****p < 0.0001; two-way ANOVA with Tukey’s multiple comparison correction).. Two-way ANOVA with Tukey’s multiple comparison correction. All cell assay read-outs (cell titer blue; luminescence) normalized to protein expression levels.

To confirm that this enhanced cytoprotection operated independently of G-protein activation, we hypothesized that the inclusion of the C-terminal G_t_ fusion peptide— which occupies the cytoplasmic transducin-binding cleft^27,171^ —would sterically occlude signaling to endogenous G_t_ (in this case, the G_t_-G_s_ chimera). Using the live-cell glosensor assay described above, we found that the bovine eRHO proved to be significantly and completely signal-silent relative to WT bovine RHO, establishing that bovine eRHO functions purely as a non-signaling, high-affinity biophysical sink for atRAL (Fig. 3G). By contrast, the *P. alecto* eRHO retained canonical G-protein signaling, even displaying evidence of light activation, with no significant difference from *P. alecto* WT RHO (Fig. 3H), indicating species-specific conformational differences in how the C-terminal fusion interacts with the cytoplasmic cleft, consistent with *P. alecto* RHO existing in an equilibrium of MII and MIII states. Together, the signal-silent bovine eRHO definitively disentangles active-state atRAL sequestration from G-protein signaling, proving that expanding the physical opsin sink alone is sufficient to rescue cells from acute retinoid toxicity.

### High-affinity atRAL sequestration enhances the regeneration and signaling of wild-type RHO

Beyond acute cytotoxicity, free atRAL acts as a diffusible agonist that rebinds empty opsin apoprotein in the dark, stabilizing an active-like state that inhibits regeneration with 11CR and generates desensitizing dark noise (see introduction). We hypothesized that a high-affinity atRAL sink like bovine-eRHO could modulate the chromophore-binding dynamics of neighboring wild-type RHO by scrubbing free atRAL from the local environment. Through absorbance spectroscopy, we first confirmed that unlike WT bovine RHO —which regenerates with 11CR to produce a classic ∼498 nm dark-state peak— bovine eRHO cannot be regenerated with 11CR, and instead requires regeneration with atRAL, displaying a 380nm absorbance peak consistent with conformational selectivity (Fig. 3I-J). However, when WT bovine RHO apoprotein was incubated with atRAL in the dark, we also observed a 380nm λ_max_ peak (Fig. 3J), confirming the spontaneous formation of an active-like opsin/atRAL complex ^19^. We reasoned that this spontaneous agonist binding directly competes with 11CR regeneration, consistent with the diminished visual sensitivity and delayed dark adaptation observed *in vivo* when atRAL clearance is impaired in humans and mice ^102–110^. We further hypothesized that the presence of eRHO may mitigate this agonist interference by sequestering atRAL, allowing endogenous WT RHO to regenerate with 11CR. We tested this by investigating the regeneration of the dark-state RHO apoprotein with 11CR after *pre-treatment* with atRAL—mimicking conditions of impaired atRAL clearance—and in the presence or absence of eRHO apoprotein. Pre-treatment of WT bovine RHO apoprotein with atRAL produced a 380 nm peak, and subsequent treatment with 11CR failed to produce a dark state 498 nm peak, demonstrating complete agonist interference (Fig. 3K). A similar observation was made with eRHO apoprotein alone (Fig. 3L). Strikingly, when eRHO and WT RHO were co-incubated at 1:1 molar ratios, WT RHO successfully regenerated with 11CR to form a robust ∼498 nm dark-state peak despite prior atRAL challenge (Fig. 3M). We confirmed the integrity of this regenerated pigment by photobleaching the sample to shift the peak to 380 nm, followed by hydroxylamine cleavage (Fig. 3M). This demonstrates that high-affinity atRAL sequestration actively shields neighboring dark-state opsins from agonist interference, facilitating dark adaptation.

We next investigated whether this competitive sink also alters the signaling dynamics of light-activated WT RHO. Because G-protein activation is maximal when R* is covalently bound to atRAL in the MII state, and bovine eRHO is completely signal-silent (Fig. 3G), any change in cellular signaling during co-expression must reflect eRHO-driven effects on WT RHO active-state dynamics. Using the GloSensor assay in HEK293T cells, we compared light-evoked signaling across cells expressing WT RHO alone, eRHO alone, or a 1:1 co-transfection of WT RHO and eRHO (holding WT RHO plasmid concentration constant). While eRHO alone failed to signal, co-expression of eRHO with WT RHO resulted in a striking ∼2-fold amplification of maximal WT RHO signaling (Fig. 3N). This signaling boost persisted even when challenged with a 2- to 3-fold molar excess of exogenous atRAL (Fig. S6). We propose that the presence of the ultra-stable eRHO sink alters local mass-action equilibria^101^, preventing the premature dissociation of atRAL from light-activated WT RHO and stabilizing its signaling-competent MII conformation. Altogether, these results demonstrate that high-capacitance atRAL sequestration dually enhances photoreceptor performance: it stabilizes the signaling state of light-activated RHO upon illumination while shielding neighboring unbleached apoproteins from agonist interference to accelerate dark-state regeneration. This suggests that high MII stability in RHO may benefit rod survival and performance, by conferring cytoprotection against atRAL induced light stress, and modulation of neighboring RHO sensitivity via accelerated regeneration and signaling.

### Convergent evolution of enhanced RHO capacitance in high-irradiance human populations

Having established that RHO capacitance of atRAL exerts major physiological effects on retinal function, and that natural selection dynamically tunes this capacitance across mammalian species to buffer specific environmental loads, we hypothesized that these same evolutionary pressures could shape human biology. To determine if human populations exhibit geographic variation in opsin capacitance tied to historical solar flux, we curated a panel of *RHO* missense variants. We primarily focused on Variants of Uncertain Significance (VUS) that have been clinically documented and have no observable defects (R21C, G51A, R69H, E122G, V157I, M216L, R252P, A333V, T320N, T340M)^32,172–174^, hypothesizing that they may reflect natural variance in atRAL sequestration instead. We also included A299S, a human allele that drives extended retinal release times and occurs naturally in the highly stable *P. alecto* (bat) RHO^142,175^. All alleles expressed and localized to the membrane when expressed heterologously as expected (Fig. S3). Following spectroscopic validation of normal expression levels, characterization of atRAL release (Fig. S1, S4; Table S1) and cytoprotection (Fig. 4A), the panel clearly segregated into two functional phenotypes: most variants displayed enhanced atRAL sequestration and cellular resistance to atRAL cytotoxicity relative to wild-type (R21C, R69H, G51A, E122G, V157I, M216L, A299S, T320N, A333V) and a minority exhibiting wild-type atRAL binding with no change in basal susceptibility (R252P, T340M).

**Figure 4:**
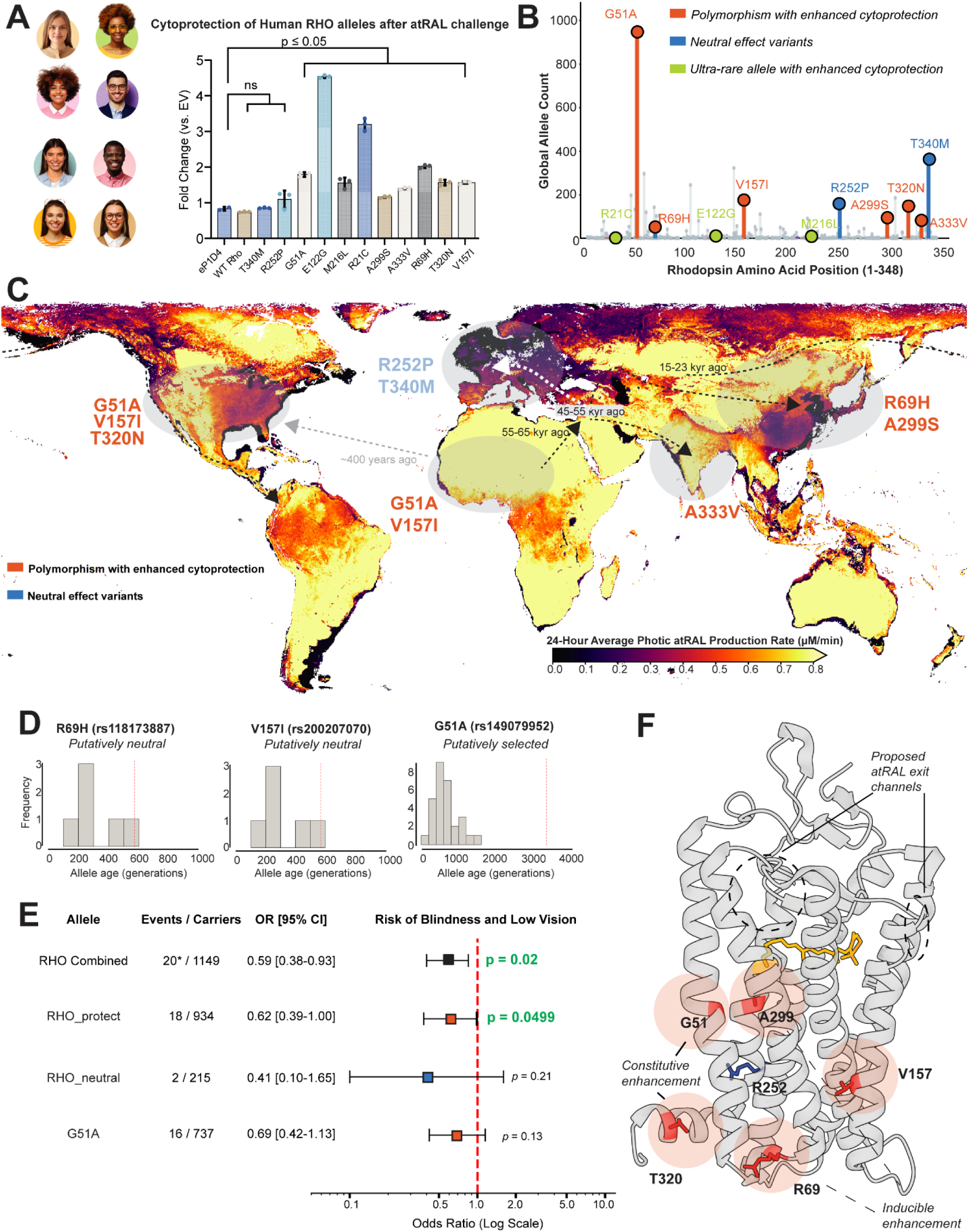
Recent human evolution of enhanced RHO capacitance is associated with high-irradiance geographies and a reduced risk of blindness. **(A)** Relative cell viability of human RHO variants in HEK293T cells challenged with 50 µM atRAL for 4hrs, normalized to protein expression **(B)** Global allele counts for the characterized RHO variants (gnomAD), ultra-rare alleles (<10) are shown, as are protective and neutral variants with greater or lesser cell viability than WT RHO. **(C)** Map denoting satellite-derived 24-hour average irradiance and expected photic atRAL production rates. Human dispersal routes are shown with dates in thousands of years (kyr). Mapping of human alleles (count >10) to genetic ancestry groups and geographical zones with the highest allele frequencies. **(D)** Allele ages (dashed vertical line) for R69H, V157I and G51A relative to a background distribution of frequency matched variants in RHO exons. G51A age estimates place it as significantly older than background (p<0.03). **(E)** Analysis of alleles and carriers in the Vanderbilt University medical biobank (BioVU) for the phenotype of blindness and low vision. Plot shows odds ratios (ORs), 95% confidence intervals, and p-values. To increase statistical power, all alleles of interest present in BioVU were aggregated into a protective (G51A, A299S, T320N, A333V, and V157I) and neutral (R252P and T340M) carrier groups **(F)** Crystal structure of Meta-II (PDB ID: 3PQR). Structural localization of convergently evolved protective variants (red) are outside of proposed atRAL exit channels and instead localize near important intramolecular interactions controlling stability of the light-activated conformation (see discussion).

To determine if this functional divergence correlated with geography or ancestry, we cross-referenced our panel with the Genome Aggregation Database (gnomAD v4.1), a large public database of global allele frequencies (Fig. 4B; parallel versions of all analyses using the 1000 Genomes database are also presented in Fig. S7). As expected, based on the high levels of constraint estimated for RHO exons (Fig. S8), all alleles are at low frequency (<1%), with a few being ultra rare (allele count <10 in gnomAD) and thus excluded from further genetic investigation (R21C, E122G, M216L). Stratifying the remaining variants by gnomAD-defined “Genetic Ancestry Group” (a proxy for continental ancestry) revealed a striking geographical pattern (Fig. 4C; Table S2). Both non-protective alleles (R252P, T340M) are significantly more common in European individuals compared to the rest of the world (Fisher’s exact test, T340M: odds ratio=2.92, p=4.00×10^-13^, R252P: odds ratio=7.09, p=1.56×10^-11^). In contrast, alleles conferring enhanced cytoprotection (G51A, V157I, R69H, A299S, T320N, A333V) are at higher frequency in the East Asia, South Asian, African/African American, and Admixed American groups. Specifically, the G51A and V157I alleles are dramatically more common in African/African American individuals relative to the rest of the world (G51A: odds ratio=30.4, p<10^-16^, V157I: odds ratio=17.96, p<10^-16^), and T320N is more common in Admixed Americans (Fisher’s exact test, T320N: odds ratio=2.28, p=0.019), and A299S and A333V are largely private to East and South Asian individuals, respectively (<10 allele counts found outside these groups). To determine if these patterns aligned with geographic light environments, we derived a continuous 24-Hour Average Photic atRAL Production Rate mapped across a global coordinate grid (Fig. 4C). This 25-year satellite-derived climatology (every February from 2001-2025) natively integrates historical diurnal cycles, atmospheric attenuation, localized surface albedo and pupillary constriction, converting raw environmental photon flux into intracellular atRAL generation rates (µM/min) via our established biophysical scaling constant. Notably, satellite retrievals across Northern and Central Europe were persistently obscured by dense winter stratus clouds, illustrating a highly attenuated, low-flux photic environment. In stark contrast, equatorial latitudes and regions characterized by arid climates or seasonal snowpack consistently registered the highest baseline atRAL generation rates, although we note that this approach is a proxy for environmental light exposure and does not directly measure atRAL.

When we combine information about which genetic ancestry groups have the highest frequencies of each missense allele with our photic map, the putative evolutionary forces maintaining RHO mutations emerge. Variants conferring enhanced cytoprotection are at their highest frequencies in groups with ancestry derived from high-irradiance geographical zones where solar flux drives the largest 24-hour average photic atRAL loads (specifically, G51A/V157I in African/African American, R69H/A299S in East Asian, and A333V in South Asian individuals). In contrast, European populations historically migrated into higher latitudes characterized by pronounced cloud cover low albedo, and canopy coverage, predicted to result in substantially reduced annual photic atRAL production rates (Fig. 4C). Thus, we hypothesize that within this low-light context, the relative absence of severe environmental retinoid stress potentially removed the atRAL clearance bottleneck, permitting functionally benign—non-protective—variants (R252P; T340M) to rise to higher frequencies unique to these relatively dark geographies.

### The G51A protective allele is ancient and under non-neutral selection

To understand whether the low frequency protective variants are being maintained by non neutral forces, we next extracted allele ages from two recent sources^176,177^. While estimates for A299S, T320N, and A333V were not available, the G51A allele that is most common in African/African American individuals arose ∼2,222–3,328 generations ago (∼55,500–83,200 years across dating models; Fig. 4D, right panel; Table S2): this age estimate is significantly older than other frequency matched variants in RHO exons (p<0.03), suggesting that G51A has persisted longer than expected under drift or purifying selection alone. In contrast, when we conducted the same analyses with allele age estimates for the R69H and V157I variants, we found that they were evolutionarily young (∼570 and ∼571 generations, respectively), with age estimates that fell within the background range of other similarly low frequency variants in RHO exons (p=0.48 and 0.52, respectively). The ancient, non-neutral retention of G51A implies that structural expansion of the *RHO* capacitor putatively provided a distinct survival advantage, buffering the toxic retinal loads generated during the human transition into extreme-irradiance environments. In summary, the geographic distribution of human *RHO* variants reveals that high-light environments can promote the long-term maintenance of certain mutations in an otherwise constrained gene.

### Mutations enhancing RHO capacitance are associated with a reduced risk of blindness

We hypothesized that if natural selection actively maintained the expanded-capacity G51A allele to buffer phototoxicity in our ancestors, this intrinsic structural mechanism should concurrently protect against the long-term, cumulative retinoid toxicity associated with modern retinal degenerations. To test this prediction *in vivo*, we interrogated the clinical phenotypes of allele carriers using the Vanderbilt University medical biobank (BioVU), focusing on end-stage visual failure—specifically the clinical phenotype of “Blindness and low vision”. Analysis of the ancient G51A allele suggested a potential protective trend against blindness (Odds Ratio [OR] = 0.69; 16 events among 737 carriers); however, due to the rarity of the allele, this individual association did not reach statistical significance (*p* = 0.13; Fig. 4E). To improve statistical power and further evaluate this association, we employed a burden-based approach by aggregating carriers of all seven functionally characterized variants available in the BioVU database (G51A, V157I, R252P, A299S, T320N, A333V, and T340M) into a combined carrier group. This combined group showed a statistically significant reduction in the risk of severe vision loss, exhibiting 41% lower odds of developing blindness and low vision compared to non-carriers (OR = 0.59; *p* = 0.02). To determine if this protective effect was specifically driven by the expanded atRAL buffering capacity we observed *in vitro*, we further stratified the cohort into two distinct functional categories: a “Protective” group comprising the high-capacitance alleles (G51A, A299S, T320N, A333V, and V157I) and a “Neutral” group comprising the functionally wild-type alleles (R252P and T340M). Strikingly, despite the reduction in statistical power inherent to splitting the cohort, the Protective group remained significantly associated with reduced risk of blindness (*p* = 0.0499; OR = 0.62). The Neutral variant group did not reach statistical significance, although the direction of effect remained protective (*p* = 0.21; OR = 0.41). Together, these clinical data support our biochemical model *in vivo*: alleles that physically expand R* capacitance to sequester toxic atRAL natively protect the human retina against progressive, blinding disease.

## Discussion

For the past century and a half, RHO’s function has been viewed primarily as visual; our findings support a reframing of RHO as more than a photosensor: our evidence demonstrates that RHO is also a light buffer, protecting against the desensitization and phototoxicity of atRAL by sequestering the chromophore post-light activation (R*). Just as an electronic capacitor intrinsically performs two functions^124^—absorbing catastrophic voltage surges^125^ while simultaneously decoupling high-frequency static to preserve signal fidelity^126^—our results demonstrate that the atRAL capacitance by R* serves this exact dual biological function. By dynamically sequestering and releasing atRAL in equilibrium, R* buffers photic atRAL “loads”, affording structural cytoprotection and sensitivity increases in a manner structurally independent of its intrinsic signaling function. Importantly, this single-receptor capacitor model represents a fundamental departure from established GPCR clearance mechanisms, discussed below. Our evidence also establishes that R* capacitance varies across species, likely tuned by physiologically variable atRAL burden. We conclude that long R* lifetimes across dim-light species are a byproduct of high atRAL binding affinity, and not necessarily an adaptation for enhancing intrinsic R* signaling as previously proposed^178^, especially since our results demonstrate that the most stable R* orthologs drive the weakest signaling (however see below sections for proposed mass action effects on signaling). Indeed, Human RHO displays robust signaling but is an intrinsically weak capacitor with a short R* lifetime that affords no significant cytoprotection. This likely reflects an evolutionary prioritization of rapid visual recovery seen in diurnal animals like ourselves^23,179^. That said, our results indicate natural human R* capacitance may have been affected by evolutionary pressures to some degree, resulting in mutations (e.g. African/African American G51A) that persist in high-irradiance geographies as genetic modifiers of blindness and low vision (with pleiotropic or epistatic costs that likely underlie their low frequency). As a whole, our results suggest that most humans experience an evolutionary mismatch in modern or high-irradiance light environments, rendering the human retina uniquely susceptible to atRAL-associated macular degenerations—a vulnerability that can be biophysically countered by expanding opsin capacitance.

### Light activated RHO balances agonist buffering with signaling

Within classical pharmacology, receptor-mediated ligand buffering is conventionally treated as a specialized evolutionary workaround restricted to non-signaling ‘decoy’ receptors^180,181^. These include the non-signaling atypical chemokine receptors which facilitate ligand clearance by β-arrestin recruitment ^118,119^, as well as the (ET_B_/ET_A_) endothelin-1 clearance pathway, which serve as a two-receptor ligand sink-signaling system thereby modulating intercellular signaling with varying spatially-determined signal efficacies that rely on virtually irreversible ligand binding followed by receptor internalization and lysosomal degradation ^182–185^. Our findings reveal that GPCR-mediated atRAL buffering—which must be reversible to promote recycling to the inverse agonist 11-*cis* retinal— does not require such spatially-restricted, multi-receptor or trans-cellular division of labor. Instead, rhodopsin acts as a self-contained, autonomous homeostatic unit. The very same signaling-competent receptor molecule that propagates visual transduction also moonlights as its *own* ligand sink (as opposed to paracrine signaling of atypical chemokine receptors and ET_B_/ET_A_), utilizing its empty apoprotein state to directly buffer the local concentration of its own toxic and desensitizing agonist. This represents an elegant, single-protein autoregulatory mechanism that bypasses the evolutionary cost of maintaining auxiliary decoy or clearance pathways. Furthermore, we must distinguish this intrinsic capacitor mechanism from geometry-driven ‘rebinding’ phenomena observed in GPCR signalosomes or lipid rafts. In densely clustered receptor domains, a dissociated non-covalent agonist is frequently recaptured by neighboring receptors due to local spatial confinement and restricted lateral diffusion, artificially slowing the macroscopic off-rate and mimicking a high-affinity state^186–190^. By contrast, our evidence shows that rhodopsin capacitance is not a passive artifact of receptor crowding. Rather, it is a conformationally driven, covalent chemical trap. Agonist sequestration is governed by the state-dependent affinity of the active conformation itself which dynamically selects and covalently neutralizes the highly reactive atRAL aldehyde through Schiff base linkage. This is further substantiated by our in vitro assays where high-affinity agonist buffering remains fully intact in detergent-solubilized, monomeric preparations, demonstrating that this capacitor phenotype is a hardcoded biophysical property of the receptor’s active-state conformational landscape rather than an emergent property of spatial clustering. It would be an interesting area of future investigation to determine if spatial clustering of rhodopsin within paracrystalline arrays^191,192^ enhances the intrinsic agonist buffering function of rhodopsin.

### Light activated RHO protects against atRAL toxicity and desensitization

Previously, RHO’s function was regarded purely as signal detection limited by thermal noise^193–195^. Through this framework, the increased stability of light-activated RHO (R*;MII) in nocturnal vertebrates was interpreted as an adaptation to enhance signaling efficiency ^140,141,196,197^. Building on evidence that R* reversibly rebinds atRAL in equilibrium^101^, our results demonstrate that evolution tunes this binding affinity across species to afford cytoprotective capacitance as a retinoid sink. This buffers atRAL photic generation against metabolically limited visual cycle clearance rates, thereby serving a role similar to PE and RGR ^89,116,117^. As established by Schafer et al., atRAL acts as an agonist that exclusively rebinds to the active (Ops*) conformation, while 11CR selectively binds the inactive (Ops) conformation^19,101^. Thus, what appears as a temporal delay or a steady-state ‘trapping’ of retinal reflects a highly dynamic equilibrium where atRAL continually releases and rebinds to the receptor. Because the active Ops* conformation can transiently persist after the atRAL ligand dissociates, it continually draws free atRAL back into the binding pocket^101^. By evolutionarily stabilizing this empty Ops* state—as seen in the *P. alecto* RHO and the G51A human variant—nature effectively tunes the receptor to act as a sustained retinoid sponge. Crucially, our UV-Vis kinetic analyses reveal that in *P. alecto* RHO, this sustained retention is reinforced by an atRAL-sensitive shift from MII into the non-signaling MIII photointermediate (Fig. 1M–P). Long regarded as a slow, vestigial decay product^91,145,157^, MIII retains covalently bound atRAL via a re-protonated Schiff base without engaging transducin^91,145^, allowing dim-light specialists to expand retinoid buffering capacity while avoiding aberrant G-protein activation. Thus, our experimental and evolutionary findings validate long-standing theoretical predictions that MIII serves a direct physiological role in buffering toxic atRAL during bright-light bleaches^198,199^. This is most incumbent on species with high rod sensitivity that are exposed at times to light, like the visual megabat *P.alecto* which roosts in open canopies, exposing its highly amplified retina to severe daytime solar flux^146,147,168,169^. This tunable sequestration confers protection against atRAL induced cell death, with more stable R* showing the greatest increase in cell viability. It will be of major interest to characterize R*-mediated cytoprotection across animal visual ecologies to test this hypothesis further. The association of high-capacity alleles with a reduced risk of low vision or blindness, occurring within high irradiance human populations, strongly suggests that a major physiological function of the opsin structure under bright light may be to sequester toxic chromophore and protect the retina.

This intrinsic buffering capacity also has implications for scotopic sensitivity, with major evolutionary and translational implications. Following photoactivation, if free atRAL is not rapidly cleared from the membrane, it diffuses back into the empty receptor to form a non-covalent *opsin/atRAL* complex^98,99^. While distinct from the MII state, this complex activates transducin at a rate several orders of magnitude higher than empty opsin^96,98,99^. As demonstrated in *Abca4-/-* knockout models, failure to clear this diffusible agonist results in the persistent activation of the visual cascade, manifesting as elevated dark noise and severe delays in dark adaptation^102,104,113^. Indeed, our results demonstrate the unnaturally stable and signal-silent eRHO affects the chromophore binding dynamics of WT RHO in both the apoprotein state, and in the light activated state, increasing regeneration and signaling respectively. Firstly, our evidence indicates that eRHO shields endogenous WT RHO apoprotein from agonist interference, thereby promoting regeneration of the dark state with 11CR. This dark adaptation is required for maximal photoreceptor signaling in the scotopic ERG flash intensity series^200^. This implies that eRHO can potentially rescue the delayed dark adaptation defects and diminished quantum-catch typical of atRAL clearance diseases such as Stargardt disease and *ABCA4*-mediated macular degenerations^102–105^. Secondly, bovine-eRHO—which cannot signal—dramatically increased the signaling of WT RHO. Critically, eRHO through high active-state stability, pushes the equilibrium towards atRAL sequestration as previously established^101^. We propose that in this mass action context, endogenous WT RHO, when light-activated, is also pushed away from atRAL release by eRHO and towards rebinding and reformation of the signaling competent MII state. It is important to note that arrestin is missing from our *in vitro* assays, whereas *in vivo*, arrestin terminates signaling within milliseconds^201^. Thus, our observation of increased WT signaling *in vitro* due to increased MII lifetime may be an artifact which does not translate *to in vivo*.

Nevertheless, this is mechanistically consistent with our observations of eRHO modulating the chromophore binding dynamics of WT endogenous RHO. We propose that eRHO acts to stabilize endogenous MII after light activation, but once the latter conformationally decays, eRHO then serves a second function in promoting recovery of the dark state by mitigating agonist interference. It is tempting to speculate that *P. alecto* evolved an increased MIII subpopulation for this exact reason: noise mitigation without risk of signaling from a re-formed MII, thus preserving dim-light sensitivity. This putative elegance afforded by high R* stability is a testament to the brilliance of evolution and leveraging it into translational approaches is an exciting area of future investigation.

This boost in sensitivity is of clear evolutionary benefit for dim-light specialists, where full light bleaches are exceedingly rare, and instead maximizing signal over dark noise is the critical variable for maximizing visual performance^202,203^. While our experiments utilize an artificially heterogenous receptor population *in vitro* (eRHO + WT RHO), in a given dim-light species, these ecologically relevant partial and even single photon bleaches would be expected to result in an analogously heterogenous mixture consisting of a sub-population of R* and neighboring dark state receptors^204,205^. Our results indicate that high R* stability may have therefore evolved to shield neighboring dark state receptors from agonist interference, which results in dark-state receptors spontaneously binding atRAL as we demonstrated. This shielding would promote dark adaptation and minimize dark noise—phenotypes of major importance for dim-light species visual performance. This raises the possibility that high R* stability evolved for sensitivity, as previously proposed^141^, yet through different and counter-intuitive mass action mechanisms hitherto not described (e.g. mitigating agonist interference post-bleach). If so, it is possible that the cytoprotection afforded by R* stability and atRAL sequestration, may be a convenient side-effect, rather than the original target of natural selection. We propose that species such as humans which do not historically operate in photon-scarce natural conditions relative to other species, may not normally benefit from the effects of high R* stability—faster dark adaptation and increased signaling intensity—consistent with human RHO being unstable and affording very low atRAL sequestration. In the modern era, however, atRAL-associated macular degenerations are highly prevalent and represent a leading cause of global blindness ^206^, and we propose the lack of a cytoprotective R* as seen in other species, may be causally associated. This is supported by our finding that human patients with alleles that increase atRAL sequestration and cytoprotection display a significantly reduced risk of blindness. Thus, among species, humans may therefore be uniquely suited to benefit from synthetic atRAL capacitance for its cytoprotective effects, which would constitute an artificial rebalancing of this pleiotropic trade-off, away from diurnal visual performance and towards visual resilience.

### Structural mechanisms of R* capacitance for atRAL

Importantly, our investigation of human alleles reveals insights into the structural and functional basis of enhanced atRAL sequestration. The location of alleles that enhance sequestration (Fig. 4F) suggests they do *not* exert their effects through steric occlusion of the proposed exit channel^207^ but rather stabilize intramolecular hydrogen bonding networks that regulate atRAL hydrolysis and binding affinity. Originating in distinct, geographically isolated populations, we propose that these alleles (G51A-African/African American, R69H/A299S-East Asian, and A333V-South Asian) convergently afford enhanced cytoprotection via divergent structural and functional mechanisms. Specifically, G51A, R69H and A333V alleles act as ‘constitutive sinks’ that sequesters roughly 50-300% more atRAL than WT under basal conditions (Fig. S4; Table S1), whereas A299S and T320N function as an ‘inducible sponge’ that matches WT under low atRAL but massively upregulates sequestration during atRAL stress by ∼3-fold (relative to WT at ∼2-fold; Fig. S4; Table S1). Structurally, G51A and A299S interact with the N55-D83 intramolecular hydrogen bond network that stabilizes the Schiff base and the kinetics of the active state^148,208,209^. Meanwhile, previous modeling indicates that R69H and T320N converge to destabilize the same structural nexus: the interface between Intracellular Loop 1 (ICL1) and Helix 8. In the dark state, wild-type T320 anchors Helix 8 through specific hydrogen bonds to TM2 (Q64) and ICL1 (H65)^210^. The bulky T320N substitution likely ruptures this tethering network, while R69H concurrently alters the electrostatic landscape of ICL1 itself. By uncoupling the packing between ICL1 and Helix 8, both mutations likely relieve physical restraint on the nearby NPxxY motif, thereby likely favoring the active Meta II conformation^211^. Moreover, site A333 is at the junction where the C-terminal tail becomes unstructured^212^, and we speculate that a loss of flexibility caused by A333V may decrease the conformational entropy of the unstructured C-terminal tail. The fact that geographically isolated populations convergently evolved divergent structural solutions to overcome similar environmental stressors point to historical environmental light stress exerting a genuine selective pressure on human retinoid handling, consistent with our population genetic analyses.

### Evolutionary and cellular context of R* capacitance

The evolutionary importance of the RHO capacitor becomes evident when contrasting the ancient divergence of rod and cone photocycles. Cone opsins are evolutionarily tuned for rapid decay of the active state, releasing atRAL almost immediately to maintain rapid temporal resolution and visual recovery^18,23,213,214^. Rods, by contrast, operate in a high-gain system with extensive, opsin-dense membrane surfaces and photon pooling ^158^. Rods are highly susceptible to transient atRAL accumulation after moderate light bleaches of ∼12-40%^50,51^. To safely prevent free atRAL from overwhelming the cell, rhodopsin evolved a highly stable R* intermediate capable of sequestering atRAL with high affinity, regulating its release into rod cytosol. This evolutionary process is a defining feature of the functional divergence of rhodopsin from the cone opsins and appears to have increased further during vertebrate and mammalian diversification^18,123,141,196^. This delayed release comes at the cost of slower regeneration but we show this cost enables the benefit of reduced atRAL-associated phototoxicity. This trade-off may be uniquely skewed in rods, as cones appear less susceptible to such degeneration, as they possess an expanded visual cycle that expedites atRAL clearance to support the rapid retinoid recycling they require, consistent with the death of rods typically preceding that of cones in age-related macular degeneration^94,95^. A valuable future venture would be to investigate buffering capacities of cone opsins.

Our model of intrinsic RHO capacitance is complementary with the role of visual arrestin (Arrestin-1; Arr1) promoting re-binding of atRAL when bound to phosphorylated opsin (OpsP)^92,121,144^. While Arr1 has therefore long been implicated in photoprotection, our data demonstrate that atRAL buffering is not governed by a single mechanism but instead reflects coordinated tuning of both opsin stability and arrestin engagement. Evolutionary pressures appear to tune the intrinsic thermodynamic stability of the R* apoprotein itself to regulate atRAL sequestration, while arrestin provides an additional, regulatable layer of control over ligand retention and release. This bipartite system may confer remarkable evolutionary tuning strategies, such as in the sperm whale, which possesses an unstable R* that likely facilitates only weak atRAL binding affinity^215^, yet has an Arr1 that drives enhanced atRAL sequestration in R*^122^. Our results also raise the intriguing possibility that enhanced Arr1 binding in dim-light specialists evolved to minimize atRAL-induced noise, which would be of clear benefit for these organisms’ visual performance. More broadly, variation in arrestin-opsin interactions may tune not only atRAL handling but also kinetics of phototransduction, linking photoprotection with control of signaling noise and temporal resolution of visual signaling. Systematic comparison of RHO and Arr1 orthologs – preferably in native rod outer segments – would be required to resolve how these axes are balanced across visual ecologies.

### Enhancements to R* capacitance during recent human evolution

Our study reveals that RHO capacitance is maximized in species at high photodamage risk, such as the flying fox bat, due to a combination of rod-dense retina and moderate solar flux during tree roosting^146,147,168,169^. While human populations do not typically vary in rod complement to our knowledge, populations do vary in environmental light exposure. Our analysis of high-resolution 2001–2025 satellite data quantified regional irradiance differences and revealed that protective alleles occur overwhelmingly in high irradiance populations (G51A, V157I, A299S, T320N, A333V, R69H). However, this modern map likely represents a conservative lower bound of the true historical selective pressure. Dating indicates that G51A arose ∼2,222–2,566 generations ago (∼55,500–64,100 years; Table S2), broadly coinciding with the African Middle Stone Age—an epoch defined by climatic volatility and massive desertification during which anatomically modern humans shifted from foraging behaviors in shaded forests to diurnal persistence hunting in high-albedo savannas, likely drastically increasing their daily environmental photic load, although the magnitude of these effects is uncertain^216,217^. It stands to reason that opsin variants would be beneficial during this period, as early-onset visual impairment could reasonably be expected to drastically reduce foraging capacity.

While we had insufficient information to date the A299S and A333V alleles, we determined that they along with R69H, are at highest frequencies in East Asian and Indian populations established along Out-of-Africa expansion routes (∼55,000-45,000 years ago; Fig. 4C^218^), specifically the northern dispersal into Central and East Asia, and the Southern Dispersal Route into the Indian subcontinent. In East Asia during the Pleistocene, the landscape was dominated by the treeless Mammoth Steppe where the absence of a light-absorbing forest canopy would have generated unrelenting snow albedo during winter months^219^. Similarly, migration into the Indian subcontinent coincided with an epoch punctuated by recurrent monsoon failures and protracted aridity^220,221^. The resulting persistence of the Thar Desert and open, arid scrublands likely subjected early South Asian populations to intense, unattenuated sub-equatorial solar flux and high mineral albedo^221^. Furthermore, modern satellite irradiance estimates across East and South Asia are heavily attenuated by contemporary anthropogenic aerosols (global dimming^222,223^).The raw photon flux driving the evolution of these expanded capacitors was likely substantially harsher than modern climatology suggests. This ancient climatology combined with current geographic distributions suggests that A299S and A333V may have resisted purifying selection alongside G51A, independently expanding the R* capacitor to buffer the severe environmental photic loads encountered by early migrating populations.

Interestingly, our screen identified ultra-rare variants (E122G, M216L, and R21C, all with allele counts <50 in gnomAD) that conferred robust atRAL protection. We hypothesize that these represent instances of biophysical trade-offs. While these specific amino acid substitutions incidentally confer a biochemical resistance to retinoid toxicity, E122G introduces a massive blue shift in spectral sensitivity (λ_max_ = 478.4 nm vs. 494nm for WT; Fig. S1) which likely historically impaired visual function enough to restrict them to near-zero frequencies globally.

### Clinical implications of natural and synthetic atRAL capacitance

These protective alleles identified from high irradiance populations are low frequency, indicating that human RHO remains under strong purifying selection to maintain rapid atRAL release, likely to favor regeneration of the dark state, similar to the cone opsins and consistent with the diurnal visual ecology of humans^23^. Our results indicate that this evolutionary prioritization of recovery over protection has devastating clinical consequences in modern populations. The consequences of unbuffered atRAL are best exemplified by Stargardt macular dystrophy, a congenital disease caused by mutations in the ABCA4 transporter. ABCA4 normally facilitates clearance of N-ret-PE from photoreceptor disc membranes, and its dysfunction leads to accelerated accumulation of atRAL and toxic downstream byproducts such as A2E^224,225^. Accumulation of A2E and other lipofuscin fluorophores is a defining feature of Age-related Macular Degeneration (AMD). Beyond atrophic degeneration, the consequences of atRAL accumulation may extend to vascular pathology^67^ suggesting that R* buffering capacity may influence susceptibility to exudative (wet) AMD and diabetic retinopathy. The significantly reduced risk of blindness or low vision we detected in carriers of high R* capacitance alleles, suggests that the ancestral prioritization of visual recovery (low R* capacitance) now constitutes an evolutionary mismatch within modern light environments where atRAL-associated macular degenerations represent a leading cause of global blindness ^206^. Our results also mirror epidemiological data, which has long established that, across varied geographies, individuals of African and Asian ancestry have a significantly reduced incidence of late-stage Age-Related Macular Degeneration (AMD)^226^. While this protection in individuals of African descent is largely attributed to higher overall ocular and choroidal melanin content— which acts as a physical shield against light toxicity^227–229^—this speaks to the necessity of photoprotective mechanisms. The existence of p.Gly51Ala provides a genetic “proof of concept” for a melanin-independent protective pathway. Together, the natural optimization of these variants supports the hypothesis that reducing atRAL-induced cytotoxicity is a universal, viable therapeutic strategy for preventing retinal degeneration across all ancestries.

By directly buffering atRAL, modulation of RHO capacitance may potentially overcome many of the limitations of current therapies, which primarily target the visual cycle and have failed to offer a safe and effective treatment option^11,230–232^. Gene replacement strategies for *Abca4* in patients with Stargardt disease have been limited by vector packaging constraints given the large size of the gene^233–235^. Furthermore, restoring ABCA4 function would facilitate atRAL clearance but may be insufficient to resolve acute atRAL burden under photic stress given that RHO exceeds ABCA4 by ∼120-fold and ABCA4 shows only moderate maximal transport in disc membranes (Vmax ≈ 4.9 nmol·min⁻¹·mg⁻¹)^236^. Other therapies, such as Syfovre and Izervay address downstream complement-driven inflammation, a key component of AMD and retinal degeneration pathophysiology; however, limiting the initial accumulation of toxic retinoids may offer a more effective strategy than intervening at later stages of disease progression^32^. It should be noted that because the condensation of atRAL into N-ret-PE is concentration-dependent, the continuous siphoning of free atRAL by the RHO capacitor is predicted to starve the formation of downstream bisretinoids like N-ret-PE and A2E. We also suspect that buffering free atRAL can rescue the delayed recovery of dim-light sensitivity often seen in Stargardt patients, a symptom we propose is caused by ‘dark signaling’ (i.e., the light-independent activation of dark state RHO into R* apoprotein followed by atRAL binding).

Ultimately, framing rhodopsin as not merely a photosensor but a tunable cytoprotective buffer opens a therapeutic avenue for STGD and AMD upstream of current approaches. While these other interventions primarily target the downstream sequelae of retinal degeneration—such as complement-driven inflammation— deploying optimized eRHO variants to intercept free toxic aldehydes represents a direct, biomimetic solution. Our characterization of intrinsic RHO capacitance parallels both pharmacological and naturally occurring strategies for mitigating retinal degeneration. Current therapeutic approaches include primary amines such as retinylamine or Emixustat, which function as chemical scavengers that transiently sequester free atRAL similar to the neutralizing Schiff base of R* ^11,67,237^. However, these ATR-sequestrants have major side effects on normal human vision (dyschromatopsia and delayed dark adaptation) by inhibiting RPE65 ^11,230–232^. eRHO has a clear advantage in this regard, as an evolved specific capacitor of atRAL without off target effects on the visual cycle.

### Limitations and Future Directions

It is useful to list caveats/limitations in our experimental designs. First, we purified RHO in detergent micelles, a condition known to favor the Meta II state ^121,155,238^; while our cross-species comparisons remain internally consistent, validation in native disc membranes or nanodiscs will be essential to better approximate physiological equilibria and kinetics. Second, extensive *in vivo* experiments will be required to determine whether R* stability and retinal sequestration provide meaningful protection to photoreceptors over long timescales in the intact retina.

## Materials and Methods

### RHO Plasmid synthesis

RHO coding sequences from *H. sapiens*, *B. taurus*, and *P. alecto*, were acquired from NCBI gene database and altered to contain a 1D4 affinity tag (TETSQVAPA) at the c-terminal loop. Sequences were synthesized using IDT™ gBlocks™ and cloned into p1D4-hrGFP II (Addgene, Plasmid #71317) using HIFI assembly (NEBuilder® HiFi DNA Assembly and NEB® 5-alpha Competent E. coli). RHO seqs were first verified through sanger sequencing (GENEWIZ from Azenta Life Sciences) and whole plasmids were then verified using long read sequencing (PlasmidSaurus).

### RHO purification and spectroscopic assays

Respective RHO sequences were transiently transfected using polyethylenimine lipofection at 1.5μg PEI and 15µg plasmid DNA per 10cM dish using HEK293T cells (obtained from ATCC) at 90 to 95% confluency. 24-hours post transfection media was refreshed, and cells were harvested 24-hours later at room temperature. Cells were harvested from plates using buffer made of PBS, 10 mg/mL aprotinin (RPI), 10 mg/mL leupeptin (RPI), and washed twice over. Cells went through 2-hour incubation and nutation at 4°C with 5μM 11CR (provided by Dr. Rosalie Crouch at the Medical University of South Carolina). Post retinal regeneration, cells were solubilized in buffer containing 50 mM Tris pH 6.8, 100 mM NaCl, 1 mM CaCl2, 1% dodecylmaltoside, 0.1 mM PMSF) for 2 hours followed by overnight immunoaffinity purification using Rho 1D4-Tag Affinity MagBeads (Cube Biotech). Beads were washed three times with initial wash buffer (50 mM Tris pH 7.0, 100 mM NaCl, 0.1% dodecylmaltoside) and twice over with a secondary wash buffer (50 mM sodium phosphate, 0.1% dodecylmaltoside; pH 7.0). RHO proteins were eluted the following day with 1D4 peptide (Cube Biotech) at 5mg/ml for 2 hours. All steps, including buffers and incubations, were done at 4°C. Recombinant RHO protein were validated and quantified through absorbance measurements using Cary Spectrophotometer (Agilent Cary UV-Vis compact Peltier) at 20°C.

RHO decay via retinal release can be monitored by measuring the intrinsic fluorescence of tryptophan 265, which is located near the retinal-binding pocket of RHO. Tryptophan exhibits distinct excitation and emission maxima at 295 nm and 330 nm, respectively. In the presence of bound retinal—either 11CR or atRAL—fluorescence is quenched due to energy absorption by the chromophore. Fluorescence increases only upon release of retinal, allowing for a clean and direct readout of retinal dissociation. This system enables real-time tracking of apo-protein formation through discrete increases in tryptophan fluorescence over time. The resulting fluorescence curve follows a first-order exponential rise-to-maximum, which is fitted by the equation *y* = *y*_0_ + *a*(1-*e^-kt^*) where half-life is calculated as *t_1/2_* = ln(2) / *k*. While MII decay and retinal release are mechanistically distinct, prior studies^101^ have shown that these events are tightly coupled, with retinal release occurring shortly before full MII decay.

To quantify the binding capacity of the “sink,” the fraction of protein bound (θ) at steady state was determined. This parameter characterizes the magnitude of the fluorescence increase upon the addition of hydroxylamine (HA). For each experimental condition (e.g., control and 2µM exogenous retinal), θ was calculated as θ = HA Jump/F_post_ where HA jump represents the absolute change in fluorescence following HA addition, and F_post_ is the maximum fluorescence plateau reached at the end of the assay. The apparent dissociation constant (K_d,app)_ was calculated for each condition using a quadratic binding equation to account for total ligand and protein concentrations. The total protein concentration (R_total_) was defined as 0.25μM. The total ligand concentration (L_total_) was calculated as the sum of the endogenous retinal (assuming 1:1 regeneration, 0.25μM) and the exogenous atRAL added (e.g., 2μM) resulting in an L_total_ of 2.25μM. K_d, app_ was derived as follows: K_d,app_ = (1-θ) (L_total_ - (θ • R_total_))/ θ.

The buffering duration, represented by the time constant τ, was determined from the observed half-life t_1/2_ of the regeneration or decay process for each respective condition. The time constant was calculated using the relationship: τ = t_1/2_ / ln(2)

Retinal release was measured by diluting recombinantly purified RHO to a final concentration of 0.25 μM in 400 μL of wash buffer 2. When testing with additional atRAL, the solution was brought to a concentration of 2µM. Fluorescence measurements were performed using a Cary Eclipse fluorescence spectrophotometer (Agilent) with the temperature maintained at 20 °C. The instrument was set to acquire readings every 30 seconds with a 2-second integration time, using an excitation wavelength of 295 nm (1.5 nm slit width) and emission detection at 330 nm (10 nm slit width).

Baseline fluorescence was recorded for 5 minutes before photoactivation using a fiber optic lamp (Fiber-lite, Dolan-Jenner). No detectable activation occurred from the excitation beam prior to light exposure. After photoactivation, fluorescence was monitored continuously until the rate of increase decelerated and plateaued for approximately 5 minutes. When quantifying RHO sequestration of atRAL, measurements were paused and hydroxylamine (HA; Alfa Aesar, Thermo Fisher Scientific, Waltham, MA, USA) was added at a ∼50 mM final concentration, corresponding to a large molar excess over RHO. Measurements then resumed and continued until a second fluorescence plateau was observed, typically within 10 minutes.

The kinetics of RHO regeneration were monitored via absorbance spectroscopy using an Agilent Cary UV-Vis spectrophotometer equipped with a compact Peltier temperature control unit. Bovine wild-type and engineered RHO variants were purified according to the previously described protocol, with the specific exclusion of the retinal regeneration step to maintain the protein in its apo-opsin form. Specifically, 11CR was omitted for WT RHO, and atRAL was omitted for engineered variants. Since opsin concentration cannot be determined directly via UV-Vis without a bound chromophore, concentrations were derived from previous purification yield data. To initiate the assay, equimolar amounts of each opsin (50 pmol) were combined in a cuvette. The samples were incubated at 20°C, and baseline absorbance spectra were recorded every minute for 5 minutes. Following the fifth baseline scan, 50 pmol of atRAL was added to the apo-protein mixture. The reaction was monitored continuously for 15 minutes. Finally, 11CR was added to allow for final regeneration, and spectra were recorded for an additional 10 minutes to reach equilibrium.

To monitor Metarhodopsin III (MIII) formation following light activation, UV-Vis absorbance spectroscopy (Agilent Cary UV-Vis compact Peltier) was performed across the 250–650 nm spectrum. MIII was distinguished by its characteristic λmax at ∼480 nm. An initial dark-state scan of 1 μM rhodopsin in 0.1% dodecylmaltoside (DDM) and the secondary wash buffer was recorded as a control. For conditions containing an 8-fold excess of all-*trans*-retinal (8 μM atRAL), the baseline atRAL absorbance was blanked prior to scanning to eliminate spectral overlap with MII RHO (λmax ∼380 nm). Following dark-state confirmation, samples were light-bleached for 30 seconds using a gooseneck fiber optic illuminator (Fiber-lite, Dolan-Jenner). Kinetic absorbance spectra were subsequently recorded every 30 seconds for 3 hours at 20 °C. The scan rate was set to 3000nm/minute and averaging time was 0.02 seconds. Data processing and kinetic analysis were performed using GraphPad Prism.

### Cell Viability Assays

HEK293T cells were cultured at 37 °C with 5% CO₂ in Dulbecco’s Modified Eagle Medium (DMEM; Gibco) supplemented with GlutaMAX™, 10% (v/v) fetal bovine serum (FBS; Gibco), and 1× penicillin–streptomycin (Gibco). Cells were seeded at 2.0 × 10⁴ cells/well in 100 µL of complete medium into black, tissue-culture– treated, 96-well skirted plates (Olympus) 24 h prior to transfection. HEK293T were transfected with the p1D4-hrGFP II expression construct encoding the rhodopsin sequence from the species of interest using polyethyleneimine (PEI) as described previously, without media exchange post-transfection. At 48 hours post-transfection, cells were incubated with 5 µM 9-cis-retinal (9CR; Sigma-Aldrich) diluted in complete medium (final DMSO content negligible) for 2 hours under light-protected conditions. Following the 9CR incubation, cells were light-bleached using a fiber optic lamp (Fiber-lite, Dolan-Jenner) containing an IR filter to minimize heat. Immediately after bleaching, cells were treated with 50 µM all-*trans*-retinal (atRAL; Sigma-Aldrich), diluted directly into complete medium to maintain negligible DMSO content, for 4 hours under light-protected conditions. Three hours after atRAL addition (i.e., one hour before the endpoint), 20 µL of CellTiter-Blue® reagent (Promega) was added directly to each well (final ratio 1:5, reagent: medium) according to the manufacturer’s instructions. Plates were returned to the incubator (37 °C, 5% CO₂) for >1 hour, protected from light, before fluorescence reading. Fluorescence was measured using a BMG LABTECH VANTAstar plate reader with the pre-established resorufin protocol: excitation at 544 nm, emission at 590 nm, single flash, single scan per well, and no well scanning pattern. Background signal was measured in blank wells (medium + CTB without cells) and subtracted from all readings prior to analysis. To normalize expression of retinal-binding competent RHO mutants, each sample was purified across ten 10-centimeter plates. Absorbance values were then measured for each variant and the corresponding λ_max_ was used as a normalization factor to account for variations in expression levels, allowing for adjustment of cell viability results.

ARPE-19 cells were cultured at 37 °C with 5% CO₂ in Dulbecco’s Modified Eagle Medium (DMEM; Gibco) supplemented with GlutaMAX™, 10% (v/v) fetal bovine serum (FBS; Gibco), 661W were cultured at 37 °C with 5% CO₂ in Dulbecco’s Modified Eagle Medium (DMEM; Gibco) supplemented with GlutaMAX™, 10% (v/v) fetal bovine serum (FBS; Gibco), Hydrocortisone 21-hemisuccinate (40ng/mL; Sigma H-2270), Progesterone (40ng/mL in 200 proof ethanol; Sigma P-8783), Putrescine (16mg; Sigma P-7505), and Beta-mercaptoethanol (20uL Sigma M7522).Both 661W and ARPE-19 were seeded at a density of 2.4 x 10^7^ cells/mL and electroporated using the Neon NxT Cell Electroporation System (10 uL kit, ThermoFisher Scientific). Electroporation was performed with 1ug plasmid DNA per reaction (1200V, 2 pulses, 20ms). Assay was then conducted identical to HEK293T cells except at 24hrs after electroporation.

### Analysis of Human rhodopsin alleles

We downloaded allele counts per variant of interest from gnomAD v4.1.1, which includes exome and whole genome sequencing data from the following predefined genetic ancestry groups: Middle Eastern, Remaining, Admixed American, African/African American, European (non-Finnish), South Asian, Ashkenazi Jewish, East Asian, European (Finnish), Amish. We excluded variants for which the minor allele had <50 allele counts reported globally, as these ultra rare variants are challenging to parse in terms of evolutionary or population genetic patterns. For the remaining variants (G51A, A299S, A333V, T340M, R252P, T320N, V157I), we used Fisher’s exact tests to establish differences in the frequency of the minor allele between a focal genetic ancestry group (for example, African/African American) and all other groups represented in gnomAD. We also confirmed patterns of population specificity by extracting allele frequency information for the same variants from a combined dataset of whole genome sequences from the 1000 Genomes Project (1kGP) and Human Genome Diversity Project (HGDP)^239^. These databases revealed highly similar patterns of global variation compared to our results from gnomAD (Fig. S7).

Given the high levels of constraint on human RHO exons, the mutations of interest are all at very low frequencies. As a result, traditional tests for positive selection (e.g., population branch statistic, integrated haplotype score), which typically focus on variants with minor allele frequencies >5%, are not appropriate^240^. Instead, we asked whether any of the low frequency mutations in this constrained gene have been tolerated longer than would be expected under drift or purifying selection alone. To do so, we downloaded allele age estimates^177^ for all SNPs in RHO exons; within this set, age estimates were available for two protective variants (G51A, R69H) and one non-protective variant (V157I). To derive empirical p-values for each variant, we calculated the proportion of allele age estimates that were greater than the focal variant, focusing on a background distribution of frequency matched variants in RHO exons.

### eRHO design

*Bos taurus* (bovine) and *Pteropus alecto* (bat) *e*RHO were locked in the active conformation through substitution on transmembrane helix 6 (TM6) (M257Y)^170^, stabilization through disulfide-cross linking (N2C; D282C), and chimeric fusion of the C-terminal tail mimetic from the Gt alpha subunit (Gt_α_CT-HA; EEVLEDLKSVGLF)^101^. Briefly, the N-terminal tail is connected to the third extracellular loop (EL3), and by extension TM7, by the disulfide bridge formed between cysteines at site 2 and 282. The thermal stability of the active conformation is greatly increased by the constraint of the disulfide bridge allowing opsin to stably sample MII. Steric hindrance is caused by the tyrosine substitution between TM6 and TM7 forcing an outward shift of the TM6 helical bundle. The outward shift, also seen in MII, allows for opening of the cytoplasmic face of the apoprotein and therefore binding of Gt_α_CT mimetic. The mimetic fused at the opsin C-terminus preferentially binds the inner cleft of the RHO acting as a wedge and sterically maintaining the outward tilt of TM6. Finally, an additional trafficking sequence (TETSQVAPA) was appended after the Gt_α_CT fusion (this construct is modeled off that of Schafer et al ^101^).

### GsX GloSensor cAMP Assay

Rhodopsin–G protein signaling, were assessed through a live-cell cAMP reporter assay adapted from Ballister et al^156^. using the GloSensor™ 22F system (Promega). HEK293T cells were transfected with plasmids encoding the opsins of interest, the GloSensor cAMP reporter, and Gst chimeras in which the C-terminal 13 amino acids of Gαs were replaced by those of target Gt, as described previously. HEK293T cells were co-transfected with Lipofectamine at a 100:100:1 ratio (Opsin:GloSensor:Gst), then incubated with 9-cis-retinal (10 µM), and pertussis toxin (125 ng/mL) overnight. The following day the cells were seeded into a 96-well white plate under red light conditions and treated with luciferin. The cells were read by a plate reader for a baseline reading, then light activated, and luminescence was recorded in real-time using a plate reader with cAMP levels inferred from the change in GloSensor signal following stimulation.^156^

### BioVU cohort and whole-genome sequencing

Analyses were conducted in the Vanderbilt University Medical Center BioVU biobank using participants with available whole-genome sequencing (WGS) data. Sequencing libraries were prepared using the Illumina DNA PCR-Free Prep protocol and sequenced on Illumina NovaSeq 6000 instruments to a mean coverage of approximately 30×. Reads were aligned to the GRCh38 reference genome using the DRAGEN pipeline, and variant calling was performed using DRAGEN germline workflows with joint genotyping. Standard sample- and variant-level quality control procedures were applied, including removal of samples with low coverage, excess heterozygosity, or discordant genetic and reported sex. Variants were restricted to those passing quality filters (e.g., genotype quality ≥20, depth ≥10, and passing variant-level filters). Additional checks were performed to identify sample swaps, relatedness, and contamination. A total of 245,351 participants were included in the final analysis. IRB #240655 (Gianni Castiglione).

### Blindness and low vision analysis in a human cohort

Blindness and low vision were defined using electronic health record–derived phecodes, constructed as a composite phenotype based on a predefined set of related diagnostic ICD codes (listed in Table S3). The rho_combined variable was modeled under a dominant framework, where individuals carrying at least one of the specified protective alleles were coded as 1 and non-carriers as 0. Association analyses were performed using logistic regression models adjusted for age, sex, and the top 10 principal components of ancestry. All analyses were conducted in R (version 4.3.1).

## Data, Materials, and Software Availability

All experimental spectroscopic and cellular data supporting the findings of this study are included in the article and Supplementary Information. Publicly available sequence, population genetic, and satellite climatology datasets were obtained from NCBI GenBank, gnomAD (v4.1.1), the 1000 Genomes Project and Human Genome Diversity Project (HGDP), NASA CERES (SYN1deg), and NASA MODIS (MCD43C3 Collection 6.1). Custom Python and R code used for the deterministic thermodynamic modeling, geospatial radiometric modeling, and PheWAS regression analyses will be deposited in a public repository (GitHub/Zenodo) upon publication. Individual-level whole-genome sequencing and electronic health record data from the Vanderbilt University Medical Center BioVU biobank are controlled-access human clinical data protected under patient privacy regulations (IRB #240655) and cannot be shared publicly; summary-level association statistics and phecode definitions are provided in the article and Table S3, and qualified researchers may request access through standard VUMC BioVU data-use agreements.

## Supplementary Materials for

### Supplementary Methods

#### 1. Thermodynamic Modeling of Opsin Kinetics

To evaluate the safety limits of the visual cycle, we constructed a hybrid deterministic kinetic model of retinal accumulation. We modeled the rod photoreceptor not as a signal detector, but as a biochemical compartment subject to mass-action kinetics. The model simulates the accumulation of toxic all-trans-retinal (atRAL) during continuous photic exposure. The core novelty of this framework is treating the opsin apo-protein not as an empty vessel, but as a buffer (capacitor) that sequesters free atRAL based on its thermal stability.

The system is defined by three coupled processes:

1. Inflow: Photic activation of Rhodopsin
2. Clearance: Metabolic removal of atRAL by retinol dehydrogenase (RDH)
3. Buffering: Reversible binding of free atRAL to the opsin protein

We used mass balance ODEs to estimate accumulation of total retinoid over minutes, and mass action to partition total retinoid into free vs. opsin-bound states, assuming a quasi-steady state (opsin binding rate>>>accumulation rate). The system was modeled as a competitive equilibrium between opsin-retinal binding (capacitance) and enzymatic clearance (metabolism). The concentration of free all-*trans* retinal (atRAL) was calculated as a function of ecological load (*L*), opsin stability (*K_d_*), and time (*t*).

Using the mass balance equation, accumulation of atRAL is determined by:

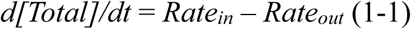

Total atRAL accumulation after *t* minutes is calculated from the integral of the Mass Balance Equation under saturation conditions. To evaluate the opsin capacitor under the most stringent, conservative conditions possible, *Rate_in_* and *Rate_out_* are treated as constants over the integration period (e.g., step functions). By forcing the model to artificially clear all atRAL before tracking accumulation, we intentionally handicap the simulation in favor of cell survival, ensuring that any capacitor failure observed in our results is driven by profound thermodynamic overload rather than simulated kinetic artifacts.

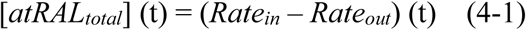

**Clearance**: We define ***Rate_out_*** as clearance of atRAL by retinal dehydrogenases (RDH). Clearance is modeled as a saturated enzymatic process operating at an apparent maximal velocity (*V_max, app_*). While the theoretical *in vitro* catalytic capacity of isolated RDH is massive—mathematically yielding an uninhibited velocity of approximately 840 uM/min—this enzyme operates via a bi-substrate mechanism strictly dependent on the local concentration of the NADPH cofactor^241^. Under intense photostress, the initial resting pool of NADPH is rapidly depleted. Because localized cofactor regeneration via the pentose phosphate pathway cannot match the massive photobleaching influx^242^, the apparent maximal velocity decreases according to Michaelis-Menten kinetics. Consequently, the clearance of atRAL ceases to be limited by the RDH enzyme itself and instead becomes strictly bottlenecked by metabolic regeneration of NADPH^48,67,88^, yielding a steady-state in vivo clearance <<840uM/min. This results in a diminished steady-state in vivo clearance ceiling giving rise to acute atRAL accumulation as previously observed^48,67,108,243^. Because empirical *in vivo* measurements cap murine rhodopsin regeneration at approximately 1% per minute (an absolute clearance rate of 30 µM/min), we model a 60 µM/min ceiling—a highly generous upper bound, intentionally overestimating the retina’s continuous metabolic capacity to ensure a conservative stress test. Crucially, recent kinetic profiling demonstrates that RDH selectively acts on free atRAL, not PE-bound adducts, and the condensation of free atRAL with PE to form N-ret-PE occurs at a rate an order of magnitude faster than RHO hydrolysis^88^. Therefore, enzymatic clearance is fundamentally kinetically disadvantaged. If atRAL is released into the membrane unbuffered, lipid-adduct formation will outcompete RDH clearance regardless of the available enzymatic *V_max_*. Consequently, the RHO capacitor is not merely an overflow backup for when the 60uM/min limit is exceeded; it is a strict thermodynamic requirement that continuously meters atRAL release to prevent instantaneous capture by PE.

##### Capacitor (Buffering/sequestration)

Because the condensation of free atRAL with local lipids to form N-ret-PE operates an order of magnitude faster than RHO hydrolysis^88^ we modeled the outer segment as a continuous thermodynamic partition. Because the forward and reverse kinetics of N-ret-PE Schiff base formation are orders of magnitude faster than the macroscopic accumulation of total retinoids (a separation of timescales), the non-protein-bound pool operates under a rapid equilibrium assumption. At steady state, ∼40% of the non-protein-bound atRAL pool exists as N-ret-PE, while 60% remains as free atRAL. The concentration of free atRAL was determined continuously by solving the mass-balance equation incorporating the receptors dissociation constant (*K_d_*):

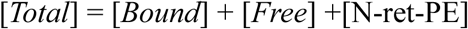

Since 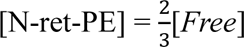, for a single endogenous receptor pool, the system resolves via a quadratic polynomial.

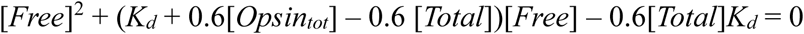

###### Derivation of Thermodynamic Capacitance from Fractional Binding (F_bound_)

Rather than relying on the kinetic decay of the Meta II state (retinal release *t_1/2_*), the thermodynamic capacitance of each opsin variant was derived directly from equilibrium binding assays. The affinity of the opsin apo-protein for all-trans-retinal (atRAL) is defined by its dissociation constant *K_d_*.

In vitro fractional binding (*F_bound_*) was measured via the retinal release fluorescence assay at 20°C in detergent solution. Assays were conducted at fixed total receptor (*R_tot_* = 0.25uM) and total ligand (*L_tot_*= 0.25uM and 2.25uM) concentrations. Because the atRAL ligand is depleted from the free pool as it binds to the opsin capacitor, the standard Michaelis-Menten approximation is invalid. Instead, the exact apparent dissociation constant *K_d, app_* was calculated algebraically by rearranging the quadratic equilibrium binding equation :

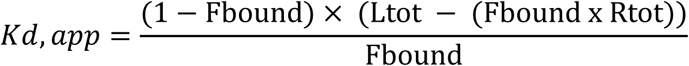

This yields the *in vitro* thermodynamic affinity for each species and allelic variant under controlled experimental conditions. Because in vitro assays were performed in detergent at 20°C, the *K_d, app_* overestimates physiological stability. Opsins are structurally stabilized by artificial detergents and exhibit tighter binding at lower temperatures. To accurately model retinal capacitance in vivo, we scaled the apparent affinity to a native rod outer segment membrane environment at 37°C. We applied a combined biophysical scaling factor based on the Arrhenius/Q10 scaling law and empirical membrane corrections^88^:

- *S_temp_* (thermal scaling): *Q_10_* ≈ 2.5. For 20°C ◊ 37°C, scaling is ≈ 5.3
- *S_mem_* (membrane factor): In vitro detergents stabilize opsins artificially compared to the native membrane. We apply a correction factor of 1.55.

The physiological dissociation constant was calculated as:

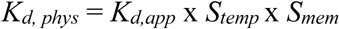

This resulting *K_d, phys_* was used to govern the mass-action sequestration of atRAL in all kinetic simulations, enabling a direct thermodynamic comparison between mammalian RHOs.

#### 2. Derivation of Photic Loads

To evaluate the thermodynamic burden of atRAL clearance on the visual cycle across distinct species, the localized “Ecological Load” (*Rate_in_*) was modeled using precise environmental luminances and morphological retinal amplification (Gain).

The human visual system was utilized to establish the baseline daylight generation rate of atRAL. We derived this baseline using two independent, convergent frameworks:

1. *Bottom-Up Biophysical Constraints* (180 µM/min): Under bright daylight (photopic) conditions, a classical mammalian rod experiences a photoisomerization rate approaching 10^5^ R* per rod per second^244^. Given a standard rod outer segment contains approximately 10^8^ rhodopsin molecules^158^, this yields a maximal fractional bleach rate of 0.1% per second, or 6% per minute. At a standard geometric outer segment opsin concentration of ∼3000 µM, this 6% bleach rate generates a peak photic influx of exactly 180 µM/min of atRAL.
2. *Top-Down In Vivo Densitometry* (120 µM/min): Classical human densitometry establishes that exposure to typical daylight (luminance ∼25,000 cd/m^2^) produces a 50% steady-state rhodopsin bleach^106,245^. At this 50% steady state, the first-order generation of atRAL (*Rate_in_*) must equal the zero-order enzymatic clearance limit (*V_max, app_*), such that the net accumulation rate is zero. Given the conservative clearance bottleneck established above (*V_max, app_* = 60 µM/min), the generation rate is constrained to 60 µM/min. The generation rate is the product of the environmental bleaching constant (*k_bleach_*) and the concentration of available dark-state rhodopsin *Rho_dark_*. Assuming a total rod outer segment rhodopsin concentration of 3000 µM, a 50% steady-state bleach leaves exactly 1500 µM of *Rho_dark_* available for photoisomerization. Solving for the bleaching constant (60 µM/min = *k_bleach_* x 1500) yields *k_bleach_* = 0.04 min^-1^, revealing that daylight consumes exactly 4% of the available rhodopsin pool per minute. Applying this constant to a fully dark-adapted ancestral human (*Rho_dark_* = 3000 µM) experiencing an acute daylight transition yields a maximal raw photic inflow rate *Rate_in,acute_* of exactly 120 µM/min (0.04 x 3000 µM).

Because this steady-state equilibrium dictates that the generation rate cannot exceed the physiological mammalian clearance bottleneck (*V_max,app_*= 60µM/min), the raw environmental influx acting on the available unbleached opsin pool equates to an acute load of 120 µM/min.

Together, these independent methodologies establish a highly constrained human daylight load ranging between 120 and 180 µM/min.

To calculate the specific load experienced by *P. alecto*, we intentionally anchored our model to the conservative lower bound of the human daylight range (120 µM/min) to prevent artificially inflating the bat’s environmental threat. This 120 µM/min baseline was then scaled to reflect chiropteran biology and roosting ecology. Megabats of the genus *Pteropus* do not echolocate and rely entirely on vision and olfaction for navigation and foraging^246^. Morphological investigations of *Pteropus* retinal ganglion cells demonstrate striking convergence with the highly sensitive retina of the cat, featuring expansive dendritic fields designed for maximum spatial summation^146^. Furthermore, electrophysiological profiling of the *Pteropodidae* family confirms extreme absolute visual thresholds—ranging from -6.30 to -6.37 log cd/m^2^ s in related species like *C. sphinx* and *R. leschenaultii*— compared to microbats with degraded visual systems^147^. This massive photon-pooling architecture is further exacerbated by the megabat’s unique retinal anatomy. The *Pteropus* retina is unusually thick (270–350µM) yet strictly avascular, relying instead on spike-like ‘choroidal papillae’ to deliver oxygen^146^. Because it lacks an intraretinal vascular network that would otherwise absorb or scatter incoming light, the *Pteropus* retina receives an unimpeded influx of photons, maximizing sensitivity but drastically increasing the photic load and atRAL generation when exposed to daylight. This extreme reliance on visual gain is mirrored in the central nervous system; *Pteropodidae* exhibit a massive expansion of the superior colliculus relative to the inferior colliculus (a 3:1 SC/IC volume ratio, compared to 1:7 in echolocating insectivorous bats), indicating profound neural dedication to processing visual input^147^. To account for this massive nocturnal photon-pooling architecture, the morphological gain for *P. alecto* was set to 5.0. Unlike cave-dwelling species, *Pteropus* roost in open tree canopies exposed to bright ambient daylight. Using standard radiometric models, the diffuse daytime illuminance of a shaded canopy approaches 50,000 lux^247^, corresponding to an environmental luminance of approximately 12,500 cd/m^2^. The *P. alecto* ecological load was therefore calculated by multiplying the conservative human baseline (120 µM/min) by the *Pteropus* gain (5.0) and the specific ratio of their environmental luminances (12,500 / 25,000 cd/m^2^), yielding a calculated ecological load of 300 µM/min.

#### 3. Geospatial Modeling of Ancestral Phototoxicity

While general species loads were modeled using discrete photic niches, the specific historical ecological loads (*I_in_*) for the human genetic cohorts (gnomAD) were derived empirically from continuous satellite telemetry to map the theoretical *uM/min* load across the global coordinate system. To quantify the ancestral mid-winter radiometric load, a 25-year climatological baseline (February 2001 – February 2025) was derived from NASA satellite observations. Downwelling solar irradiance was extracted from the Clouds and the Earth’s Radiant Energy System (CERES) SYN1deg product^248^. To account for realistic atmospheric attenuation and regional cloud cover variance natively, the adjusted all-sky surface shortwave direct and diffuse fluxes were summed to calculate the true Global Horizontal Irradiance (GHI) hitting the Earth’s surface. While irradiance is a 25-year average, the surface reflectance utilizes a representative modern winter snapshot (February 2024). Global surface reflectance was mapped using the Moderate Resolution Imaging Spectroradiometer (MODIS) MCD43C3 Collection 6.1 Climate Modeling Grid^249^. The Shortwave White-Sky Albedo (ρ) variable was isolated to represent bi-hemispherical reflectance under diffuse illumination, providing a continuous, empirical metric for regional snowpack and ground reflectivity. NASA scale factors (0.001) were applied to the raw HDF4 data to restore standard physical units.

Geospatial processing was conducted using the xarray and pyhdf libraries in Python. To correct for coordinate reference system mismatches, the CERES spatial grid was mathematically rolled from a 0–360 degree to a - 180–180 degree longitudinal domain to align with the MODIS reference frame. The combined matrices were subsequently downsampled via linear interpolation to a unified 0.25-degree resolution grid. This generated a matrix of approximately one million discrete geographic pixels, ensuring high-fidelity topological mapping while optimizing computational memory allocation for the biophysical merge.

This raw planetary radiometry was converted into an objective biological load representing the kinetic rate of all-trans-retinal (atRAL) generation. The vertical ground-reflected irradiance hitting the cornea (*E_v_*) was calculated utilizing the standard geometric view factor for a vertical surface^250^.

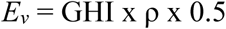

To computationally generate the Retinal Threat Matrix across a 1-million-pixel global grid without requiring recursive, pixel-by-pixel loop integrations of independent physical constants, we derived a lumped geometric-optical scalar. This allows a conversion from raw vertical irradiance (*E_v_* in W/m^2^) to the intracellular atRAL generation rate (µM/min). First, the environmental photon flux entering the eye (Φ*_env_*) is calculated from the vertical irradiance (*E_v_*) using a standard radiometric-to-retinal optical conversion constant (*K_opt_*):

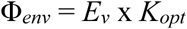

The constant *K_opt_* (0.946) is the mathematical product of the Planck-Einstein relation at the rhodopsin peak absorption wavelength (λ_max_ = 494nm), the light-adapted pupillary aperture area (3.14 mm^2^ corresponding to a 2.0 mm diameter^251^), ocular media transmittance (45%^43^), and the quantum capture cross-section of a mammalian rod outer segment (0.5 um^2^). By standardizing the aperture to the human physiological minimum (2.0 mm), we explicitly model the retinal load *after* the pupillary light reflex (PLR) has been fully exhausted. While avoiding the confounding variables of bespoke facial anatomy or behavioral squinting across different human populations, this standardized mechanical ceiling isolates the raw environmental thermodynamic pressure, revealing the specific magnitude of phototoxicity that the opsin biochemical capacitor evolved to buffer. This derived photon flux is then multiplied by a biophysical volumetric scaling constant *S_scale_* to convert raw photon counts into the specific accumulation rate of all-*trans*-retinal within the rod outer segment volume

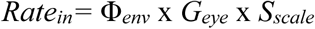

We mathematically derived *S_scale_* by anchoring the equation to the physiological reality established above. Under bright clear-sky daylight (yielding an *E_v_* of ∼400W/m^2^) and an intermediate dynamic pupillary aperture (a 6.25x multiplier), in vivo densitometry establishes a biological accumulation rate of ∼120 µM/min. Solving the equation (120 = 400 x 0.946 x *S_scale_* x 6.25) yields an *S_scale_* constant of **0.05**. By substituting this optical conversion step into the biological generation equation (where *G_eye_* = 1.0 for humans), we collapse the pipeline into a single, highly efficient computational scalar used in the global mapping script:

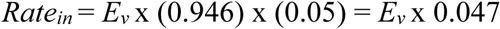

This combined multiplier (0.047) allowed for the direct, continuous transformation of the NASA W/m^2^ satellite matrices into the final biophysical Retinal Threat Matrix (µM/min) mapped in Figure 4C, ensuring exact parity between the environmental physics and the mass-action ODEs.

## Supplementary Figures

**Figure S1.**
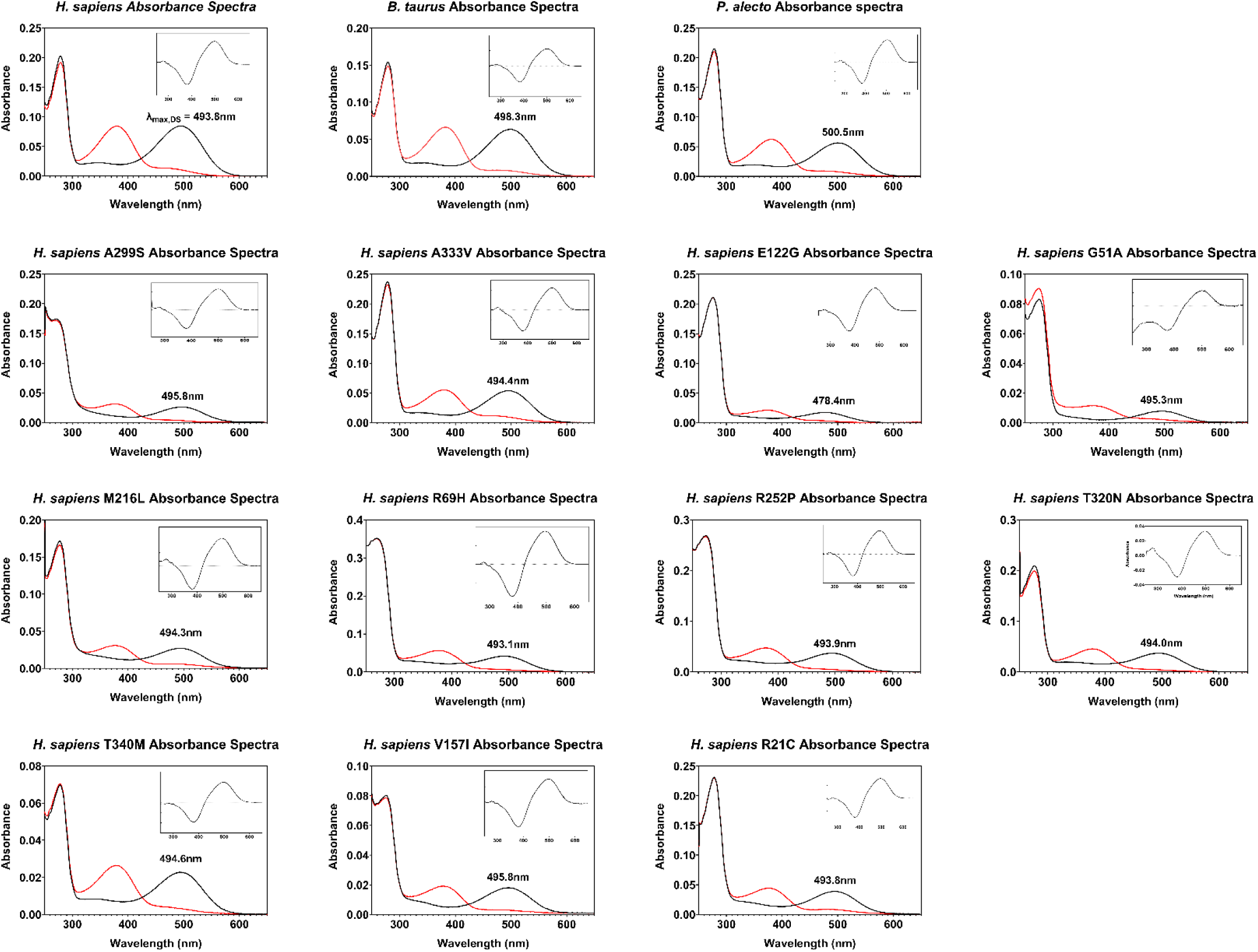
- UV-Vis absorbance spectra of purified RHO. Absorbance spectra (250–650 nm) were recorded for WT from *H. sapiens*, *B. taurus*, and *P. alecto* along with human alleles. Solid black lines indicate the dark state (DS) holo-protein, displaying the characteristic total protein peak at 280 nm and dark state peak of the 11CR bound pigment. Red lines represent the spectra following photobleaching, showing the formation of the Meta II intermediate. Insets display the difference spectra (dark-state minus bleached) used to isolate the chromophore absorbance. To precisely determine the λ_max_ values indicated in each panel, the DS spectra were curve-fit using a template for RHO absorbance as described by Govardovskii et al. (2000).

**Figure S2.**
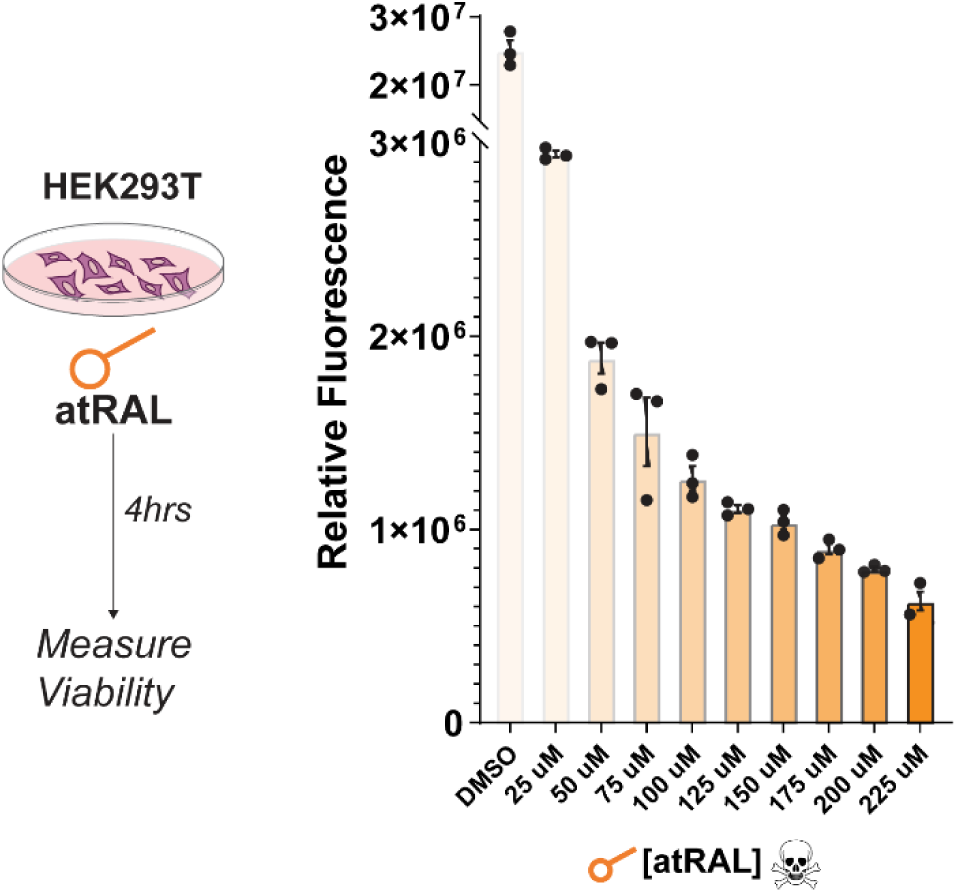
Cytotoxicity of atRAL in HEK293T cells. To assess the cyto toxic impact of exogenous retinal, HEK293T cells were transfected with an empty P1D4-hrGFPII vector and allowed to recover for 48 hours. Cells were subsequently treated with varying concentrations of atRAL (0–225µM)for 4 hours. Cell viability was quantified using the CellTiter-Blue (Promega) resazurin-based assay, with fluorescence measured on a BMG Labtech plate reader. Data are presented as Relative Fluorescence Units (RFU), where a dose-dependent decrease in fluorescence indicates a reduction in metabolic activity and cell viability corresponding to increasing atRAL concentrations. Individual data points represent technical replicates (n = 3) and error bars indicate ± SEM.

**Figure S3.**
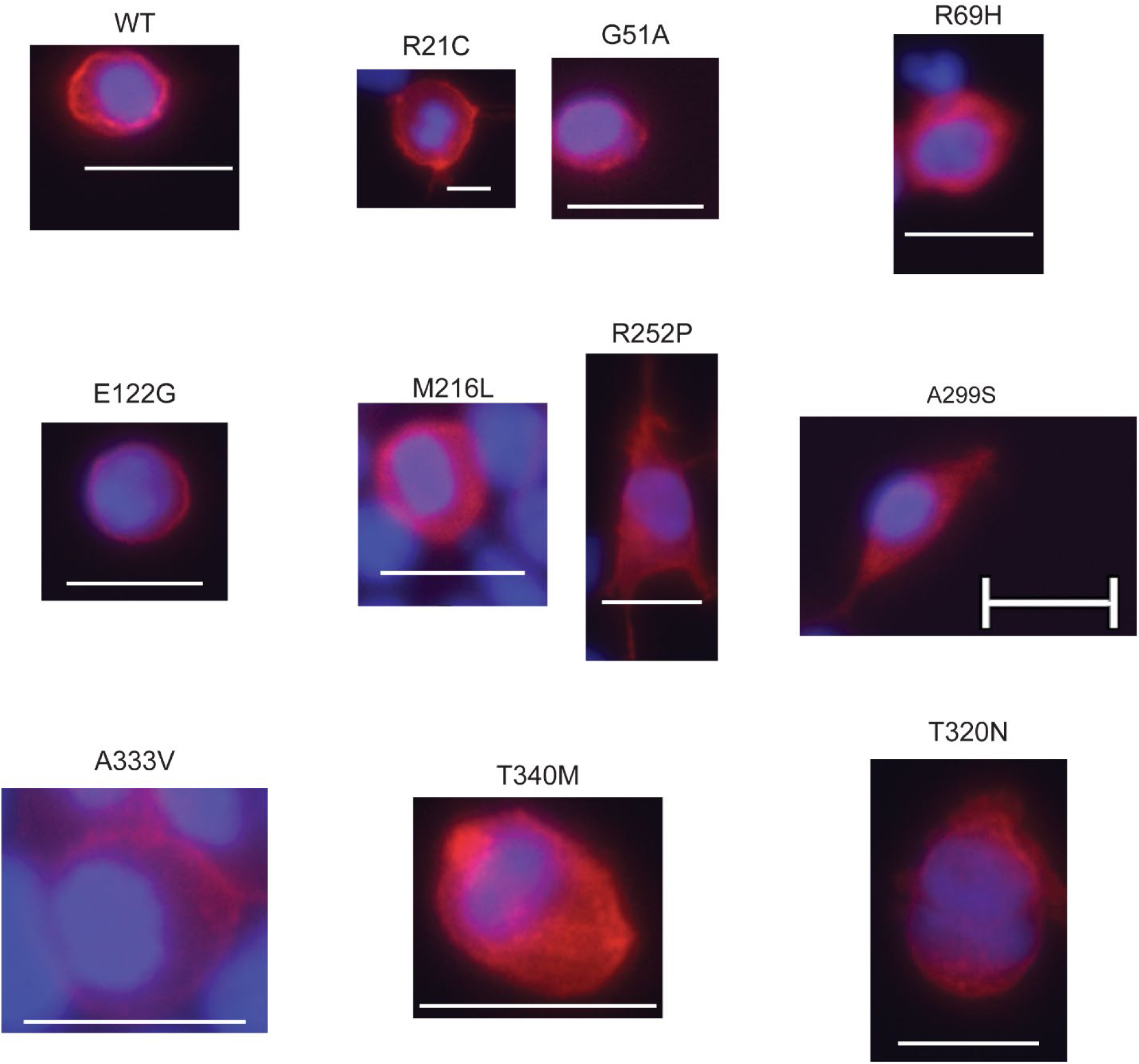
- Subcellular localization of Human RHO variants in HEK293T cells. Immunocytochemistry was performed to evaluate the expression and trafficking of wild-type (WT) and mutant Human RHO. HEK293T cells were transiently transfected with the indicated RHO variants and imaged via fluorescence microscopy. RHO (red) was visualized using anti-1D4 antibody (Abcam) followed by Alexa-488 secondary and nuclei were counterstained with DAPI (blue). WT RHO and alleles exhibit characteristic plasma membrane localization.

**Figure S4.**
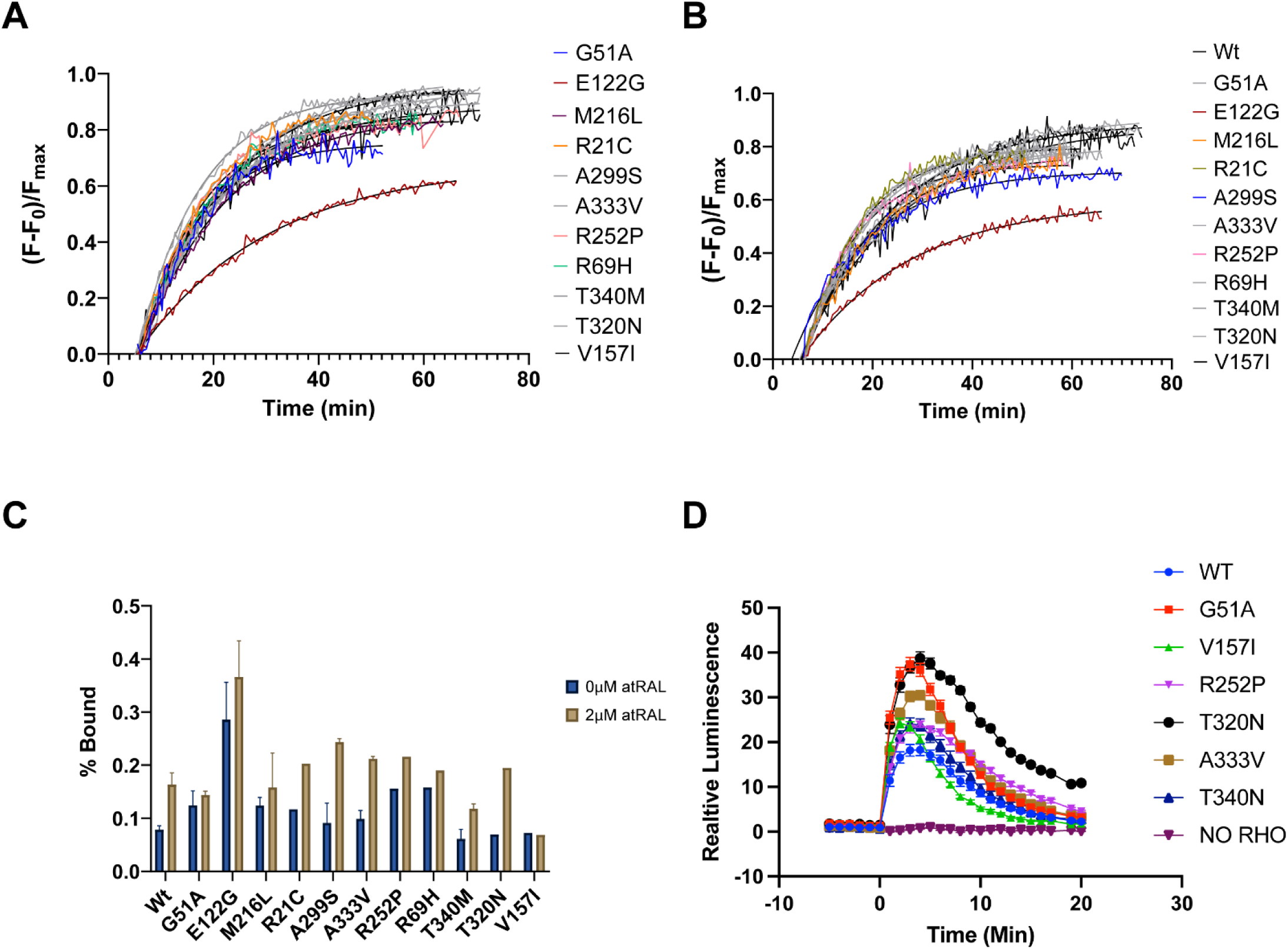
- Functional characterization of Human rhodopsin alleles of interest. **(A-B)** Retinal release assay of human RHO alleles with 0 (A) or 2 µM (B) exogenous atRAL. Fluorescence data was baselined and normalized to max fluorescence to compare MII decay and F_bound_ values. **(C)** Calculated fraction bound retinal at steady at both 0 and 2µM exogenous atRAL with error bars indicating SEM. **(D)** Glosensor assay monitoring signaling of RHO after light bleach at time 0. C and D are normalized to protein expression.

**Figure S5.**
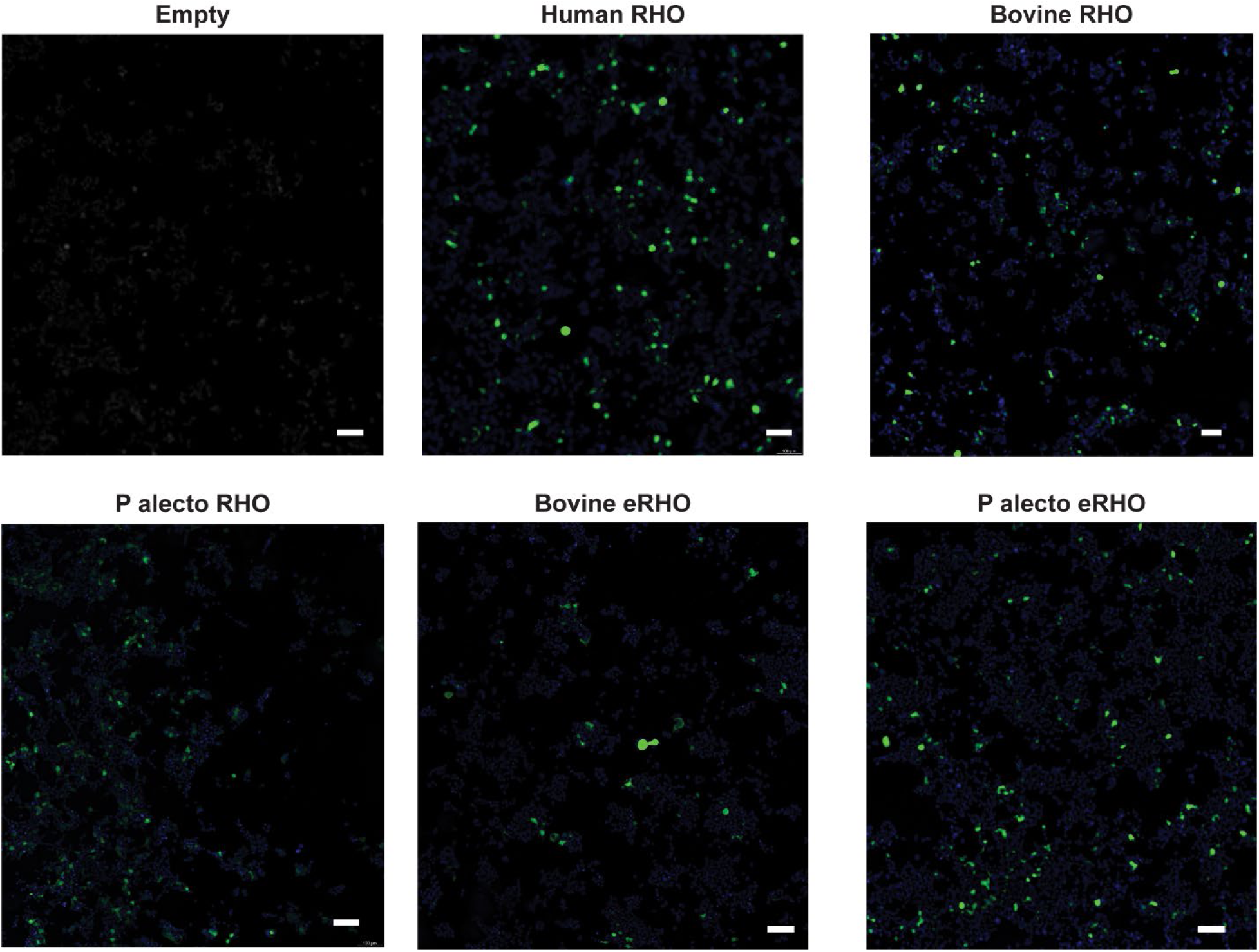
– Immunocytochemistry of Rhodopsin in HEK293T cells. Cells were transfected with empty vector, or WT RHO orthologs, or engineered RHOs. Cells were stained with anti-1D4 primary antibody, followed by Alexa-488 secondary, and imaged under widefield with computational clearing (Leica, MICA). Scale bar = 100 µm.

**Fig. S6.**
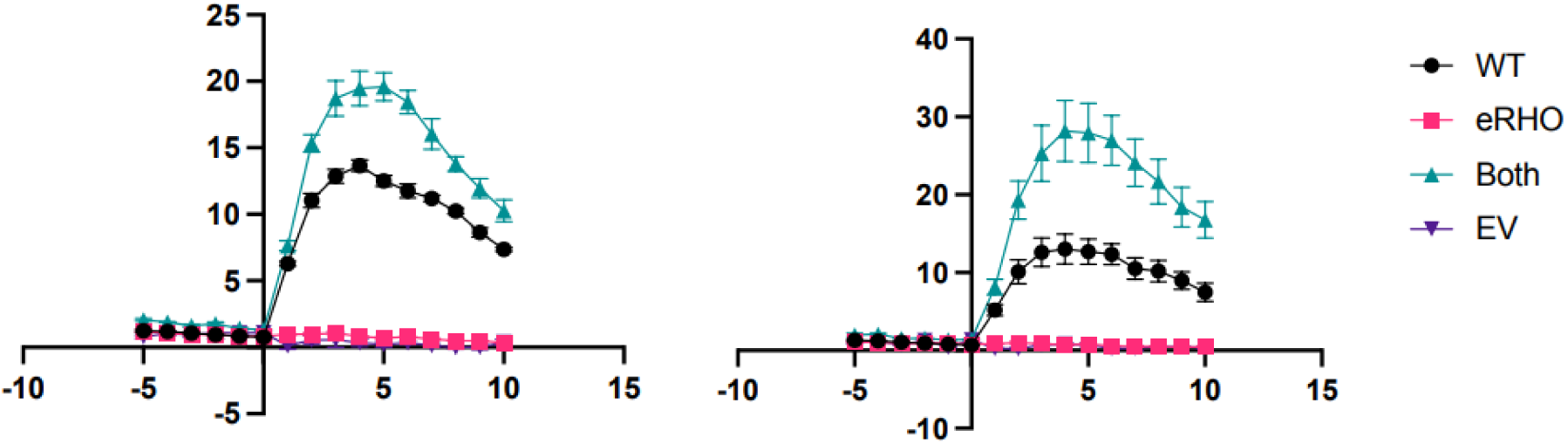
eRHO enhances WT RHO signalling. Glosensor assay of HEK293T cells following various transfections. Left, 1 µM exogenous atRAL was added prior to light bleach at time 0. Right, 100 nM.. Y-axis is luminescence normalized to protein expression. X-axis denotes time in minutes.

**Fig S7.**
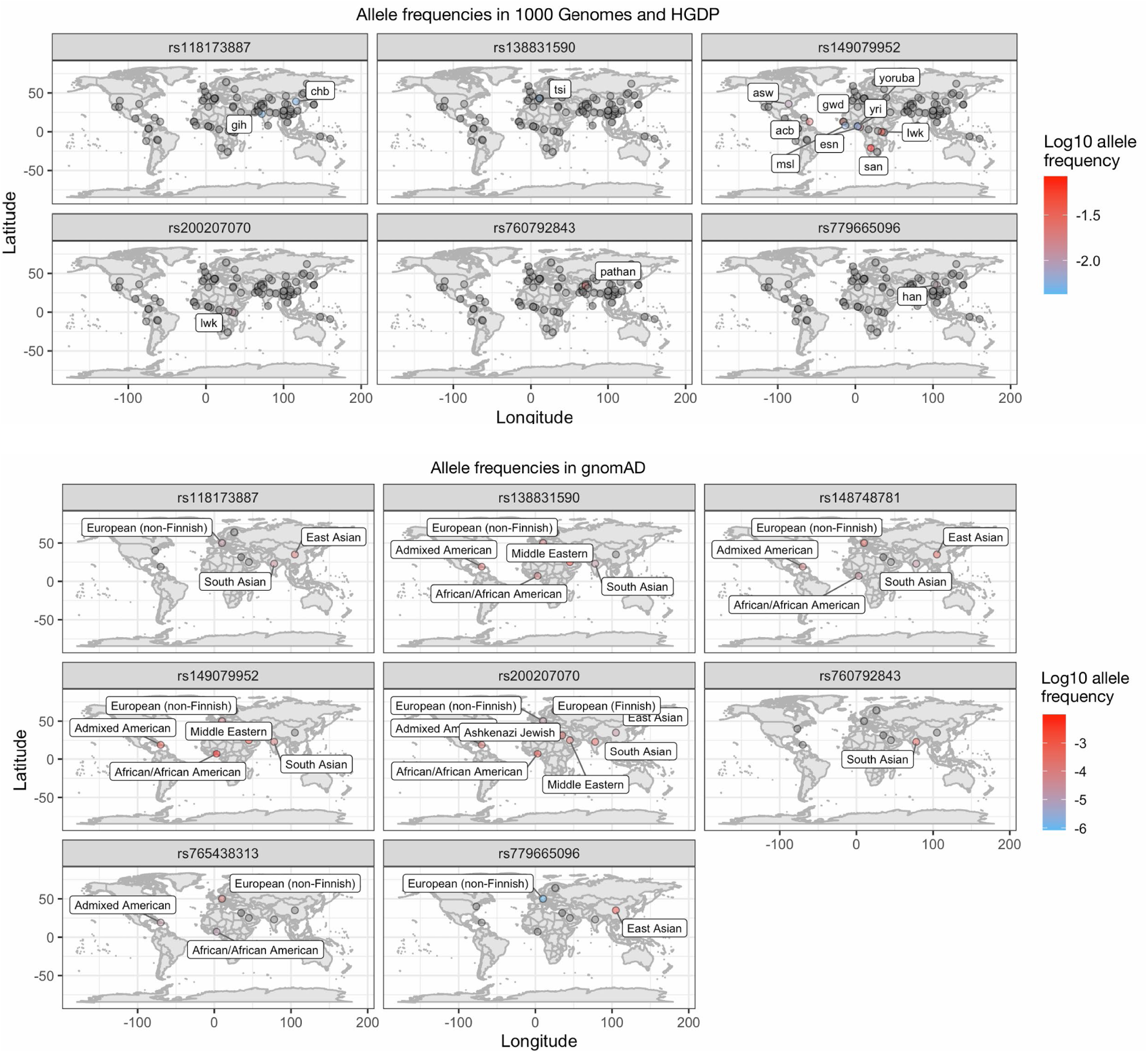
Allele frequencies in 1KG + HGDP with gnomAD v4.1. Not all variants that are in gnomAD appear in 1000 Genomes, hence different numbers of plots. In all plots, grey points indicate the allele is not present in that location, and only populations with a non-zero allele frequency are labelled.

**Fig. S8.**
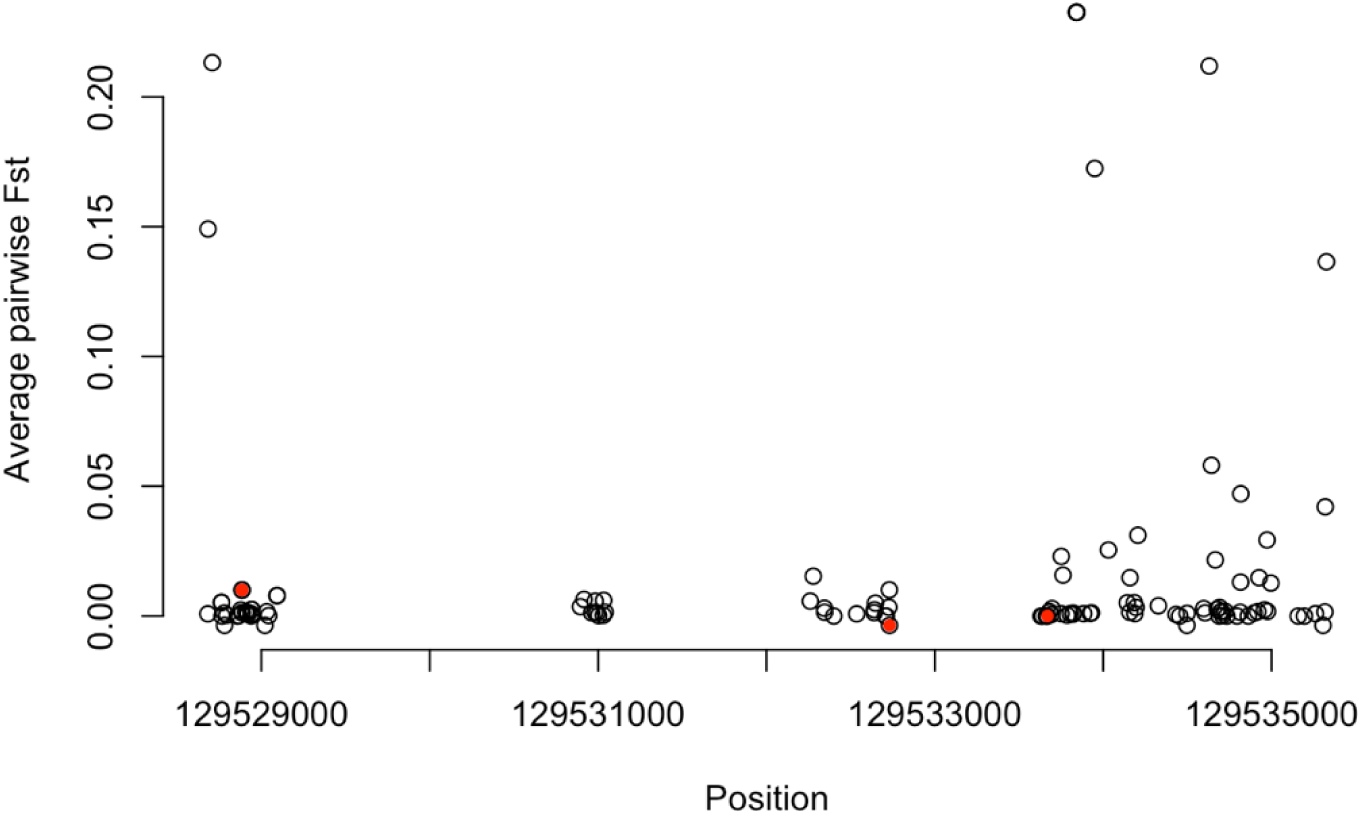
Pairwise Fst for all SNPs in RHO exons (n=114). Variants functionally characterized in this study are red.

**Table S1.**
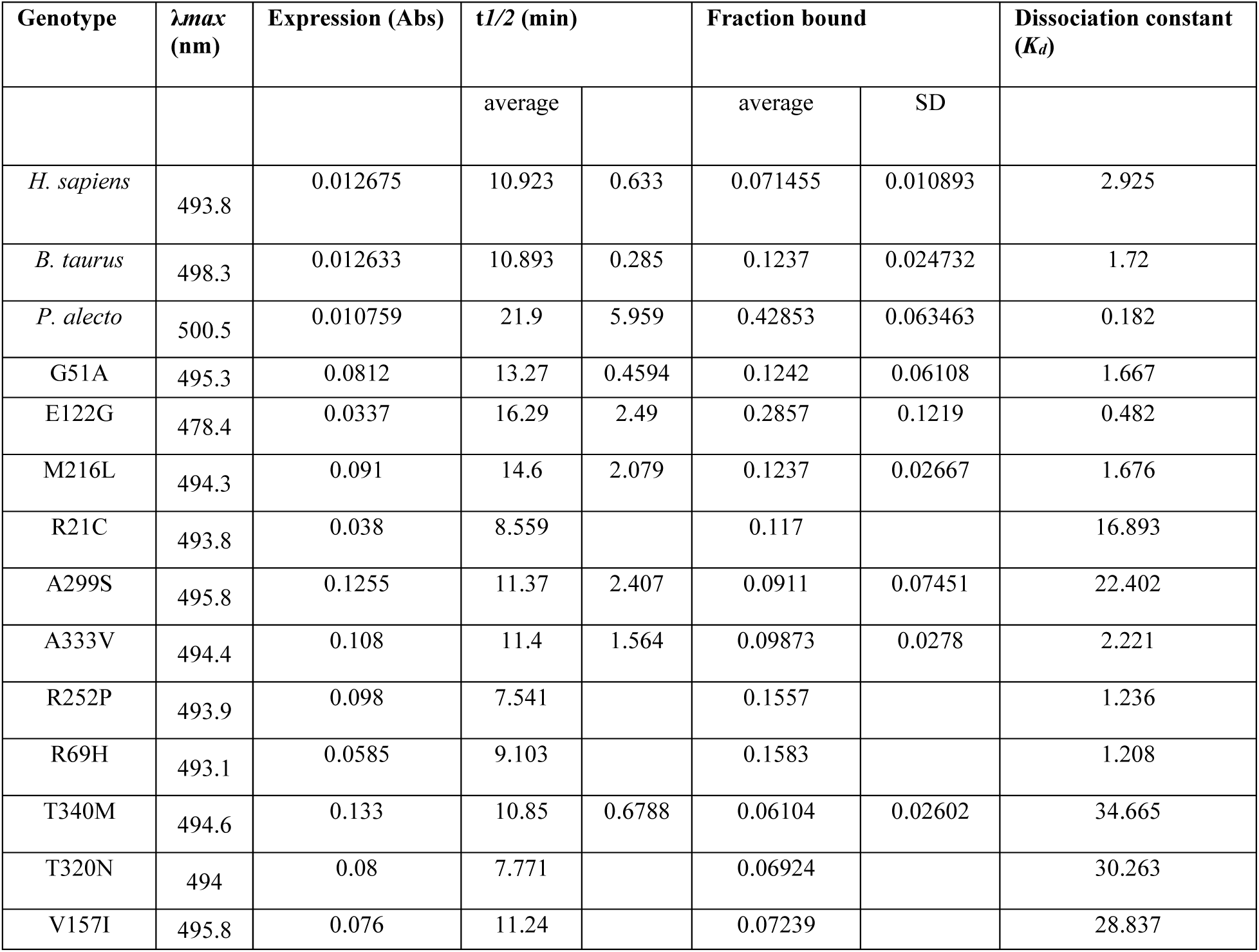
Biochemical phenotypes of RHO from different species and different human RHO mutations. All values were determined at 20°C with no exogenous atRAL.

**Table S2.**
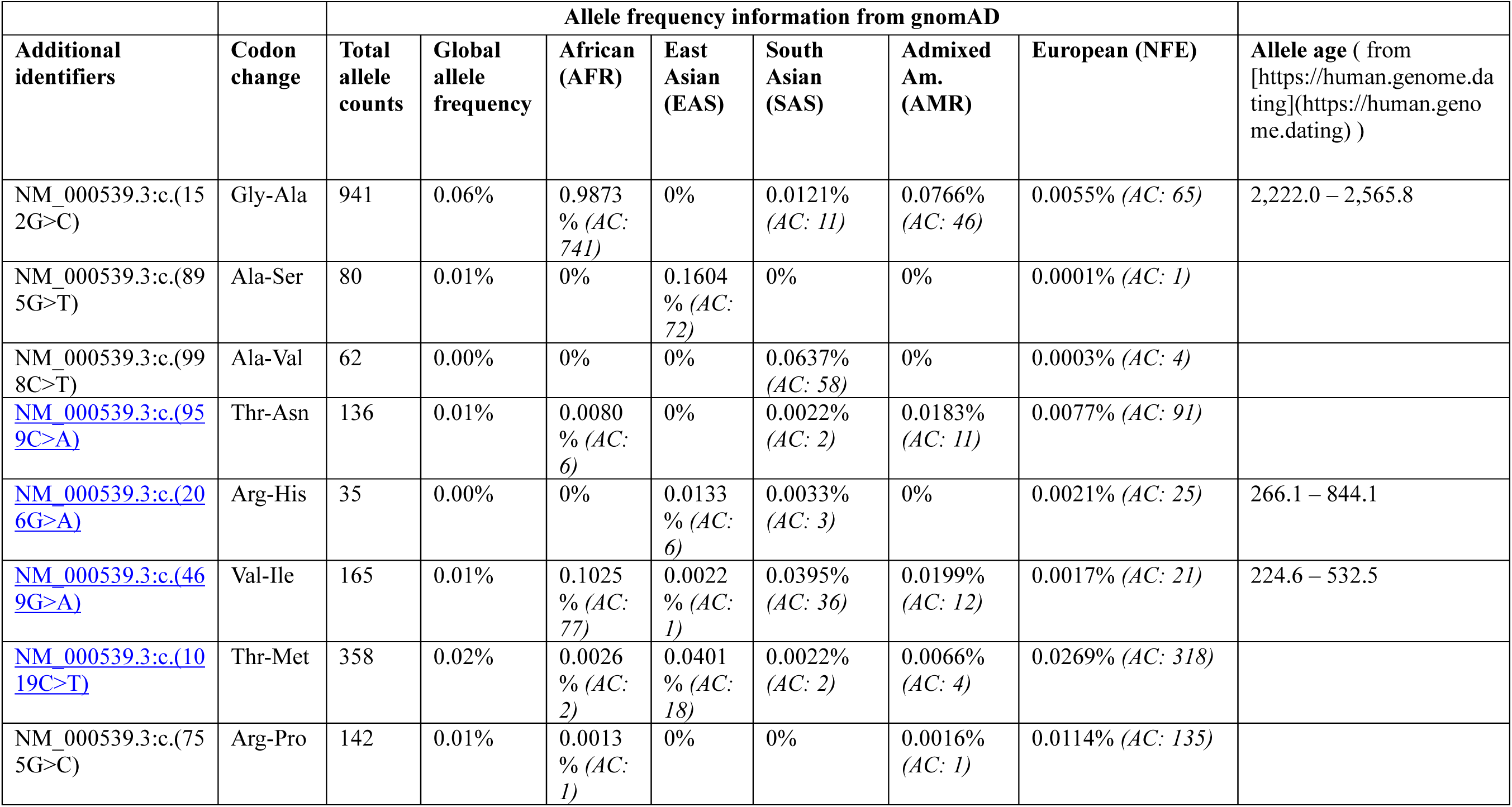
Human RHO alleles information from Gnomad and allele age information.

**Table S3.**
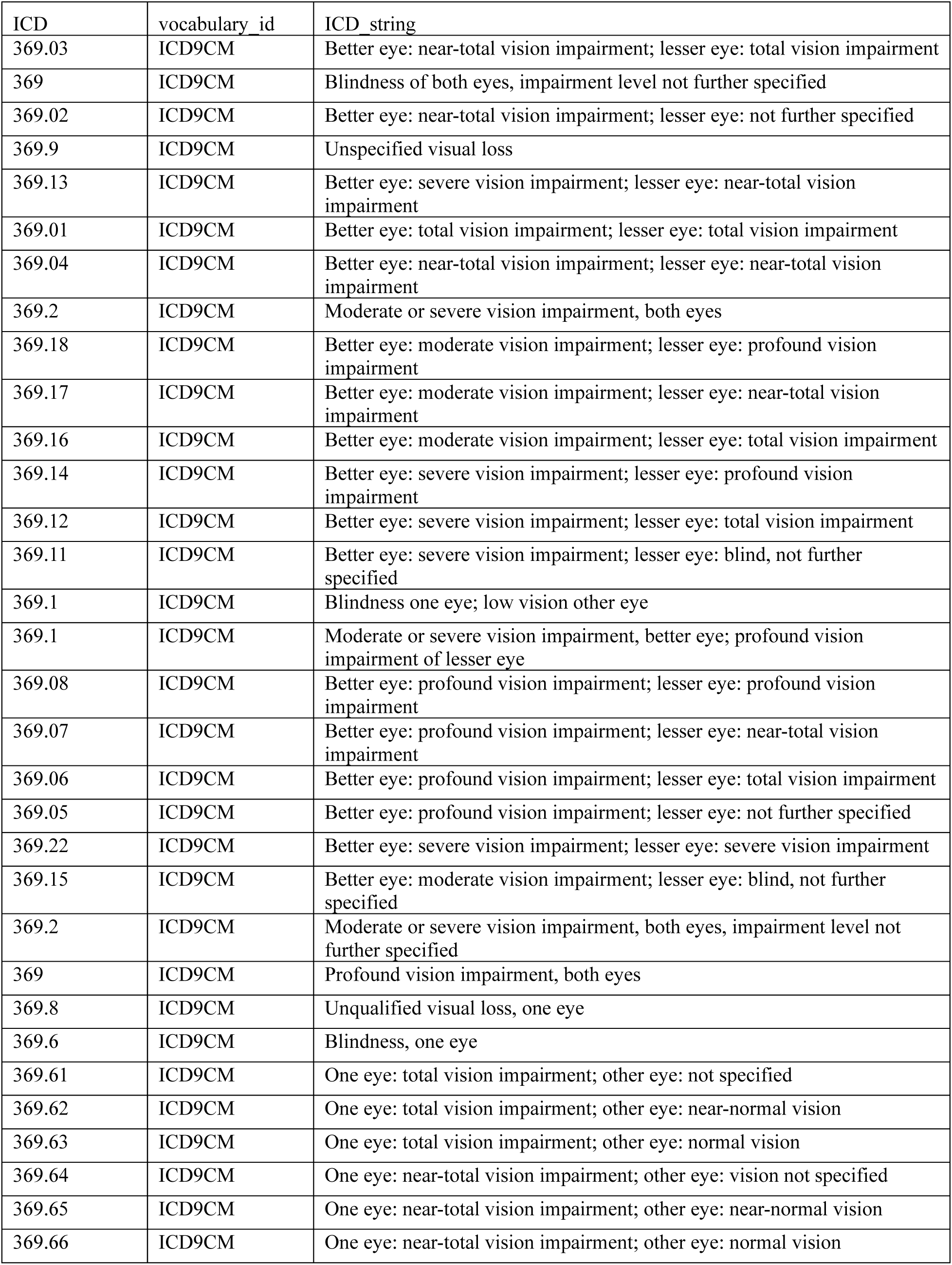
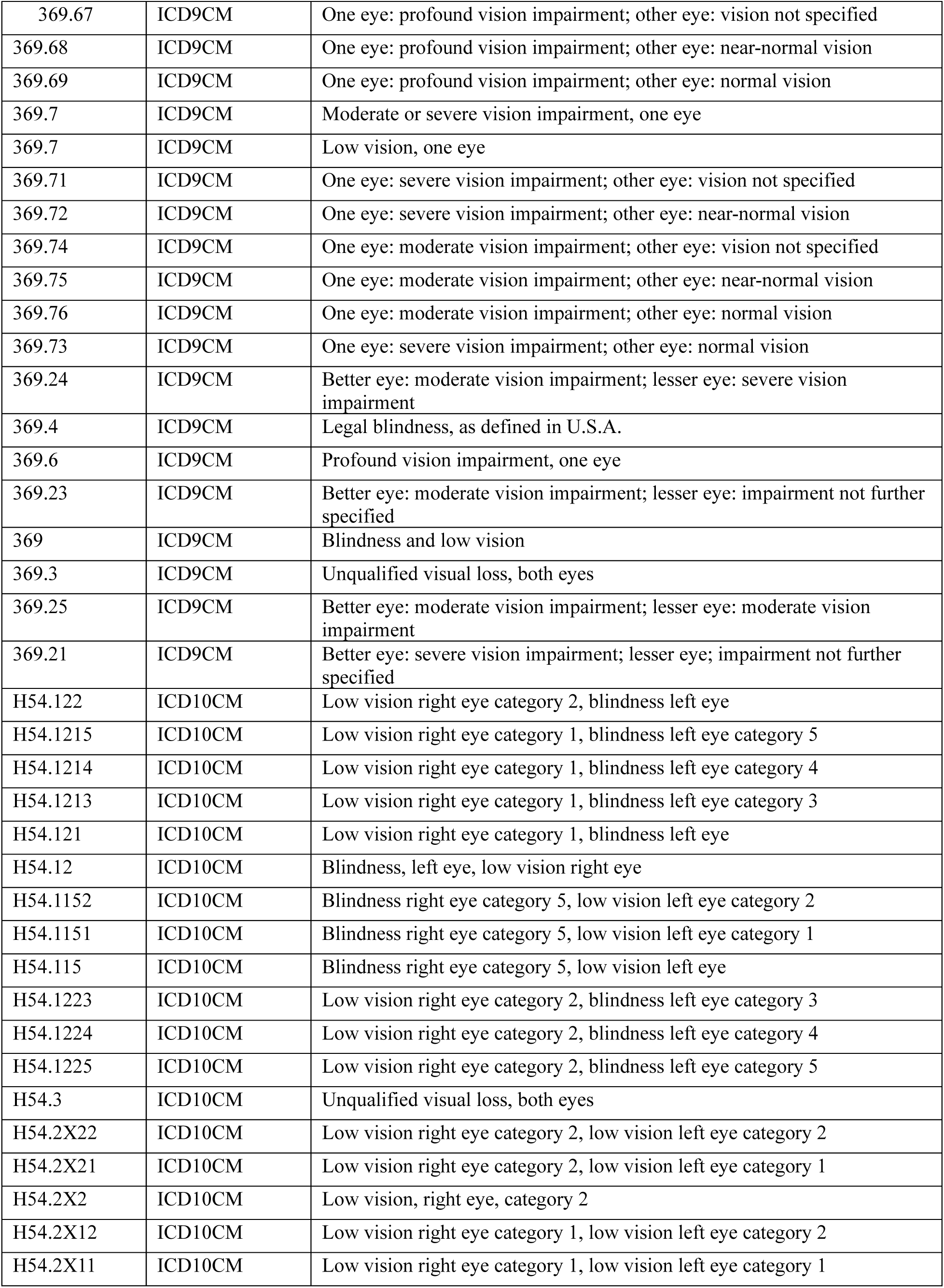
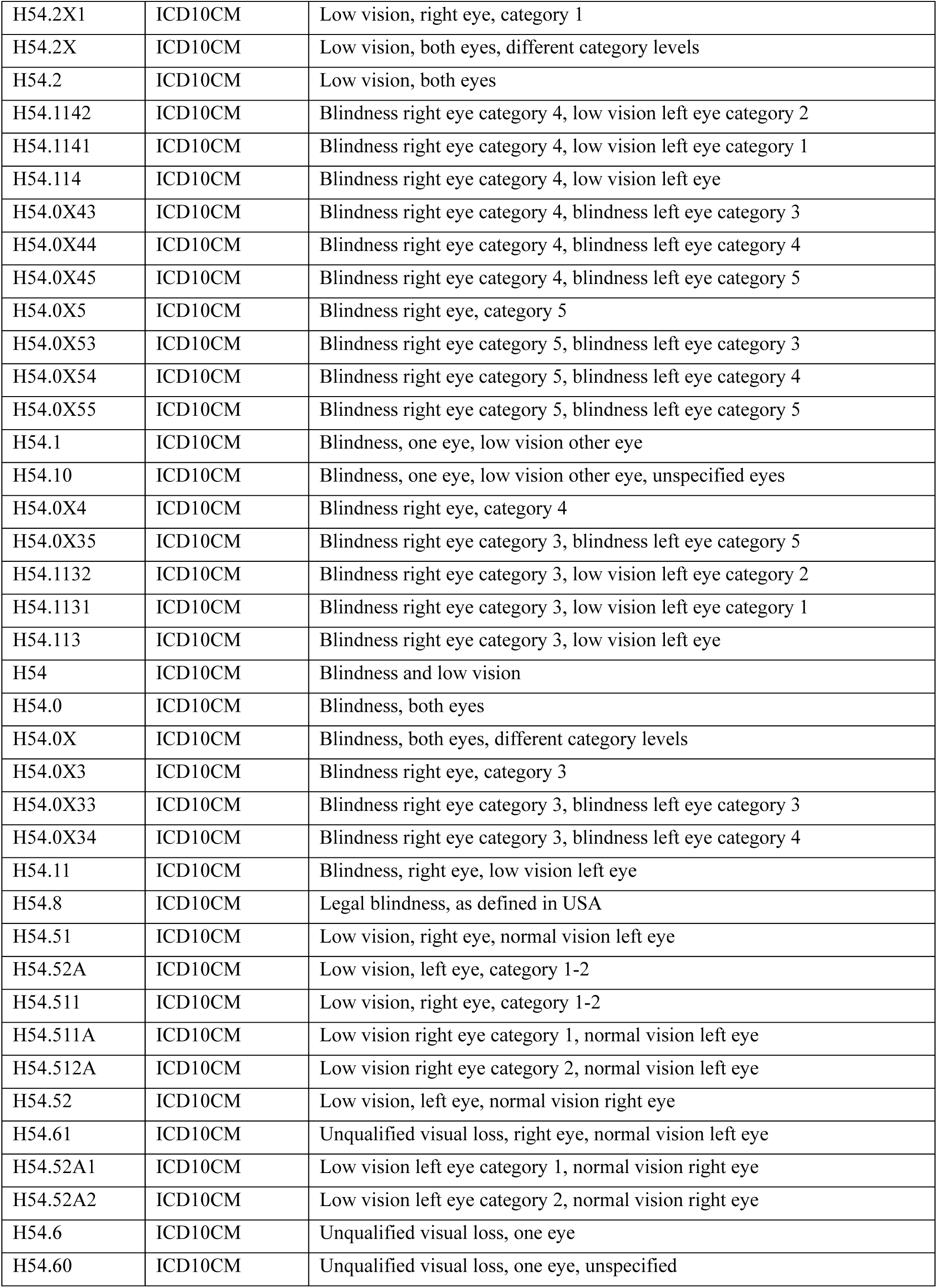
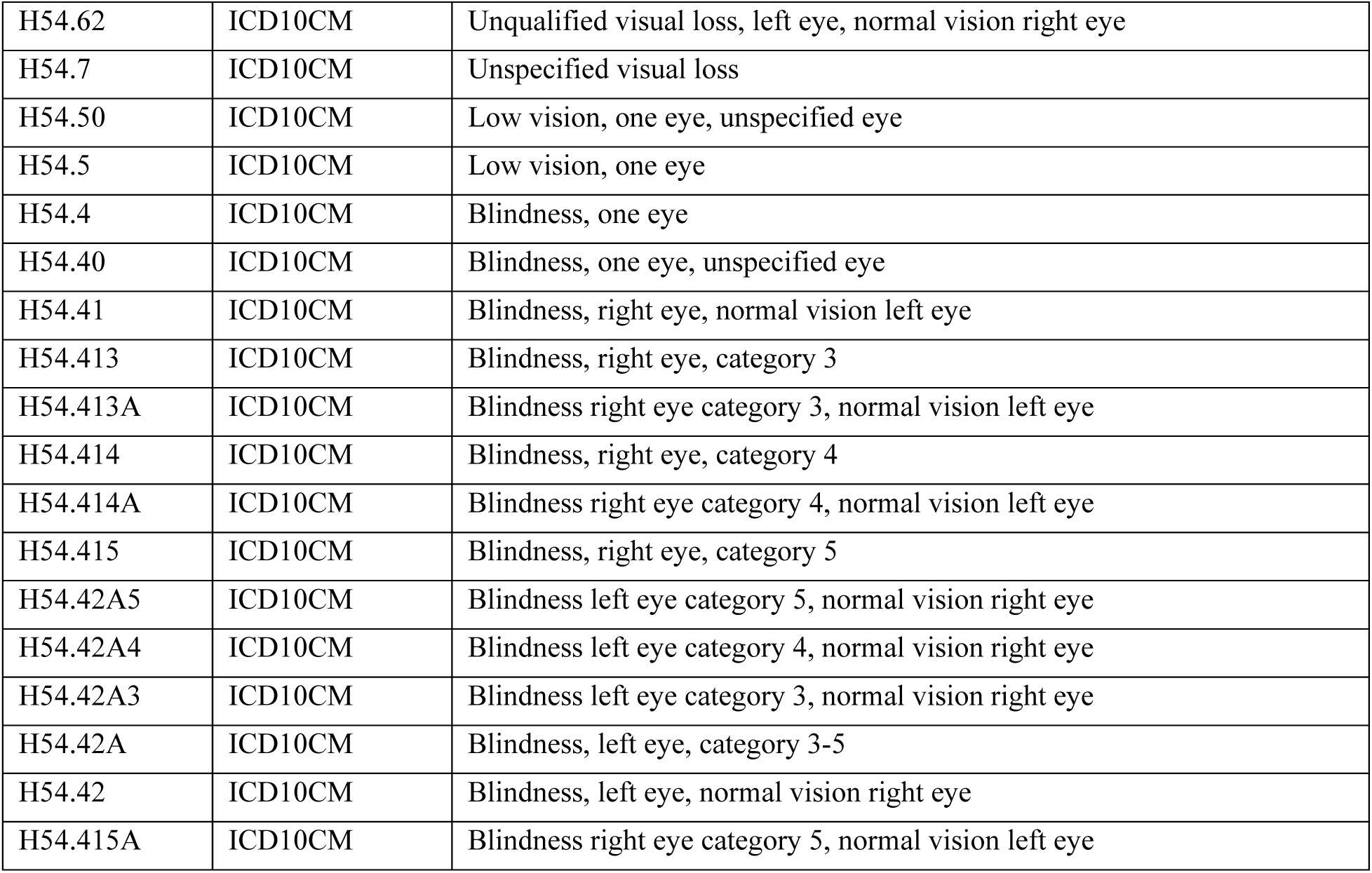
Electronic Health Record phecodes for “Blindness and low vision”.

