## Supplementary Materials for "Evolutionary tuning of rhodopsin-mediated light protection"

$$[Total] = [Bound] + [Free] + [\text{N-ret-PE}]$$

Since $[\text{N-ret-PE}] = \frac{2}{3}[Free]$, for a single endogenous receptor pool, the system resolves via a quadratic polynomial.

$$[Free]^2 + (K_d + 0.6[Opsin_{tot}] - 0.6\,[Total])[Free] - 0.6[Total]K_d = 0$$

*Derivation of Thermodynamic Capacitance from Fractional Binding* (**$F_{bound}$**):

Rather than relying on the kinetic decay of the Meta II state (retinal release $t_{1/2}$), the thermodynamic capacitance of each opsin variant was derived directly from equilibrium binding assays. The affinity of the opsin apo-protein for all-trans-retinal (atRAL) is defined by its dissociation constant $K_d$.

In vitro fractional binding ($F_{bound}$) was measured via the retinal release fluorescence assay at 20°C in detergent solution. Assays were conducted at fixed total receptor ($R_{tot}$ = 0.25uM) and total ligand ($L_{tot}$= 0.25uM and 2.25uM) concentrations. Because the atRAL ligand is depleted from the free pool as it binds to the opsin capacitor, the standard Michaelis-Menten approximation is invalid. Instead, the exact apparent dissociation constant $K_{d,\,app}$ was calculated algebraically by rearranging the quadratic equilibrium binding equation :

$$Kd, app = \frac{(1 - \text{Fbound}) \times \ (\text{Ltot} \ - \ (\text{Fbound x Rtot}))}{\text{Fbound}}$$

This yields the *in vitro* thermodynamic affinity for each species and allelic variant under controlled experimental conditions. Because in vitro assays were performed in detergent at 20°C, the $K_{d,\,app}$ overestimates physiological stability. Opsins are structurally stabilized by artificial detergents and exhibit tighter binding at lower temperatures. To accurately model retinal capacitance in vivo, we scaled the apparent affinity to a native rod outer segment membrane environment at 37°C. We applied a combined biophysical scaling factor based on the Arrhenius/Q10 scaling law and empirical membrane corrections[88]:

- $S_{temp}$ (thermal scaling): $Q_{10} \approx 2.5$. For 20°C → 37°C, scaling is ≈ 5.3
- $S_{mem}$ (membrane factor): In vitro detergents stabilize opsins artificially compared to the native membrane. We apply a correction factor of 1.55.

The physiological dissociation constant was calculated as:

$$K_{d,\,phys} = K_{d,app} \times S_{temp} \times S_{mem}$$

This resulting $K_{d,\,phys}$ was used to govern the mass-action sequestration of atRAL in all kinetic simulations, enabling a direct thermodynamic comparison between mammalian RHOs.

### Supplementary Figures

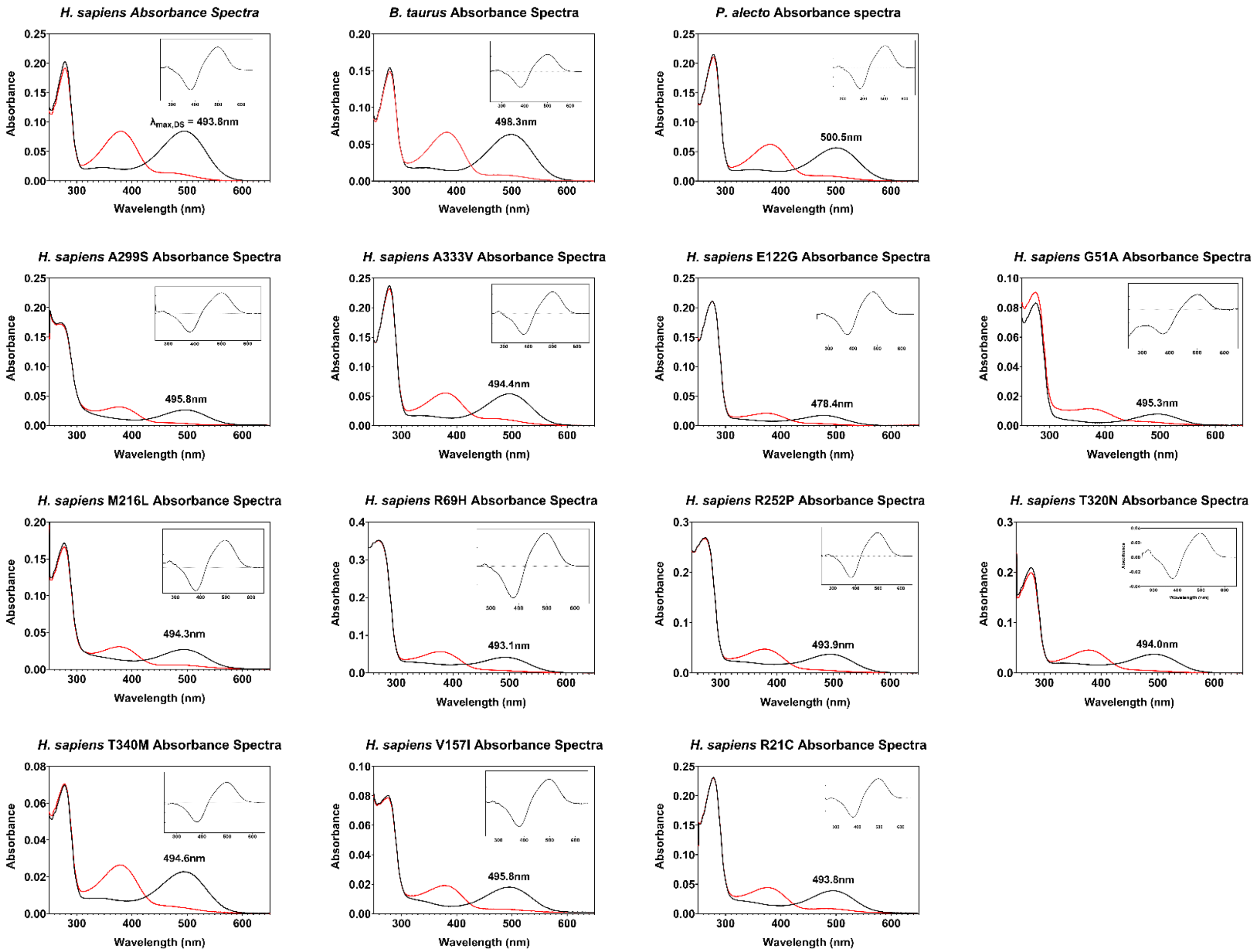


**Figure S1- UV-Vis absorbance spectra of purified RHO.** Absorbance spectra (250–650 nm) were recorded for WT from *H. sapiens*, *B. taurus*, and *P. alecto* along with human alleles. Solid black lines indicate the dark state (DS) holo-protein, displaying the characteristic total protein peak at 280 nm and dark state peak of the 11CR bound pigment. Red lines represent the spectra following photobleaching, showing the formation of the Meta II intermediate. Insets display the difference spectra (dark-state minus bleached) used to isolate the chromophore absorbance. To precisely determine the $\lambda_{max}$ values indicated in each panel, the DS spectra were curve-fit using a template for RHO absorbance as described by Govardovskii et al. (2000).

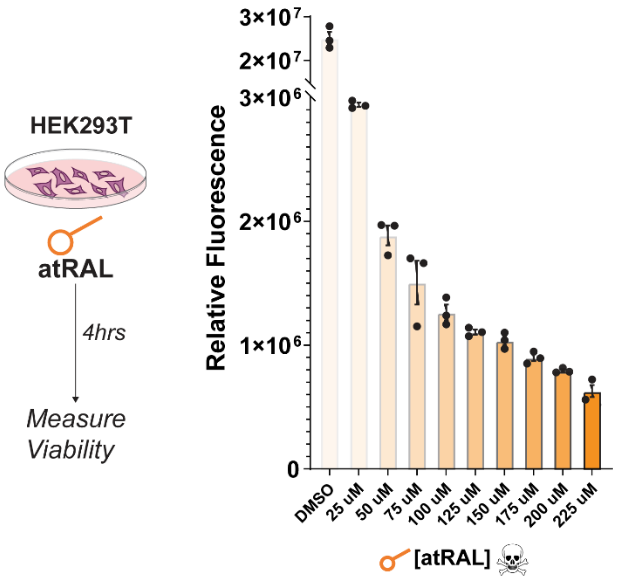


**Figure S2. Cytotoxicity of atRAL in HEK293T cells.** To assess the cyto toxic impact of exogenous retinal, HEK293T cells were transfected with an empty P1D4-hrGFPII vector and allowed to recover for 48 hours. Cells were subsequently treated with varying concentrations of atRAL (0–225µM)for 4 hours. Cell viability was quantified using the CellTiter-Blue (Promega) resazurin-based assay, with fluorescence measured on a BMG Labtech plate reader. Data are presented as Relative Fluorescence Units (RFU), where a dose-dependent decrease in fluorescence indicates a reduction in metabolic activity and cell viability corresponding to increasing atRAL concentrations. Individual data points represent technical replicates (n = 3) and error bars indicate ± SEM.

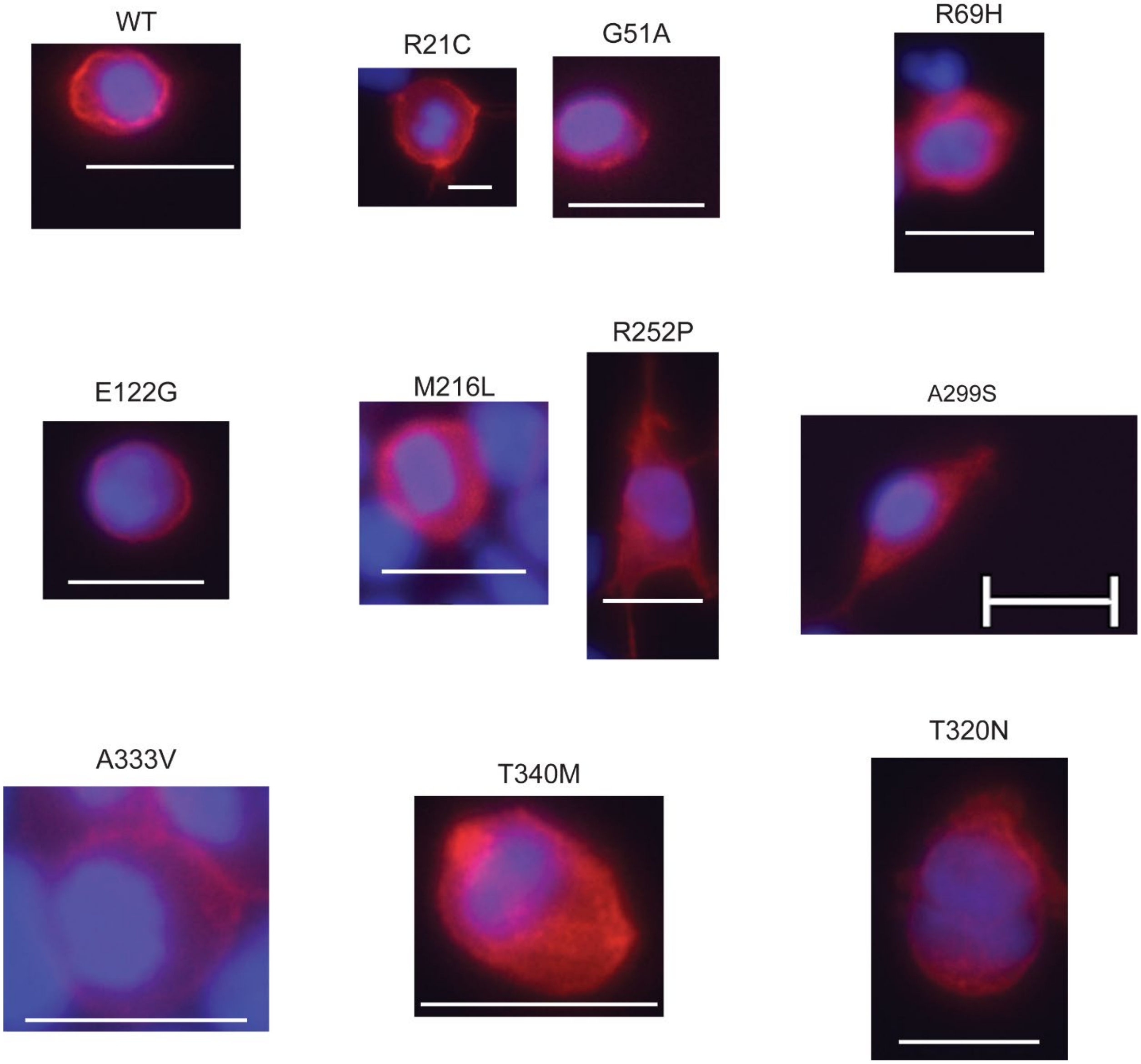


**Figure S3- Subcellular localization of Human RHO variants in HEK293T cells.** Immunocytochemistry was performed to evaluate the expression and trafficking of wild-type (WT) and mutant Human RHO. HEK293T cells were transiently transfected with the indicated RHO variants and imaged via fluorescence microscopy. RHO (red) was visualized using anti-1D4 antibody (Abcam) followed by Alexa-488 secondary and nuclei were counterstained with DAPI (blue). WT RHO and alleles exhibit characteristic plasma membrane localization.

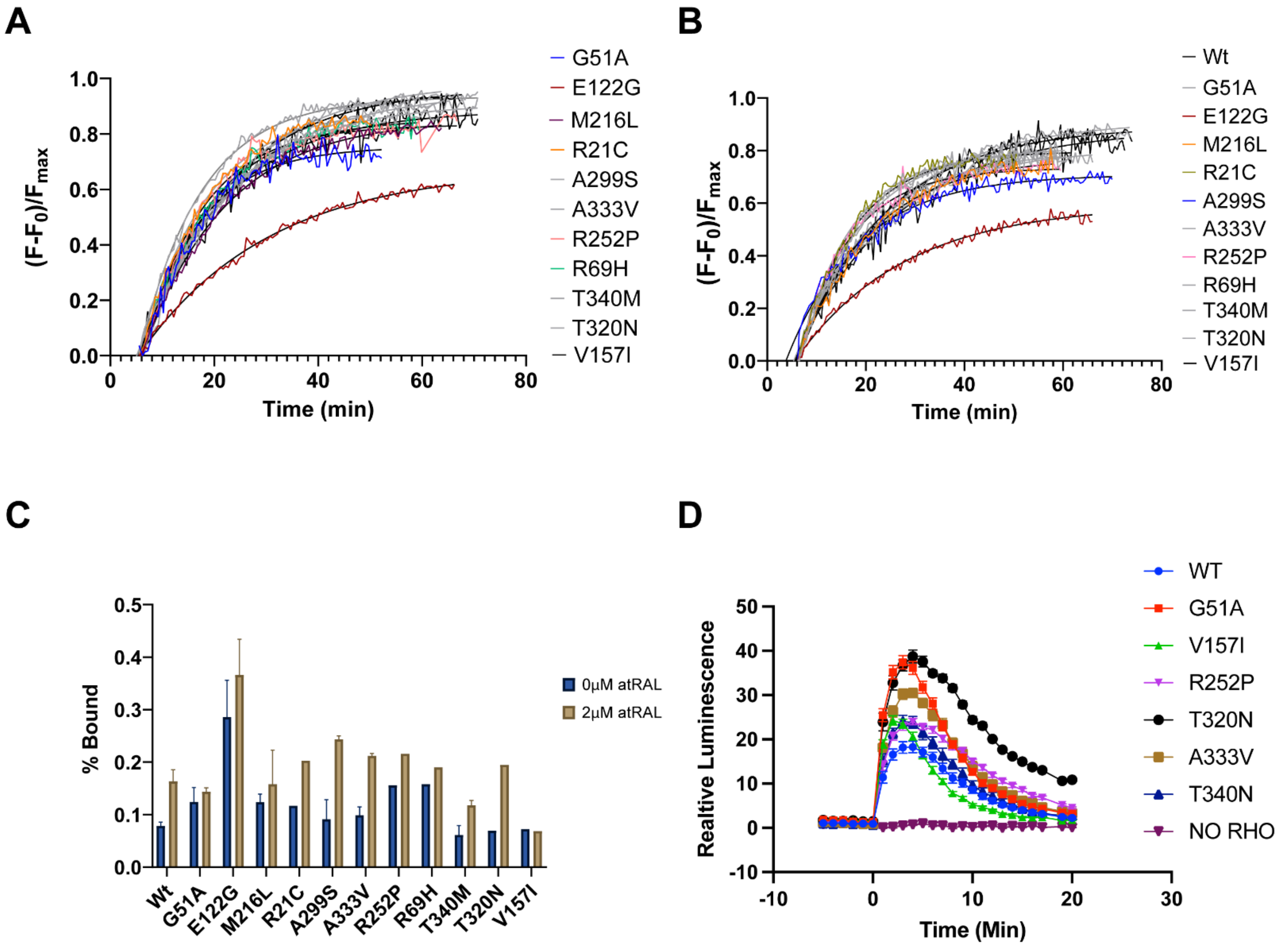


**Figure S4- Functional characterization of Human rhodopsin alleles of interest. (A-B)** Retinal release assay of human RHO alleles with 0 (A) or 2 µM (B) exogenous atRAL. Fluorescence data was baselined and normalized to max fluorescence to compare MII decay and $F_{bound}$ values. **(C)** Calculated fraction bound retinal at steady at both 0 and 2µM exogenous atRAL with error bars indicating SEM. **(D)** Glosensor assay monitoring signaling of RHO after light bleach at time 0. C and D are normalized to protein expression.

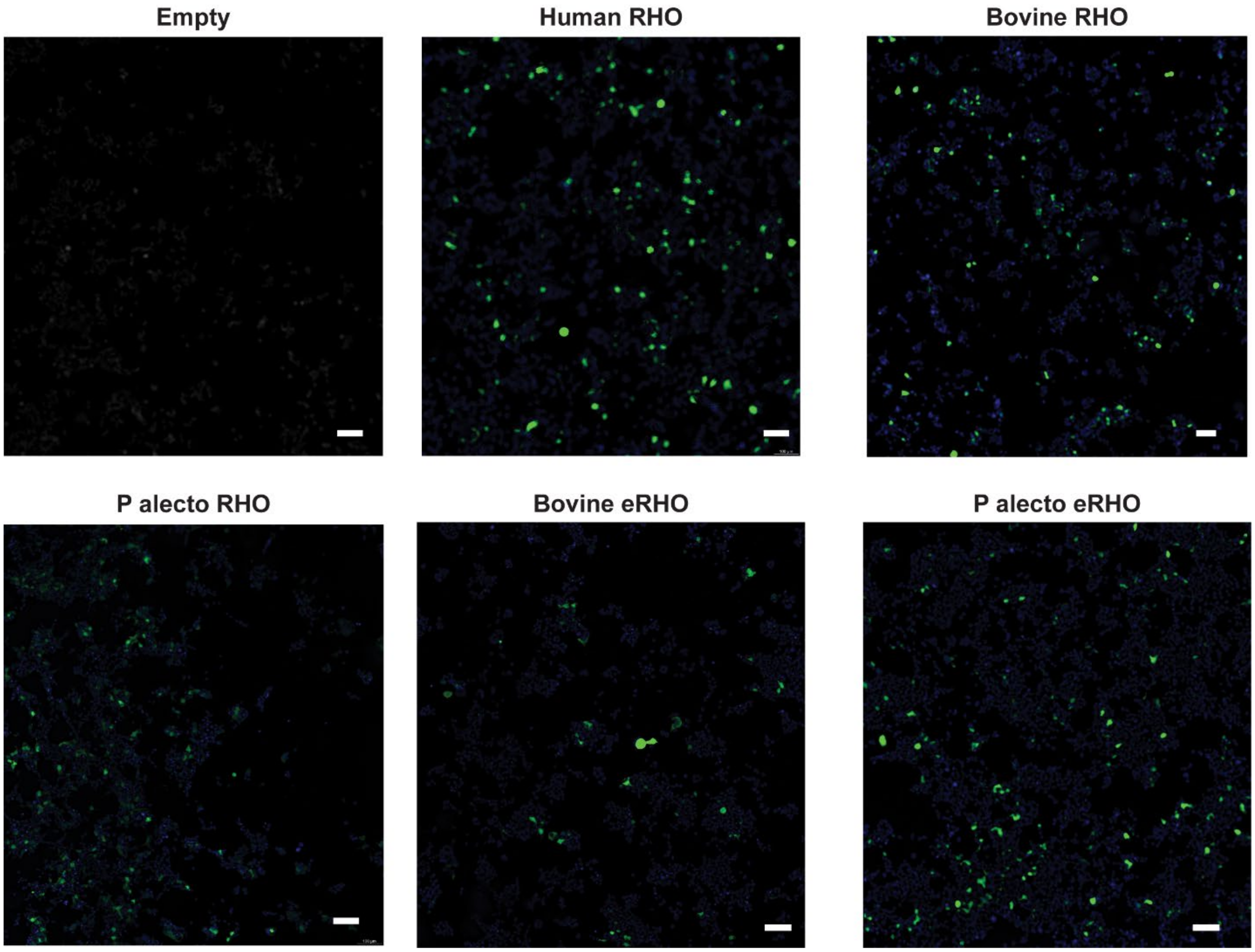


**Figure S5 – Immunocytochemistry of Rhodopsin in HEK293T cells**. Cells were transfected with empty vector, or WT RHO orthologs, or engineered RHOs. Cells were stained with anti-1D4 primary antibody, followed by Alexa-488 secondary, and imaged under widefield with computational clearing (Leica, MICA). Scale bar = 100 µm.

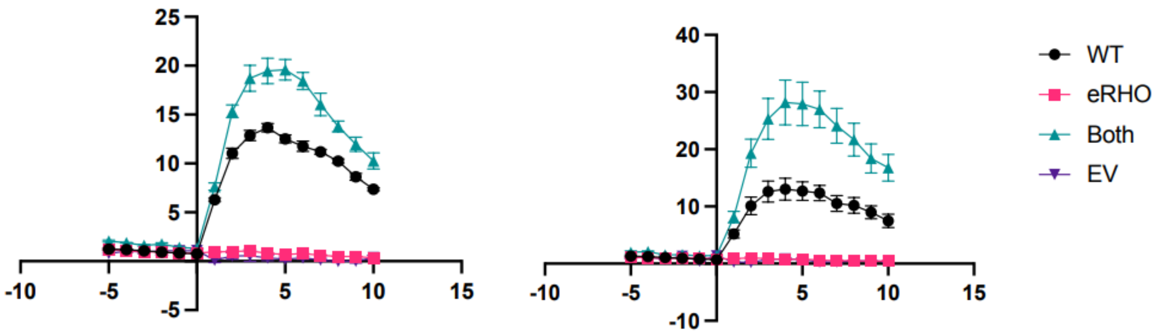


**Fig. S6. eRHO enhances WT RHO signalling.** Glosensor assay of HEK293T cells following various transfections. Left, 1 μM exogenous atRAL was added prior to light bleach at time 0. Right, 100 nM.. Y-axis is luminescence normalized to protein expression. X-axis denotes time in minutes.

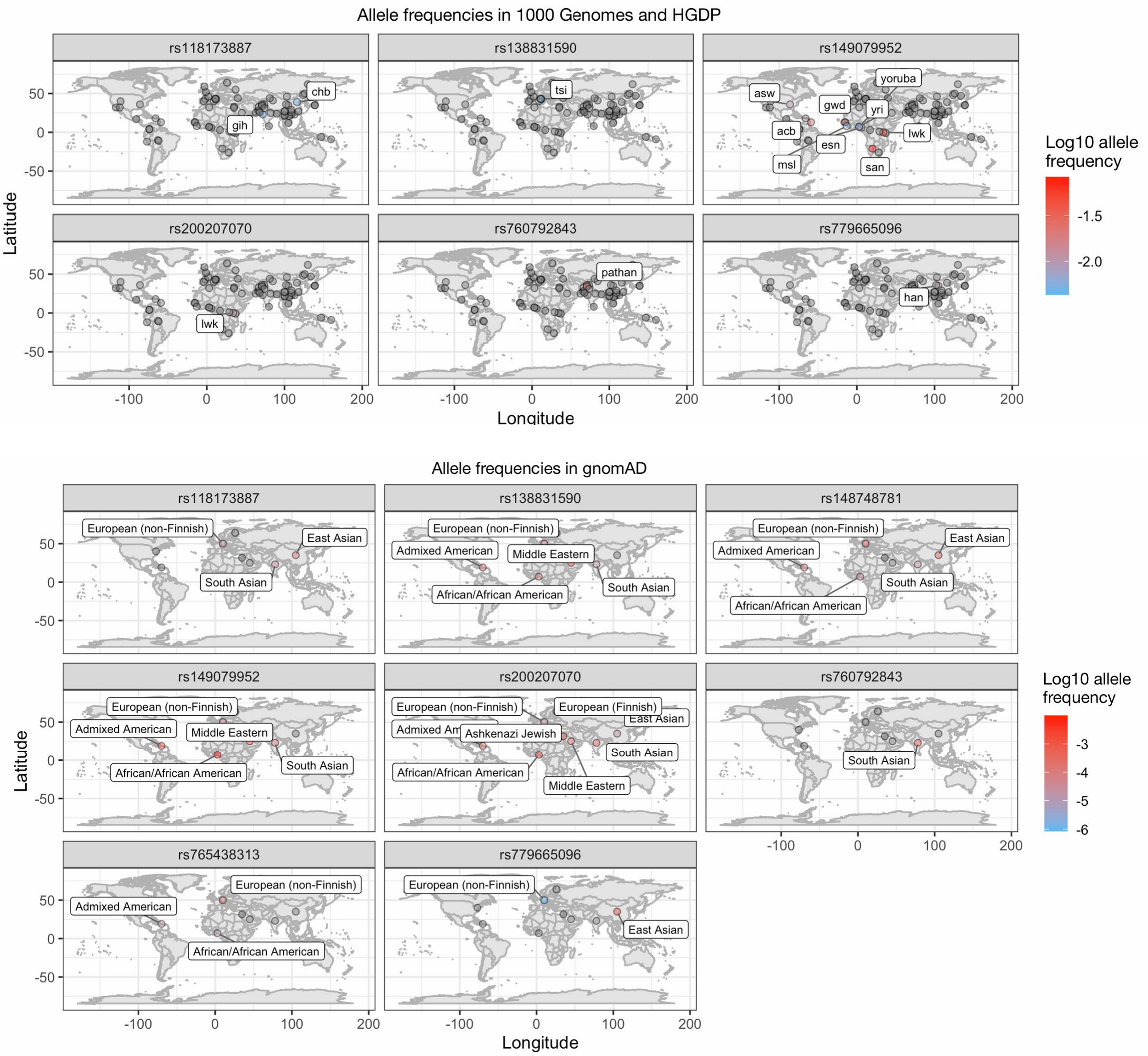


**Fig S7. Allele frequencies in 1KG + HGDP with gnomAD v4.1**. Not all variants that are in gnomAD appear in 1000 Genomes, hence different numbers of plots. In all plots, grey points indicate the allele is not present in that location, and only populations with a non-zero allele frequency are labelled.

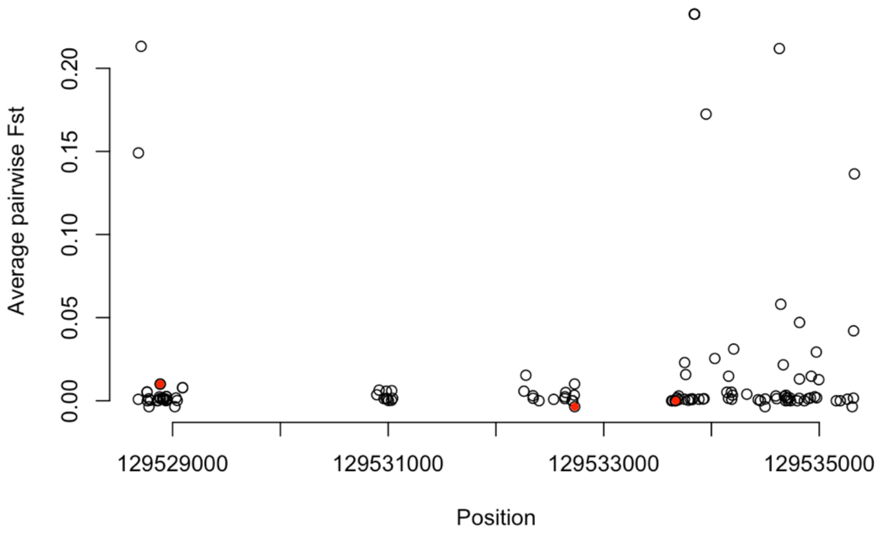


**Fig. S8. Pairwise Fst for all SNPs in RHO exons (n=114).** Variants functionally characterized in this study are red.

**Table S1**. Biochemical phenotypes of RHO from different species and different human RHO mutations. All values were determined at 20ºC with no exogenous atRAL.

| **Genotype** | **$\lambda max$ (nm)** | **Expression (Abs)** | **t*1/2* (min)** | | **Fraction bound** | | **Dissociation constant ($K_d$)** |
|---|---|---|---|---|---|---|---|
| | | | average | | average | SD | |
| *H. sapiens* | 493.8 | 0.012675 | 10.923 | 0.633 | 0.071455 | 0.010893 | 2.925 |
| *B. taurus* | 498.3 | 0.012633 | 10.893 | 0.285 | 0.1237 | 0.024732 | 1.72 |
| *P. alecto* | 500.5 | 0.010759 | 21.9 | 5.959 | 0.42853 | 0.063463 | 0.182 |
| G51A | 495.3 | 0.0812 | 13.27 | 0.4594 | 0.1242 | 0.06108 | 1.667 |
| E122G | 478.4 | 0.0337 | 16.29 | 2.49 | 0.2857 | 0.1219 | 0.482 |
| M216L | 494.3 | 0.091 | 14.6 | 2.079 | 0.1237 | 0.02667 | 1.676 |
| R21C | 493.8 | 0.038 | 8.559 | | 0.117 | | 16.893 |
| A299S | 495.8 | 0.1255 | 11.37 | 2.407 | 0.0911 | 0.07451 | 22.402 |
| A333V | 494.4 | 0.108 | 11.4 | 1.564 | 0.09873 | 0.0278 | 2.221 |
| R252P | 493.9 | 0.098 | 7.541 | | 0.1557 | | 1.236 |
| R69H | 493.1 | 0.0585 | 9.103 | | 0.1583 | | 1.208 |
| T340M | 494.6 | 0.133 | 10.85 | 0.6788 | 0.06104 | 0.02602 | 34.665 |
| T320N | 494 | 0.08 | 7.771 | | 0.06924 | | 30.263 |
| V157I | 495.8 | 0.076 | 11.24 | | 0.07239 | | 28.837 |

**Table S2**. Human RHO alleles information from Gnomad and allele age information

| | | Allele frequency information from gnomAD | | | | | | | |
|---|---|---|---|---|---|---|---|---|---|
| **Additional identifiers** | **Codon change** | **Total allele counts** | **Global allele frequency** | **African (AFR)** | **East Asian (EAS)** | **South Asian (SAS)** | **Admixed Am. (AMR)** | **European (NFE)** | **Allele age** ( from [https://human.genome.dating](https://human.genome.dating) ) |
| NM_000539.3:c.(152G>C) | Gly-Ala | 941 | 0.06% | 0.9873% *(AC: 741)* | 0% | 0.0121% *(AC: 11)* | 0.0766% *(AC: 46)* | 0.0055% *(AC: 65)* | 2,222.0 – 2,565.8 |
| NM_000539.3:c.(895G>T) | Ala-Ser | 80 | 0.01% | 0% | 0.1604% *(AC: 72)* | 0% | 0% | 0.0001% *(AC: 1)* | |
| NM_000539.3:c.(998C>T) | Ala-Val | 62 | 0.00% | 0% | 0% | 0.0637% *(AC: 58)* | 0% | 0.0003% *(AC: 4)* | |
| NM_000539.3:c.(959C>A) | Thr-Asn | 136 | 0.01% | 0.0080% *(AC: 6)* | 0% | 0.0022% *(AC: 2)* | 0.0183% *(AC: 11)* | 0.0077% *(AC: 91)* | |
| NM_000539.3:c.(206G>A) | Arg-His | 35 | 0.00% | 0% | 0.0133% *(AC: 6)* | 0.0033% *(AC: 3)* | 0% | 0.0021% *(AC: 25)* | 266.1 – 844.1 |
| NM_000539.3:c.(469G>A) | Val-Ile | 165 | 0.01% | 0.1025% *(AC: 77)* | 0.0022% *(AC: 1)* | 0.0395% *(AC: 36)* | 0.0199% *(AC: 12)* | 0.0017% *(AC: 21)* | 224.6 – 532.5 |
| NM_000539.3:c.(1019C>T) | Thr-Met | 358 | 0.02% | 0.0026% *(AC: 2)* | 0.0401% *(AC: 18)* | 0.0022% *(AC: 2)* | 0.0066% *(AC: 4)* | 0.0269% *(AC: 318)* | |
| NM_000539.3:c.(755G>C) | Arg-Pro | 142 | 0.01% | 0.0013% *(AC: 1)* | 0% | 0% | 0.0016% *(AC: 1)* | 0.0114% *(AC: 135)* | |

**Table S3**. Electronic Health Record phecodes for “Blindness and low vision”

| ICD | vocabulary_id | ICD_string |
|---|---|---|
| 369.03 | ICD9CM | Better eye: near-total vision impairment; lesser eye: total vision impairment |
| 369 | ICD9CM | Blindness of both eyes, impairment level not further specified |
| 369.02 | ICD9CM | Better eye: near-total vision impairment; lesser eye: not further specified |
| 369.9 | ICD9CM | Unspecified visual loss |
| 369.13 | ICD9CM | Better eye: severe vision impairment; lesser eye: near-total vision impairment |
| 369.01 | ICD9CM | Better eye: total vision impairment; lesser eye: total vision impairment |
| 369.04 | ICD9CM | Better eye: near-total vision impairment; lesser eye: near-total vision impairment |
| 369.2 | ICD9CM | Moderate or severe vision impairment, both eyes |
| 369.18 | ICD9CM | Better eye: moderate vision impairment; lesser eye: profound vision impairment |
| 369.17 | ICD9CM | Better eye: moderate vision impairment; lesser eye: near-total vision impairment |
| 369.16 | ICD9CM | Better eye: moderate vision impairment; lesser eye: total vision impairment |
| 369.14 | ICD9CM | Better eye: severe vision impairment; lesser eye: profound vision impairment |
| 369.12 | ICD9CM | Better eye: severe vision impairment; lesser eye: total vision impairment |
| 369.11 | ICD9CM | Better eye: severe vision impairment; lesser eye: blind, not further specified |
| 369.1 | ICD9CM | Blindness one eye; low vision other eye |
| 369.1 | ICD9CM | Moderate or severe vision impairment, better eye; profound vision impairment of lesser eye |
| 369.08 | ICD9CM | Better eye: profound vision impairment; lesser eye: profound vision impairment |
| 369.07 | ICD9CM | Better eye: profound vision impairment; lesser eye: near-total vision impairment |
| 369.06 | ICD9CM | Better eye: profound vision impairment; lesser eye: total vision impairment |
| 369.05 | ICD9CM | Better eye: profound vision impairment; lesser eye: not further specified |
| 369.22 | ICD9CM | Better eye: severe vision impairment; lesser eye: severe vision impairment |
| 369.15 | ICD9CM | Better eye: moderate vision impairment; lesser eye: blind, not further specified |
| 369.2 | ICD9CM | Moderate or severe vision impairment, both eyes, impairment level not further specified |
| 369 | ICD9CM | Profound vision impairment, both eyes |
| 369.8 | ICD9CM | Unqualified visual loss, one eye |
| 369.6 | ICD9CM | Blindness, one eye |
| 369.61 | ICD9CM | One eye: total vision impairment; other eye: not specified |
| 369.62 | ICD9CM | One eye: total vision impairment; other eye: near-normal vision |
| 369.63 | ICD9CM | One eye: total vision impairment; other eye: normal vision |
| 369.64 | ICD9CM | One eye: near-total vision impairment; other eye: vision not specified |
| 369.65 | ICD9CM | One eye: near-total vision impairment; other eye: near-normal vision |
| 369.66 | ICD9CM | One eye: near-total vision impairment; other eye: normal vision |

| | | |
|---|---|---|
| 369.67 | ICD9CM | One eye: profound vision impairment; other eye: vision not specified |
| 369.68 | ICD9CM | One eye: profound vision impairment; other eye: near-normal vision |
| 369.69 | ICD9CM | One eye: profound vision impairment; other eye: normal vision |
| 369.7 | ICD9CM | Moderate or severe vision impairment, one eye |
| 369.7 | ICD9CM | Low vision, one eye |
| 369.71 | ICD9CM | One eye: severe vision impairment; other eye: vision not specified |
| 369.72 | ICD9CM | One eye: severe vision impairment; other eye: near-normal vision |
| 369.74 | ICD9CM | One eye: moderate vision impairment; other eye: vision not specified |
| 369.75 | ICD9CM | One eye: moderate vision impairment; other eye: near-normal vision |
| 369.76 | ICD9CM | One eye: moderate vision impairment; other eye: normal vision |
| 369.73 | ICD9CM | One eye: severe vision impairment; other eye: normal vision |
| 369.24 | ICD9CM | Better eye: moderate vision impairment; lesser eye: severe vision impairment |
| 369.4 | ICD9CM | Legal blindness, as defined in U.S.A. |
| 369.6 | ICD9CM | Profound vision impairment, one eye |
| 369.23 | ICD9CM | Better eye: moderate vision impairment; lesser eye: impairment not further specified |
| 369 | ICD9CM | Blindness and low vision |
| 369.3 | ICD9CM | Unqualified visual loss, both eyes |
| 369.25 | ICD9CM | Better eye: moderate vision impairment; lesser eye: moderate vision impairment |
| 369.21 | ICD9CM | Better eye: severe vision impairment; lesser eye; impairment not further specified |
| H54.122 | ICD10CM | Low vision right eye category 2, blindness left eye |
| H54.1215 | ICD10CM | Low vision right eye category 1, blindness left eye category 5 |
| H54.1214 | ICD10CM | Low vision right eye category 1, blindness left eye category 4 |
| H54.1213 | ICD10CM | Low vision right eye category 1, blindness left eye category 3 |
| H54.121 | ICD10CM | Low vision right eye category 1, blindness left eye |
| H54.12 | ICD10CM | Blindness, left eye, low vision right eye |
| H54.1152 | ICD10CM | Blindness right eye category 5, low vision left eye category 2 |
| H54.1151 | ICD10CM | Blindness right eye category 5, low vision left eye category 1 |
| H54.115 | ICD10CM | Blindness right eye category 5, low vision left eye |
| H54.1223 | ICD10CM | Low vision right eye category 2, blindness left eye category 3 |
| H54.1224 | ICD10CM | Low vision right eye category 2, blindness left eye category 4 |
| H54.1225 | ICD10CM | Low vision right eye category 2, blindness left eye category 5 |
| H54.3 | ICD10CM | Unqualified visual loss, both eyes |
| H54.2X22 | ICD10CM | Low vision right eye category 2, low vision left eye category 2 |
| H54.2X21 | ICD10CM | Low vision right eye category 2, low vision left eye category 1 |
| H54.2X2 | ICD10CM | Low vision, right eye, category 2 |
| H54.2X12 | ICD10CM | Low vision right eye category 1, low vision left eye category 2 |
| H54.2X11 | ICD10CM | Low vision right eye category 1, low vision left eye category 1 |

| | | |
|---|---|---|
| H54.2X1 | ICD10CM | Low vision, right eye, category 1 |
| H54.2X | ICD10CM | Low vision, both eyes, different category levels |
| H54.2 | ICD10CM | Low vision, both eyes |
| H54.1142 | ICD10CM | Blindness right eye category 4, low vision left eye category 2 |
| H54.1141 | ICD10CM | Blindness right eye category 4, low vision left eye category 1 |
| H54.114 | ICD10CM | Blindness right eye category 4, low vision left eye |
| H54.0X43 | ICD10CM | Blindness right eye category 4, blindness left eye category 3 |
| H54.0X44 | ICD10CM | Blindness right eye category 4, blindness left eye category 4 |
| H54.0X45 | ICD10CM | Blindness right eye category 4, blindness left eye category 5 |
| H54.0X5 | ICD10CM | Blindness right eye, category 5 |
| H54.0X53 | ICD10CM | Blindness right eye category 5, blindness left eye category 3 |
| H54.0X54 | ICD10CM | Blindness right eye category 5, blindness left eye category 4 |
| H54.0X55 | ICD10CM | Blindness right eye category 5, blindness left eye category 5 |
| H54.1 | ICD10CM | Blindness, one eye, low vision other eye |
| H54.10 | ICD10CM | Blindness, one eye, low vision other eye, unspecified eyes |
| H54.0X4 | ICD10CM | Blindness right eye, category 4 |
| H54.0X35 | ICD10CM | Blindness right eye category 3, blindness left eye category 5 |
| H54.1132 | ICD10CM | Blindness right eye category 3, low vision left eye category 2 |
| H54.1131 | ICD10CM | Blindness right eye category 3, low vision left eye category 1 |
| H54.113 | ICD10CM | Blindness right eye category 3, low vision left eye |
| H54 | ICD10CM | Blindness and low vision |
| H54.0 | ICD10CM | Blindness, both eyes |
| H54.0X | ICD10CM | Blindness, both eyes, different category levels |
| H54.0X3 | ICD10CM | Blindness right eye, category 3 |
| H54.0X33 | ICD10CM | Blindness right eye category 3, blindness left eye category 3 |
| H54.0X34 | ICD10CM | Blindness right eye category 3, blindness left eye category 4 |
| H54.11 | ICD10CM | Blindness, right eye, low vision left eye |
| H54.8 | ICD10CM | Legal blindness, as defined in USA |
| H54.51 | ICD10CM | Low vision, right eye, normal vision left eye |
| H54.52A | ICD10CM | Low vision, left eye, category 1-2 |
| H54.511 | ICD10CM | Low vision, right eye, category 1-2 |
| H54.511A | ICD10CM | Low vision right eye category 1, normal vision left eye |
| H54.512A | ICD10CM | Low vision right eye category 2, normal vision left eye |
| H54.52 | ICD10CM | Low vision, left eye, normal vision right eye |
| H54.61 | ICD10CM | Unqualified visual loss, right eye, normal vision left eye |
| H54.52A1 | ICD10CM | Low vision left eye category 1, normal vision right eye |
| H54.52A2 | ICD10CM | Low vision left eye category 2, normal vision right eye |
| H54.6 | ICD10CM | Unqualified visual loss, one eye |
| H54.60 | ICD10CM | Unqualified visual loss, one eye, unspecified |

| | | |
|---|---|---|
| H54.62 | ICD10CM | Unqualified visual loss, left eye, normal vision right eye |
| H54.7 | ICD10CM | Unspecified visual loss |
| H54.50 | ICD10CM | Low vision, one eye, unspecified eye |
| H54.5 | ICD10CM | Low vision, one eye |
| H54.4 | ICD10CM | Blindness, one eye |
| H54.40 | ICD10CM | Blindness, one eye, unspecified eye |
| H54.41 | ICD10CM | Blindness, right eye, normal vision left eye |
| H54.413 | ICD10CM | Blindness, right eye, category 3 |
| H54.413A | ICD10CM | Blindness right eye category 3, normal vision left eye |
| H54.414 | ICD10CM | Blindness, right eye, category 4 |
| H54.414A | ICD10CM | Blindness right eye category 4, normal vision left eye |
| H54.415 | ICD10CM | Blindness, right eye, category 5 |
| H54.42A5 | ICD10CM | Blindness left eye category 5, normal vision right eye |
| H54.42A4 | ICD10CM | Blindness left eye category 4, normal vision right eye |
| H54.42A3 | ICD10CM | Blindness left eye category 3, normal vision right eye |
| H54.42A | ICD10CM | Blindness, left eye, category 3-5 |
| H54.42 | ICD10CM | Blindness, left eye, normal vision right eye |
| H54.415A | ICD10CM | Blindness right eye category 5, normal vision left eye |
